# Geometric constraints on the architecture of mammalian cortical connectomes

**DOI:** 10.1101/2025.09.17.676944

**Authors:** Francis Normand, Mehul Gajwani, Trang Cao, Jace Cruddas, Arshiya Sangchooli, Stuart Oldham, Alexander Holmes, Peter A. Robinson, James C. Pang, Alex Fornito

## Abstract

The intricate network of axonal fibres that forms the mammalian cortical connectome has a complex topology, being organized in a way that is neither completely regular nor random, as well as a characteristic topography, in which specific anatomical locations are imbued with distinctive connectivity profiles. The mechanisms that give rise to such properties remain a mystery. Here, we formulate a simple analytic model derived from neural field theory that prioritizes physical constraints on connectome architecture by assuming that connectivity is preferentially concentrated between pairs of cortical locations that facilitate the excitation of resonant geometric modes of the cortex. We show that the model outperforms existing approaches in reproducing multiple topological and topographical properties of cortical connectomes mapped at spatial scales spanning orders of magnitude in humans, chimpanzees, macaques, marmosets, and mice, as mapped with either non-invasive diffusion magnetic resonance imaging or invasive viral tract-tracing. Our findings thus point to a fundamental role of geometry in shaping the multiscale architecture of cortical connectomes that has been conserved across 90 million years of evolution.

---

The >16 billion neurons of the human cortex are linked by ∼150,000 km of axons making approximately 164 trillion synapses (*1*). This intricate web of connectivity, called the cortical connectome, shows a complex organization that is neither completely random nor regular (*2–4*), and which provides an anatomical substrate for the coordinated dynamics that underlie our thoughts, emotions, and behaviour (*2–4*). The specific mechanisms that shape the complex architecture of the cortical connectomes of human and other mammals remain a mystery.

Mathematical models can be used to understand such mechanisms by specifying rules to grow synthetic networks that reproduce key features of empirical connectomes (*5–8*). These features are commonly quantified in relation to network topology and topography. Network topology refers to how pairs of neural elements are connected to each other, regardless of their physical locations. Network topography refers to the specific way in which neural elements are interconnected in physical space and thus corresponds to the spatial embedding of the network (*4, 9*). Numerous studies have shown that the connectomes of human and non-human species are not only topologically complex (*3, 4*), but that they also have a characteristic spatial topography; for instance, highly connected regions in primates, known as hubs, are commonly localized within transmodal association cortices (*10–13*).

Extant models treat the cortex as a graph of discrete nodes linked by edges. The nodes represent neurons, neuronal populations, or macroscopic brain regions (depending on the spatial scale of interest), while the edges represent axonal connections between nodes (*3, 4*). When the models are defined in this way, edges established by a stochastic process dependent only on the distance between nodes is sufficient to capture numerous topological properties of empirical connectomes (*5,6,14–17*). This distance-dependence typically conforms to an exponential distance rule (EDR), in which the probability of connecting any two nodes declines as an approximately exponential function of their physical separation. Empirically, this EDR has been found to dominate the organization of connectomes mapped at different resolutions and in different species, including the microscale connectome of *Caenorhabditis elegans* (*7*), mesoscale connectomes of mouse (*18, 19*), marmoset (*20*), and macaque (*17,21,22*), and macroscale connectome of humans (*14,23*). The EDR has therefore been proposed as a universal principle of connectome organization (*17*) that is thought to arise from a biological pressure to minimize the metabolic costs of network wiring (*9, 24–29*).

Other work indicates that models combining an EDR with a homophilic “like attracts like” attachment rule that favours connections between nodes with similar connectivity profiles can better capture the connectome’s topology than an EDR alone (*5,6,8*). This form of homophilic attachment may result from Hebbian activity-dependent plasticity mechanisms (*30*), selection pressures to maintain economic trade-offs between wiring cost and adaptive function (*9, 27, 28*), or a more general process in which areas with similar cellular and molecular properties are more likely to connect with each other because they share concurrent maturational trajectories (*6, 7, 31–34*).

Despite these advances, graph-based models used thus far are limited in three key respects (for other limitations, see Ref. (*35*)). First, they depend on how the network is discretized (i.e., how cortical nodes are defined), which can overlook physical constraints on dynamics (*36*) and restricts the model to the spatial scale of the discretization, thus neglecting the cortex’s intrinsically multiscale architecture (*13, 37*). Second, most such models only capture the topology of binarized empirical connectomes (i.e., they only model the presence/absence of a connection) and ignore their weighted architecture (i.e., variations in the strength of different connections, which often span several orders of magnitude; for a recent exception, see (*38*)). Third, despite their success in reproducing the topology of connectomes, current models do not capture topographical properties, such as the spatial locations of network hubs (*6, 7*). This is a critical failure, given that diverse histological, molecular, and functional properties are spatially embedded in highly specific ways that are central to cortical organization and function (*13, 39–44*).

Here, we propose a different approach that draws inspiration from empirical observations that mature connectomes are shaped by an interplay between molecular gradients early in development that guide axons to their targets (*45, 46*) and the activity-dependent processes that determine which connections are strengthened and which are eliminated after an initial period of overgrowth (*47–57*). Rather than attempting to capture the intricacies of these complicated dynamical processes, we instead derive a simple model that approximates the net effect of such physically constrained processes on brain connectivity, as viewed through the lens of neural field theory (NFT). NFT refers to a class of mathematical models that treat large-scale neuronal dynamics (i.e., on spatial scales >0.5 mm) as waves of activity traveling on a spatially continuous cortical sheet governed by averaged cellular processes related to firing rates, soma voltage, and synaptic inputs. The approach has been used to explain diverse physiological phenomena observed with local field potentials, electroencephalography, and functional magnetic resonance imaging (*58–67*).

Under one extensively validated variant of NFT, spatiotemporal patterns of brain activity can be modelled as excitations of the natural modes of cortical geometry (*68*), known as *geometric eigenmodes*. These eigenmodes or modes are the preferred spatial resonances or standing waves of dynamics, with each mode defined by a specific spatial frequency (*68, 69*). The general preference of diverse physical and biological systems, including the brain, to express resonant patterns (*70–77*) suggests that the activity-dependent processes shaping connectome development will favour retention and augmentation of connections that facilitate their excitation. As a corollary, we should expect that axonal connections will preferentially link pairs of cortical locations that satisfy two criteria: (1) the locations reside close to antinodes of the geometric eigenmodes, which correspond to loci of maximal amplitude fluctuations in standing wave dynamics (*78, 79*); (2) their amplitudes on any given eigenmode have the same sign, meaning that the two locations are spatially in-phase during the oscillation. Connections between pairs of locations satisfying these criteria can act as communication channels, facilitating the spread of a focal input such that the energetic threshold to excite an eigenmode is lowered (*78, 79*) (Fig. 1b; for further details, see the Theory section below). These considerations lead to the hypothesis that pairs of cortical locations satisfying the above two criteria across many different eigenmodes will be preferentially connected. We test this hypothesis by developing a simple model that can be directly derived from the formalism of NFT. We show that this model can generate connectomes that successfully capture the binary and weighted characteristics of both the topology *and* topography of empirical connectomes measured via non-invasive diffusion magnetic resonance imaging (MRI) or invasive tract-tracing in humans, chimpanzees, macaques, marmosets, and mice, and at spatial resolutions spanning orders of magnitude. The generalizability of our model points to a fundamental and universal role of cortical geometry in sculpting the architecture of cortical connectomes.

**Figure 1:**
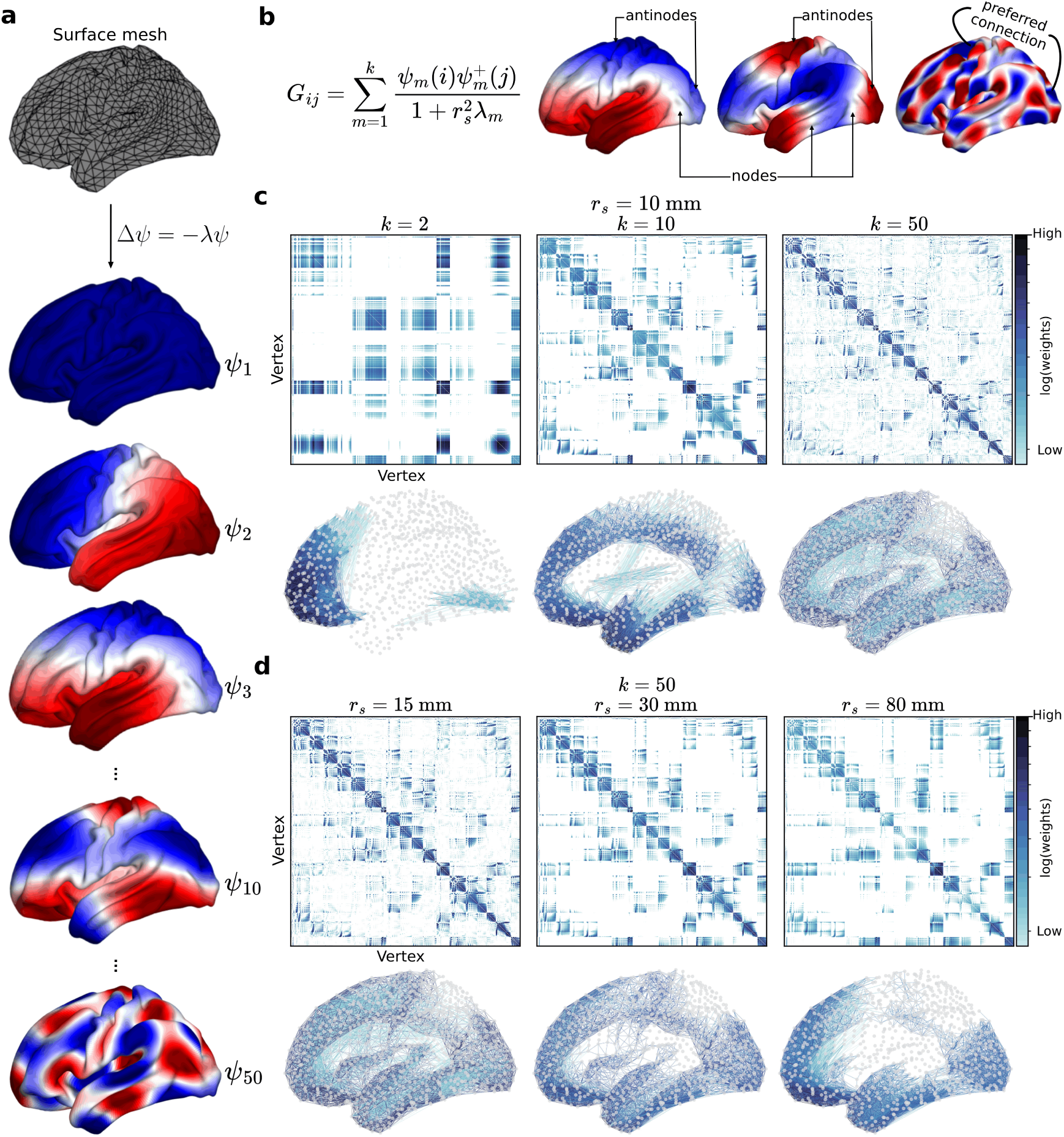
A Geometric Eigenmode Model (GEM) of the connectome. a: Human geometric eigenmodes of the Laplace-Beltrami Operator, Δ, which describes the geometry of a 2Dtriangular mesh representation of the neocortical surface. An example surface mesh comprising 4,386 vertices is shown at the top, with example eigenmodes below. The modes, 𝜓, which correspond to preferred, resonant spatial patterns of cortical excitation, are ordered from long to short spatial wavelengths (top to bottom). The wavelengths are inversely proportional to the square root of the eigenvalues, 𝜆. Red and blue colours differentiate regions that oscillate with opposite amplitudes, thus defining a standing wave of dynamics (note that the sign/colour is arbitrary). b: Mathematical expression describing the GEM (left) and a schematic demonstrating the locations of nodes and antinodes on three eigenmodes (right). The GEM favours connections between pairs of cortical locations, *i* and *j*, at locations close to antinodes with the same sign in each *m*th mode, 𝜓_*m*_, summed across the first 𝑘 modes. 𝜓_*m*_^+^ denotes the pseudoinverse of 𝜓_*m*_ and the normalization term favours contributions from long-wavelength modes (i.e., modes with small 𝜆_*m*_), with *r*_*s*_ corresponding to the length scale of wave propagation (see Methods for derivation). The free parameters 𝑘 and *r*_*s*_ are fitted to optimize the similarity between model and empirical connectomes. c: Example vertex-resolution connectomes constructed using different numbers of modes, 𝑘, for fixed *r*_*s*_ = 10 mm. The examples from left to right show the results for 𝑘 = *{*2, 10, 50*}*. d: Example vertex-resolution connectomes constructed using different length scales, *r*_*s*_, for fixed 𝑘 = 50. The examples from left to right show the results for *r*_*s*_ = *{*15, 30, 80*}* mm. For panels c and d, the top plot shows the connectivity matrix with low to high weights in logarithmic scale coloured from light blue to black, while the bottom plot shows the corresponding graph-based representation with circles denoting neocortical surface mesh vertices and lines denoting edges coloured according to weight.

## Theory

Within a broad class of NFTs, cortical dynamics can be modelled using a spatial propagator, or Green’s function, which determines the spatiotemporal response of a system (i.e., the cortex) to an impulse stimulus (*65, 80, 81*). Hence, the total cortical synaptic response 𝜙(**r**, 𝑡) at spatial location **r** and time 𝑡, due to incoming pulses from all other locations **r**^′^ at time 𝑡^′^ can be described by the general propagator equation

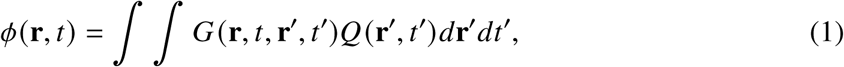

where 𝑄(**r**^′^, 𝑡^′^) represents a spike of activity at (**r**^′^, 𝑡^′^) and 𝐺(**r**, 𝑡, **r**^′^, 𝑡^′^) represents the mean propagator for axons projecting from (**r**^′^, 𝑡^′^) to (**r**, 𝑡), thus defining the connectivity kernel of the system (*64*). Note that in our analysis, each cortical location **r** corresponds to one of 4,386 vertices of a 2D triangular mesh model of the human cortical midthickness surface defined using established procedures for processing T1-weighted MRI (*82*).

Different forms of the propagator could be derived depending on the assumption of the underlying physical model that underpins Equation 1. Here, we derive the propagator from a well-validated form of NFT (*65*), which assumes that the spatiotemporal response, 𝜙(**r**, 𝑡), in cortical location **r** at time 𝑡 due to a stimulus can be approximated by a damped wave equation,

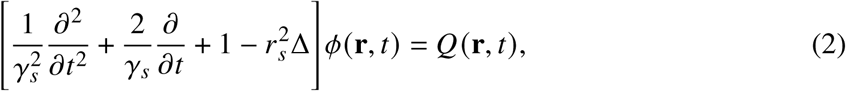

where Δ is the Laplace-Beltrami operator (LBO) that captures the geometry of the 2D neocortical surface manifold (see Methods for more details), 𝛾_*s*_ = 𝑣/*r*_*s*_ is the damping rate of the wave, 𝑣 is the characteristic axonal propagation velocity, and *r*_*s*_ is the characteristic range of the axons that determines the spatial length scale of wave propagation. Note that the spatial component of the dynamics in Equation 2 is determined purely by the cortical geometry, as encoded by the LBO, which can be shown to prioritize EDR-like distance-dependent interactions (*65, 69, 80, 83*). Accordingly, the spatial modes of the dynamics—also called standing waves—correspond to the resonant modes of the geometry of the cortex and can be obtained by solving the eigenvalue problem for the LBO, which is also known as the Helmholtz equation, Δ𝜓 = −𝜆𝜓 (see Fig. 1a and Methods).

The resulting eigenvalues are in increasing order (0 = 𝜆_1_ ≤ 𝜆_2_ ≤ … ≤ 𝜆_*m*_) such that the geometric eigenmodes (or simply modes), 𝜓, are ordered from long to short characteristic spatial wavelengths (wavelength ≈ 2𝜋/√𝜆) (Fig. 1a). Each eigenmode thus corresponds to a preferred spatial pattern of cortical excitation, each with a characteristic spatial frequency and topographical layout of nodes and antinodes (Fig. 1b). Nodes correspond to locations with zero amplitude that do not oscillate over time, while antinodes show maximal amplitude fluctuations, oscillating between high and low values. The resonant excitations of a given eigenmode can be triggered most effectively by stimulating one or more antinodes at a preferred oscillation frequency of the eigenmode, with the magnitude of the eigenvalue, 𝜆, being related to the energy required to excite the mode.

These considerations suggest that the excitation of a given mode will be facilitated when there are direct connections between locations close to antinodes, since the connections effectively act as communication channels through which the excitation of one antinode can be efficiently transmitted to another, thereby reducing the energy requirements for eigenmode excitation (*78, 79*) (Fig. 1b). Following this intuition, we formulate a Geometric Eigenmode Model (GEM) that models the connectome as a low-rank approximation of the time-independent (i.e., in the zero-frequency limit) connectivity propagator or Green’s function, **G**, of Equation 2 in terms of the geometric eigenmodes,

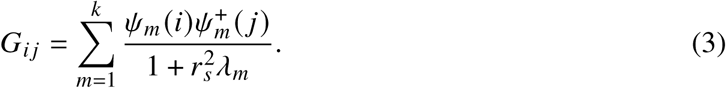

The term 𝐺_*i* *j*_ is an entry in the matrix **G** that corresponds to the strength of connection between two cortical locations *i* and *j*, and is defined as the products of the amplitudes 𝜓_*m*_ (*i*) and 𝜓_*m*_ (*j*) summed across the first 𝑘 modes. The denominator weights the sum by a decreasing function of the eigenvalues and the spatial length scale of the wave propagation *r*_*s*_ (Fig. 1b). For the cortex, *i* and *j* represent individual surface mesh vertices, referencing the spatial coordinates **r** and **r**^′^ introduced in Equation 1 and 𝜓_*m*_^+^ denotes the pseudoinverse of 𝜓_*m*_. See Methods for the full derivation of Equation 3.

Given a basis set of geometric eigenmodes, the GEM is completely deterministic with only two free parameters, 𝑘 and *r*_*s*_. The parameter 𝑘 controls how many modes contribute to the connectome model. The parameter *r*_*s*_ is the length scale in mm that controls the contribution of each mode, such that higher values increase the contribution of long-wavelength modes relative to shorter-wavelengths (see Fig. S1a). For any given parameter combination, the GEM produces a high-resolution weighted connectome mapped at the vertex level (see Figs. 1b, c, and d) in which pairs of cortical locations that exhibit large amplitude fluctuations across multiple modes, especially long-wavelength ones, are assigned stronger connection weights. Having a model that has intrinsically high resolution allows us to define connectomes at any coarser resolution, matching current widespread approaches for empirical connectome mapping that rely on atlas-based cortical parcellations to define region-scale network nodes (*4*).

## The GEM captures high-resolution topological and topographical properties of the human connectome

The simplicity of the GEM allows us to define connectomes at arbitrarily high resolutions, thus breaking any dependence on a specific parcellation atlas. As such, we first evaluate the performance of the model in capturing fine-grained details of human connectomes mapped at vertex resolution. To this end, we fit the GEM to an empirical group-average cortico-cortical connectome of the left hemisphere mapped using diffusion MRI acquired in 339 healthy adults and defined at the level of 4,386 neocortical surface mesh vertices (excluding the medial wall) derived from a group-average template surface (*82*) (see Methods).

We evaluate the performance of the GEM relative to four benchmark null models. The first benchmark, termed ‘LBO’, models connections based on a low-rank approximation of the LBO of the cortical mesh and accounts for the effects of geometry and the spatial topography of the eigenmodes. However, it uses a different weighting of eigenmode contributions compared to the weighting derived from the static limit of NFT-based wave dynamics, as in Equation 2, which is represented by the denominator of Equation 3 (see Methods). The second benchmark, termed ‘Permuted’, models connections using Equation 3, but after permuting the eigenvalues, thus testing the importance of their ordering in the normalization term derived from NFT. The third benchmark, termed ‘EDR-vertex’, models connections at random with probabilities following an exponential distance-dependent function to test the specific contribution of the geometric and NFT-derived dynamic processes implied by our model relative to a generic EDR connectivity kernel. The final benchmark, termed ‘Random’, models connections at random with edge weights following an Erdoős–Rényi-like process while preserving the density of the empirical connectomes. Note that the ‘LBO’, ‘Permuted’, ‘EDR-vertex’, and ‘Random’ models have 1, 0, 2, and 0 free parameters, respectively (see Methods for details of each null model, including their free parameters and optimization procedures).

We optimize all models using the same objective function, which is designed to capture critical elements of network topology and topography. Specifically, the objective function is given by the sum of rank correlations, 𝜌, between the following model and empirical properties: (1) edge weight sequences, based on the weights of the union of model and empirical edges; (2) binary node degree sequences; and (3) rank-based nodal strength sequences (see Methods for detailed explanation and justification).

For the GEM, a brute-force search over a parameter landscape spanning 1 mm ≤ *r*_*s*_ ≤ 20 mm and 1 ≤ 𝑘 ≤ 200 identifies optimal parameters *r*_*s*_ = 9.53 mm and 𝑘 = 108 modes (see Fig. S2a for the optimization landscape), indicating that only about 2.5% of all 4,386 available modes are required to account for the data. With this parameter combination, the model captures the rank ordering of the empirical connection weights (i.e., the edge weights correlation over the intersection of model and empirical edges is 𝜌 = 0.81; Fig. 2b) as well as the spatial topography of the network, determined using either binary or weighted degree maps, with spatial correlations of 𝜌 = 0.52 and 𝜌 = 0.47, respectively (Fig. 2c).

**Figure 2:**
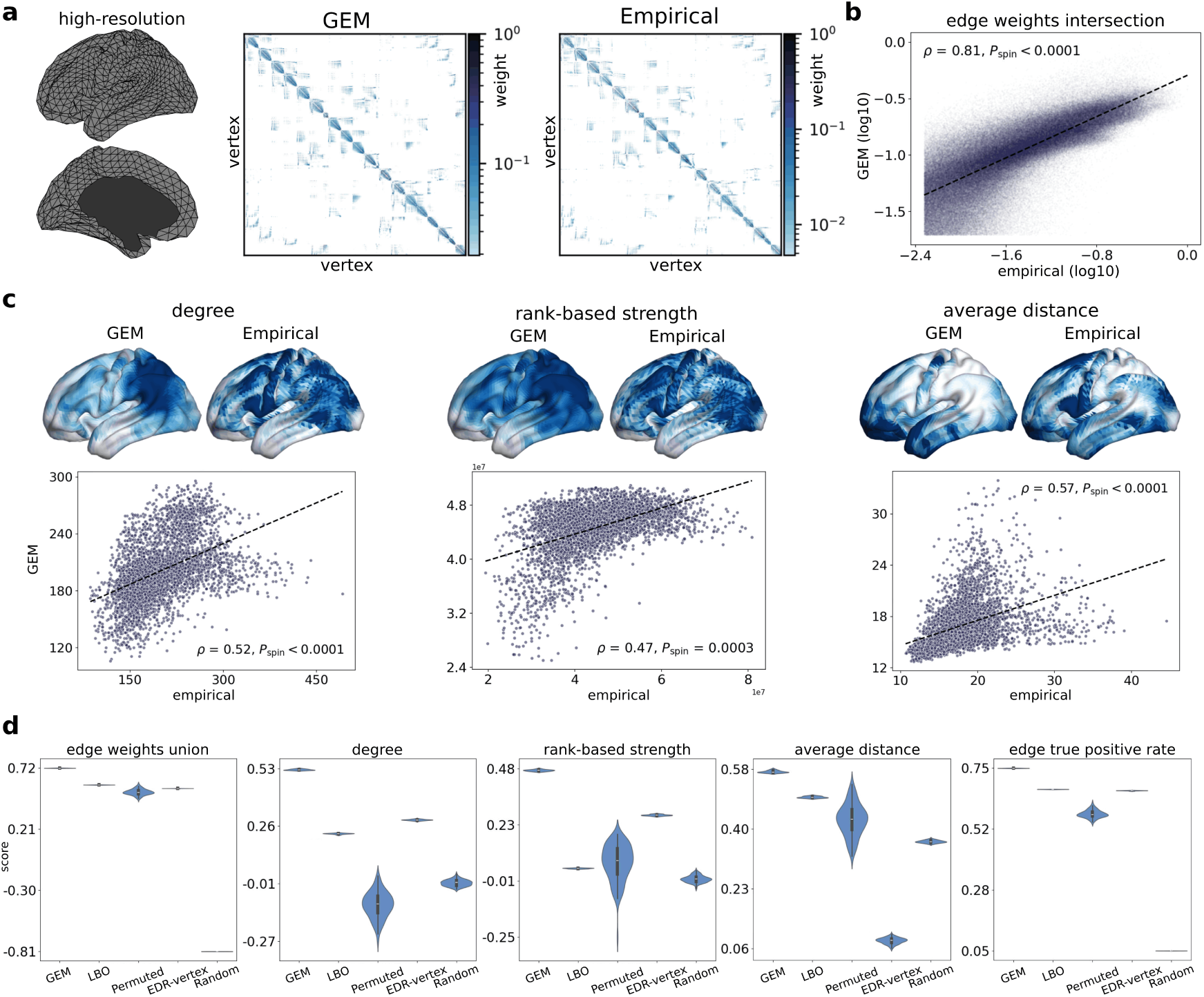
Modelling high-resolution human connectomes. **a**: Neocortical surface mesh along with optimized GEM and empirical high-resolution connectivity matrices mapped with diffusion MRI. The optimized model was generated using parameters *r*_*s*_ = 9.53 mm and 𝑘 = 108 modes, obtained by maximizing an objective function that quantifies similarities in binary and weighted topological and topographical properties of model and empirical connectomes (see Methods). **b:** Scatter plot of the common (intersection) edge weights between the GEM and empirical connectomes using the full dataset during optimization and testing. 𝑃_spin_ is the one-sided p-value estimated from 10,000 spin permutations. **c:** Spatial maps and scatter plots showing the topographical agreement between the optimized model and empirical data using the full dataset, quantified using Spearman rank correlations (𝜌). Panels from left to right show results for the degree (binary degree), rank-based strength (weighted degree), and average node connection distance. Low to high values on the spatial maps are coloured from white to dark blue. **d:** Comparison of the performance of the GEM relative to other models, as evaluated using split-half cross-validation. GEM=Geometric Eigenmode Model; LBO=Laplace-Beltrami Operator model approximating the connectivity of the mesh; Permuted=GEM computed with optimal parameters after randomizing the ordering of the geometric eigenvalues; EDR-vertex=Exponential Distance Rule model defined at the vertex level; Random=Random network defined at vertex resolution (see Methods for details).

We additionally consider two measures outside the objective function for capturing network topography and topology. For topography, we consider the node-averaged connection lengths, which encode key aspects of the connectome’s physical embedding. This measure shows a high correlation between the GEM and empirical data (𝜌 = 0.57) (Fig. 2c). These correlations are notable given the high spatial resolution of the empirical connectome, with previous studies showing that discrete graph-based models of low-resolution atlas-based connectomes cannot obtain spatial degree correlations exceeding 𝜌 = 0.2 (*7*). For topology, we consider the edge-wise true positive rate, defined as the fraction of edges present in the empirical connectome that are also retained in our model. This measure thus provides a precise quantification of the degree to which our model captures the binary topology of the empirical network. Here, we obtain a true positive rate of 75%, consistent with strong agreement between model and data.

To avoid over-fitting and allow fair comparison between the GEM and benchmark null models with different numbers of free parameters, we use a split-half cross-validation to optimize model parameters in a training set and obtain distributions of fit statistics in held-out test sets of the data (see Methods). Critically, the GEM outperforms all benchmark null models for all network metrics when cross-validated (Fig. 2d; see Fig. S2b for optimal model parameters). Moreover, the GEM’s performance under cross-validation closely matches the results obtained using the full dataset in Figs. 2b and c, demonstrating the robustness of the model’s predictions.

To further examine the model’s performance and generalizability, we consider two more complex features of network topography and topology that are not directly optimized in the objective function: the network community structure (topography) and spectral distance (topology). Community structure refers to the tendency of networks to organize into distinct subgroups of nodes, often called communities or modules, characterized by dense intra-modular and sparse inter-modular connectivity. This modular architecture has been observed across different species and spatial scales, and is widely recognized as a fundamental property of empirical brain networks (*84–94*). Spectral distance is defined as the distance between the eigenspectra of the normalized graph Laplacian of the model and empirical networks. These eigenspectra succinctly capture the core topological charac-teristics of each network, such as their degree distribution, motif and random walk architecture, and synchronizability (*95–99*). They thus offer an efficient summary of network topology that obviates the need for computationally intensive estimation of specific topological properties (e.g., shortest path lengths, motif spectra) on large matrices. For each metric considered, the GEM shows superior or comparable performance to all benchmark models (Figs. S3 and S4). The strong performance of the GEM in capturing empirical modular architecture is consistent with past work showing that such properties may be largely explained by physical constraints on connectome topology (*14, 23*). Together, these findings indicate that the GEM can successfully capture diverse aspects of highresolution connectome topology and topography. The comparison with benchmark models further indicates that the success of our model is attributable to the specific form of Equation 3, which captures the predicted effects of geometry on connectome architecture through the lens of NFT, rather than the low-order effects of cortical geometry or EDR-like connectivity on their own. In further support of this view, we show in Fig. S1b that the GEM’s performance declines more rapidly when long-wavelength modes are removed from the model first, consistent with theoretical and empirical evidence supporting the dominant contribution of lower-order modes to cortical organization (*37, 68, 69, 100, 101*), and the physical prediction that long-wavelength eigenmodes with low 𝜆 require less energy for excitation and are thus more strongly expressed (*62, 65, 68, 102*). Importantly, the GEM’s superior performance is robust to data processing choices such as whether the high-resolution connectomes are smoothed or not (Fig. S5) (*103*) or whether the connection weights are rescaled (Fig. S6) (*104, 105*) (for further explanation, see Methods and Ref. (*36*)). We further observe that the GEM performs better than all benchmark models in capturing connections across all distance ranges, but that all models struggle to capture rare, low-weight, and long-distance connections (Fig. S7). All models also fail to capture the spatial organization of nodal clustering observed in the empirical connectome (clustering reflects the tendency of neighboring nodes to form tightly interconnected groups) (Fig. S8), although the performance of the GEM in capturing this property improves substantially for atlas-based connectomes, as discussed in the next section. Our analysis thus far has relied on geometric eigenmodes extracted from a group-average neocortical surface template. We use this average surface for simplicity, because the lowest-order modes are common across brains, and because our analyses focus on a group-average connectome, which helps to minimize noise associated with mapping individual-specific connectomes with diffusion MRI (*106*). The spatial structure of the averaged surface modes can nonetheless differ from the modal structure of individual surfaces. To further evaluate the specific contribution of cortical geometry to connectome architecture, we fit the GEM to high-resolution connectomes of a subset of 100 individuals, using either the individual’s neocortical surface or the group-average template surface to derive the modes (see Methods). Despite the technical challenges in accurately mapping individual connectomes at high resolution (*103,107,108*), the performance of GEM fits to individual connectomes are comparable to the group-average connectome (Fig. S9). More importantly, using modes obtained from an individual’s cortical geometry better captures their connectome topography than when modes obtained from the group-average neocortical surface are used (Fig. S9c), underscoring the role of cortical geometry in shaping individualized connectome architecture. Together, these findings indicate that the mechanism captured by our simple model, which favours connections that facilitate excitation of resonant geometric modes of the cortex, plays a key role in shaping diverse aspects of the topology and topography of human connectomes mapped at high resolution.

## The GEM captures atlas-based topological and topographical properties of the human connectome

Despite the advantages of mapping human connectomes at high-resolution (*103, 109–114*), most connectomes are constructed using an atlas which parcellates the cortex into discrete regions that constitute the network nodes. To determine whether the GEM generalizes to connectomes mapped at the atlas level, we use the same group-average high-resolution connectome in the previous section but parcellate it into 150 regions using a well-validated atlas (*37,115*) (see Methods for details). We then parcellate the GEM-derived connectome using the same atlas, fit it to the empirical connectome (Fig. 3a), and evaluate its performance relative to four benchmark models. The first benchmark, termed ‘MI’, models connections based on wiring rules that specify a trade-off between wiring cost and homophilic attachment, with the latter operationalized using a topological measure called the matching index (MI) that promotes connectivity between pairs of nodes that are connected to other similar nodes. This model has successfully reproduced connectome topology in diverse species and is often identified as the optimal generative connectome model in head-to-head model comparisons (*5, 8, 34*). (Note that this model could not be used for high-resolution connectomes due to its computational burden.)

**Figure 3:**
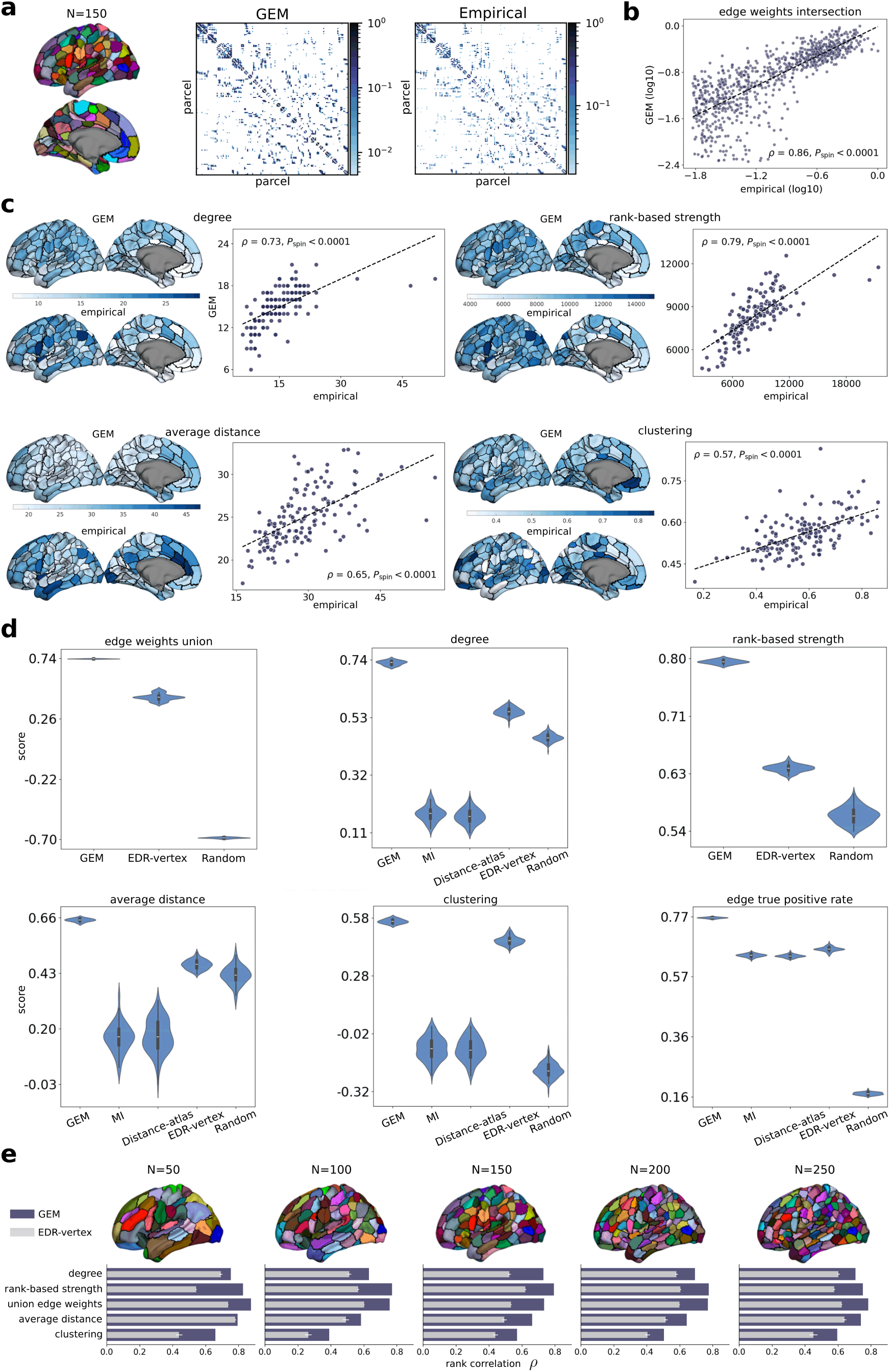
Modelling atlas-based human connectomes. **a:** Brain atlas (parcellation) used to divide the left hemisphere into N=150 regions (left). Optimized GEM and empirical atlas-based connectivity matrices (right). The optimized model was generated using parameters *r*_*s*_ = 9.53 mm and 𝑘 = 59 modes, obtained by maximizing an objective function that quantifies similarities in binary and weighted features of edge-level topology and node-level topography of model and empirical connectomes (see Methods). **b:** Scatter plot of the common (intersection) edge weights between the GEM and empirical connectome using the full dataset during optimization and testing. 𝑃_spin_ is the one-sided p-value estimated from 10,000 spin permutations (see Methods for details). **c:** Spatial maps and scatter plots showing the topographical agreement between the optimized model and empirical data using the full dataset, quantified using Spearman rank correlations (𝜌). From left to right, top panels show results for the degree (binary degree) and rank-based strength (weighted degree), while bottom panels show results for the average node connection distance and nodal clustering coefficient. **d:** Comparison of the performance of the GEM relative to other models, as evaluated using split-half cross-validation, for the 150 region-atlas. Note that the MI and Distance-atlas can only model binary connections, so we cannot evaluate their performance with respect to the rank-based strength and edge weights union correlation. GEM=Geometric Eigenmode Model; MI=Matching Index model; Distance-atlas=Power law distance rule model defined at atlas resolution; EDR-vertex=Exponential Distance Rule model defined at the vertex level and then parcellated; Random=Random network defined at vertex resolution and then parcellated (see Methods for details). **e:** GEM’s performance when parcellating the left cortical hemisphere at different resolutions (50, 100, 150, 200, and 250 regions). The GEM consistently shows strong performance, regardless of the parcellation resolution. The EDR-vertex benchmark is shown for comparison as it represents the second best model after the GEM.

The second benchmark, termed ‘Distance-atlas’, models connections following a stochastic, distance-constrained wiring process, with distances estimated between the centroids of parcellated atlas regions as commonly implemented in prior work (*5, 7, 8, 116, 117*). The third benchmark, termed ‘EDR-vertex’ is a parcellated version of the ‘EDR-vertex’ model used for the high-resolution connectome; i.e., distances are estimated, and connections are formed, at vertex resolution before being parcellated. The final benchmark, termed ‘Random’, is a parcellated version of the same random benchmark used for the high-resolution connectome. The last two benchmarks account for any biases associated with parcellating model networks defined at high-resolution (see Methods for further details of all benchmark models).

The MI and Distance-atlas benchmarks only capture binary aspects of the connectome. We therefore optimize these models using a different objective function that maximizes the sum of (1) the rank correlation of the binary node degree maps; and (2) the percentage of edges reproduced by the model (i.e., the edge-wise true positive rate). All other models are optimized with the same objective function used in the analysis of high-resolution connectomes. In secondary analyses, we compare all models after optimization using the binary-only objective function. We again perform split-half cross-validation to ensure fair comparison between models with different numbers of parameters (Fig. 3e).

Fitting the GEM to atlas-based empirical data, we find that the optimal parameters are *r*_*s*_ = 9.53 mm and 𝑘 = 59 modes, indicating that around half the number of modes required for the highresolution connectome are needed to represent the 150-region connectome of the left hemisphere, consistent with the low-pass spatial filtering effect of an atlas (*69*) (see Fig. S14a for optimal model parameters). For this parameter combination, we obtain a rank correlation between common (intersection) edge weights of the model and empirical connectomes of 𝜌 = 0.86 in the full sample (Fig. 3b). The GEM also reproduces the binary and weighted empirical degree sequences with high accuracy (𝜌 = 0.73 and 𝜌 = 0.79, respectively) (Fig. 3c, top). We again find that the GEM captures topological and topographical properties that were outside the objective function, reproducing 77% of empirical edges (topology) and achieving correlations for the average node connection distance and the clustering coefficient (topography and topology) of 𝜌 = 0.65 and 𝜌 = 0.57, respectively (Fig. 3c, bottom). As in the high-resolution connectome results in Fig. 2, the identified optimal parameters of the GEM and model performance statistics are highly consistent between the analysis of the full sample (Figs. 3b and c) and the cross-validated results (Fig. 3d; see Figs. S14a and b for the optimization landscape of non-cross validated results and optimal model parameters in the cross-validated results, respectively).

For comparison, the MI model reproduces on average 64% of edges, with a binary degree correlation of 𝜌 = 0.18 (Fig. 3d; see Fig. S14b for optimal model parameters). This result is consistent with past work showing that the MI model can reproduce elements of connectome topology but not topography (*6, 7, 35*). The GEM also surpasses the performance of the three other benchmark models (i.e., Distance-atlas, EDR-vertex, and Random) across all metrics (Fig. 3d), indicating that the success of the GEM cannot be attributed to any biases induced by parcellating a high-resolution connectome. Since the MI and Distance-atlas models can only construct binary connectomes, we also optimize the GEM and the EDR-vertex models with the same objective function for fair comparison (see Fig. S15 for optimal model parameters) and find that both the optimal parameters and the superior performance of the GEM remain unchanged (Fig. S16).

As in the analysis of high-resolution connectomes, we compare the modularity structure between the model and empirical connectomes and compute their spectral distances (see Methods). We find that the GEM captures these properties more accurately than all benchmark models (Figs. S17 and S18). The distance-dependence of the edge reconstruction accuracy for all models is also consistent with the high-resolution findings (Fig. S19 and see Ref. (*35*) for a discussion of this issue). Moreover, the strong performance of the GEM is robust to the choice of atlas resolution (i.e., nearly all 𝜌 ≥ 0.6 for resolutions spanning 50 to 250 regions; Fig. 3e), connectome edge density (Figs. S20 and S21; see Methods), connectome smoothing (Fig. S22 and see Ref. (*103*)), and edge-weight resampling (Fig. S23), supporting the model’s robustness and generalizability.

These results indicate that our simple model reproduces both topological and topographical properties of human empirical connectivity data mapped either at vertex or atlas resolutions. The GEM is thus parcellation-independent. Moreover, we show that its performance cannot be solely attributed to generic properties related to the spatial structure of the eigenmodes or EDR-like connectivity kernels (see Discussion) and that it performs better than current state-of-the-art models, supporting the hypothesis that the human cortical connectome is configured to facilitate the excitation of the resonant geometric eigenmodes of the cortex.

## The GEM generalizes to other mammals

Having demonstrated the robust performance of the GEM in capturing diverse aspects of human connectome topology and topography, we now test the model’s ability to reproduce connectome properties of four different non-human species: mouse (*118*), marmoset (*119*), macaque (*120*), and chimpanzee (*83, 121*). The connectomes are mapped with either non-invasive diffusion MRI (for the chimpanzee) or invasive tract-tracing (for the mouse, marmoset, and macaque) (Fig. 4a; see Methods for details) using a specific atlas tailored for each species, as chosen by the original data sources; i.e., 43, 55, 29, and 57 regions in one hemisphere for the mouse, marmoset, macaque, and chimpanzee, respectively. For the three primates, we calculate the geometric eigenmodes from a mesh representation of the neocortical surface of their left hemisphere comprising 37,974, 10,242, and 4,842 vertices for the marmoset, macaque, and chimpanzee, respectively. For the mouse, we calculate the geometric eigenmodes from a tetrahedral mesh representation of the right hemisphere isocortical volume comprising 7,364 vertices, since the empirical connectome is derived from anterograde tracers injected into the right hemisphere. All network models are optimized using the same (or comparable) objective function used for the analysis of human connectomes, as in Figs. 2 and 3 (see Methods).

**Figure 4:**
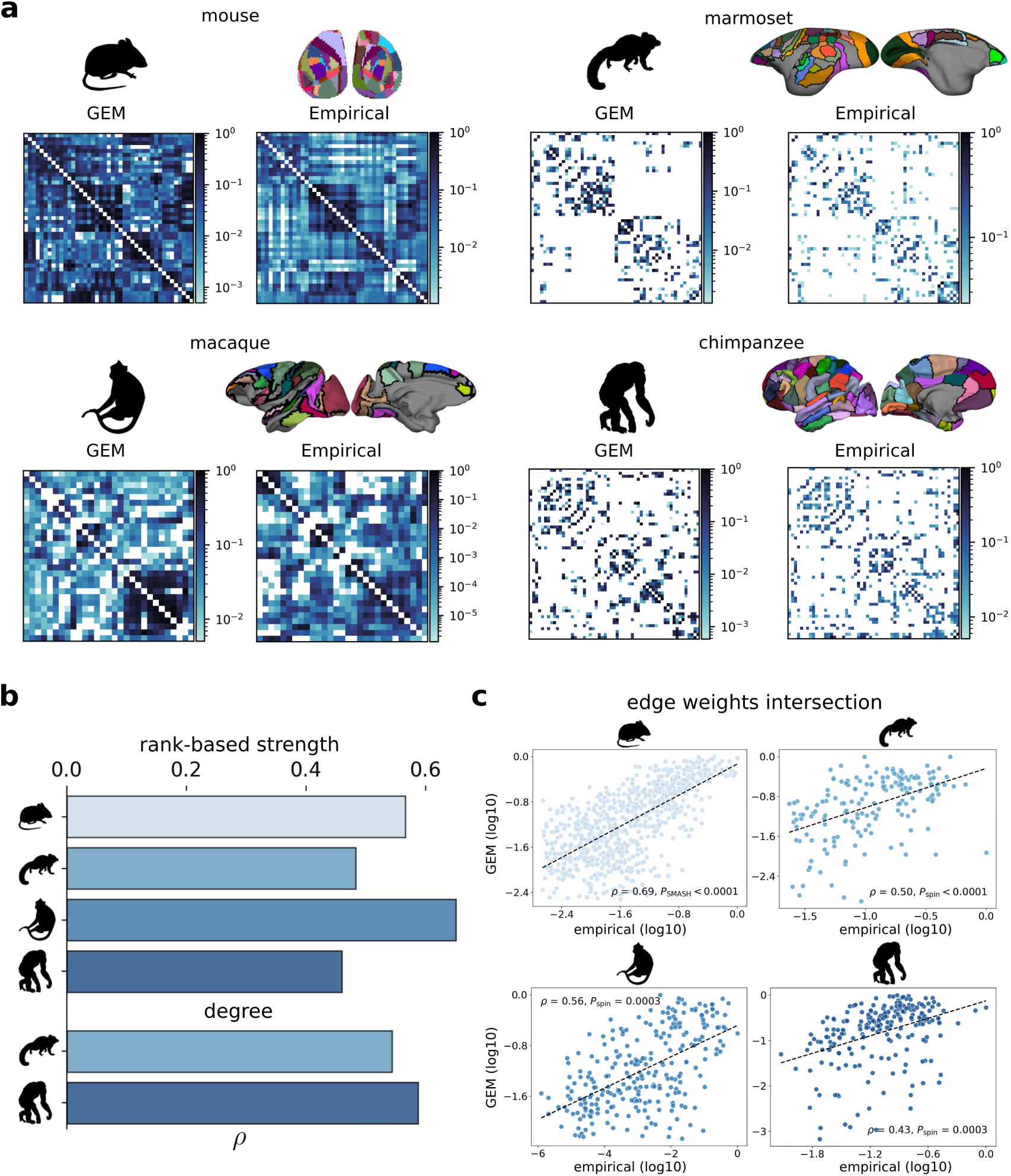
The GEM generalizes across species and imaging modalities. **a:** Brain atlases along with optimized GEM and empirical connectivity matrices for the mouse (43 regions), marmoset (55 regions), macaque (29 regions), and chimpanzee (57 regions). Brain atlases show superior/inferior views for the mouse and lateral/medial views for the primates. The chimpanzee connectome is derived from non-invasive diffusion MRI, while the other connectomes are derived from invasive tracer injections. Note that the marmoset and macaque connectomes only have partial coverage of the cortex. See Methods for details of the optimization and optimized model parameters for each species. **b:** Spearman rank correlations between the optimized model and empirical data for the weighted degree (rank-based strength; top) and binary degree (bottom) across species. Since the mouse and macaque connectomes are very dense (90% and 80% of all connections exist, respectively), we only evaluate the binary degree measure for the marmoset and chimpanzee (see Methods). **c:** Scatter plots of the common (intersection) edge weights between the GEM and empirical connectomes.

Figures 4b and c show that the GEM reproduces key topological and topographical features of the connectomes of all species investigated, with its performance being superior to the MI, Distance-atlas, EDR-vertex, and Random benchmark models defined for each animal (see Figs. S24, S25, S26, and S27 for the statistical evaluation, Fig. S28 for the topographical maps, and Figs. S29 and S30 for the performance on the dense mouse and macaque connectomes). In addition, we find that the GEM generally performs better in capturing empirical properties that are not included in the objective function, such as community structures (Figs. S31, S32, S33, and S34) and graph eigenspectra (Fig. S35). The GEM’s strong performance is also robust to the connectome and model edge thresholds (Figs. S36, S37, S38, S39, and S40), and resampling of edge weights (Figs. S41, S42, S43, and S44). As in the analysis in humans, the GEM outperforms other benchmarks at capturing connections of different lengths across all species (Fig. S45).

These findings indicate that the GEM generalizes across mammalian species, methods for mapping neuronal connectivity, and the specific choice of an atlas, supporting a central and highly conserved role for geometrically constrained processes in shaping connectome architecture.

## Discussion

Here we propose a simple model of connectome architecture that prioritizes physical and geometric constraints on brain connectivity, as predicted by the biophysical dynamics derived from NFT (*64, 65, 79*). The model recapitulates key aspects of both the binary and weighted representations of empirical connectome topology and topography in human (Figs. 2 and 3) and non-human species (i.e., mouse, marmoset, macaque, and chimpanzee; Fig. 4). The model relies solely on cortical geometry and is highly scalable, being applicable to connectomes mapped across a wide range of spatial resolutions and imaging modalities.

Mature cortical architecture is shaped by multiple processes, including axonogenesis, pathfinding, overgrowth, elimination and pruning, and activity-dependent plasticity (*45, 46, 122, 123*). These processes jointly influence which inter-regional connections are ultimately retained in adulthood, thus determining the binary topology of the connectome, and which connections are stronger than others, thus determining weighted connectome architecture. Our model suggests that cortical geometry constrains the mechanisms that underpin these biological processes in a way that favours the retention and strengthening of connections that facilitate the excitation of diverse, particularly long-wavelength, geometric eigenmodes. The GEM may also encompass the effects of geometric constraints on the spatial expression profiles of early patterning molecules that guide axons to their targets (*13,37*), although these effects are not directly addressed in the present work. More precisely, our model represents a low-rank approximation of the Green’s function associated with the time-averaged dynamics of the NFT developed in Ref. (*65*). In this model, and many other variants of NFT (*47, 58–61, 124, 125*), neuronal activity propagates as waves through a continuous cortical medium. Geometric eigenmodes of the cortex correspond to the standing waves of these dynamics, and thus represent a fundamental basis set that can be used to reconstruct any spatiotemporal pattern expressed by the cortex in the linear regime (*67–69, 126*). The superior performance of the GEM thus aligns with mounting empirical evidence that wave-like activity plays a crucial role in brain development and synaptic plasticity (*127–130*).

Here we used a minimalist approach in developing our model, trying to use simplifying assumptions where possible (e.g., we relied on a temporal mean-field approximation) to allow us to derive an analytic expression for modelling connectome architecture. Rather than serving as a detailed biophysical model of all the intricacies of cortical wiring through development, our model illustrates the more general influence of geometry and wave-like dynamics (approximated here in the zero-frequency limit) in such processes. Our model also extends theoretical work defining formal links between spectral representations of structural and functional connectivity (*64, 131*).

One potential interpretation of our results is that connectome architecture is shaped by the resonance of spatial eigenmodes, such that brain regions with similar eigenmode profiles will be preferentially connected due to the energy-efficient and frequency-matched communication pathways between them (*78, 79, 110, 132*), which facilitates eigenmode excitation. Resonant processes are ubiquitous in biological systems, including the brain, where neural circuits exploit resonance in temporal dynamics such that inputs matching the intrinsic frequency of individual neurons are selectively amplified to support frequency-based information routing, coherent communication, and adaptive reconfiguration of network activity during different cognitive states (*79, 133–137*). Although our model was derived in the static limit of NFT, future work could incorporate dynamics to better explore the potential influence of such resonant processes.

Within our model, the contribution of each mode in Equation 3 reflects its relative expression in the assumed mean-field neural dynamics (*65*). More specifically, the GEM favours contributions from eigenmodes with long wavelengths as a direct consequence of the transformation of the eigenvalues of the LBO via the term (1 + *r*^2^𝜆_*m*_)^−1^ in Equation 3 (*79*); hence, low-order modes with smaller eigenvalues exert a stronger influence on connection weights between pairs of cortical locations (Figs. S1a and b). This dominance of low-order modes aligns with past theoretical and experimental work underscoring a major contribution of these modes to cortical organization and dynamics (*37, 68, 69, 100, 102, 138, 139*). These observations likely result from a more general physical principle in which higher levels of energy are required to excite modes with shorter wavelengths (or with higher spatial frequencies), which in turn makes their excitations weaker (*62, 65, 68, 102*). This energy dependence is analogous to the well-known Boltzmann distribution in statistical mechanics, which describes how the probability of observing a system in a certain state decreases exponentially as a function of the energy associated with that state. In our model, shorter-wavelength modes are attenuated via the length scale parameter *r*_*s*_ and the natural ordering of the eigenvalues of the LBO (see Fig. S11).

We note though that the connectome model presented here focuses exclusively on intra-hemispheric cortico-cortical connectivity and does not account for inter-hemispheric connections. This focus on a single hemisphere aligns with many prior modelling studies in the literature (*5–7*) and stems from the limited ability of diffusion MRI to accurately reconstruct long-range inter-hemispheric tracts and the limited data available for tracer-based connectomes in non-human species, which generally only offer comprehensive coverage for a single hemisphere (*20, 118, 120*). Furthermore, our model does not consider connectivity with non-neocortical areas, which play an important role in shaping cortical organization and whole-brain dynamics (*13, 39, 140–147*). We also assume a uniform spatial scale parameter *r*_*s*_ across the cortex, despite evidence that different neuronal populations exhibit different characteristic propagation scales (*81*). The strong performance of the GEM suggests that these details are not necessary to capture many aspects of empirical connectome data. More detailed models that account for spatial heterogeneities, non-neocortical interactions, and more intricate physiological processes may improve model fits, but come at the cost of additional model parameters and complexity. In lieu of such complexity, the sufficiency of our simple model in capturing many key aspects of connectome topology and topography across diverse measurement techniques, resolution scales, and species separated by millions of years of evolution underscores the universal and highly conserved role that geometry plays in shaping cortical connectome architecture.

## Acknowledgments

This research was supported by Monash University’s eResearch capabilities, including the M3 MASSIVE high-performance computing facility. HCP data were provided by the HCP, Wu–Minn Consortium (principal Investigators D. Van Essen and K. Ugurbil; no. 1U54MH091657) funded by the 16 NIH Institutes and centres that support the NIH Blueprint for Neuroscience Research, and by the McDonnell Center for Systems Neuroscience at Washington University. We thank Dr Ben D Fulcher and Dr Patrick Desrosiers for helpful discussions regarding this manuscript and Dr Martijn van den Heuvel for providing the chimpanzee connectome data.

## Author contributions

F.N., M.G., T.C., J.C., P.A.R., J.C.P. and A.F. conceptualized the model. F.N. performed the investigation. F.N., J.C.P. and A.F. administered the project. F.N. developed visualizations. M.G., J.C.P., A.S., and S.O. generated data. A.H. reviewed the code. J.C.P. and A.F. acquired funding. J.C.P. and A.F. supervised the project. F.N., J.C.P. and A.F. wrote the original draft. All authors reviewed and edited the final manuscript.

## Data and materials availability

Data and code will be made available upon acceptance of the manuscript.

