## Supplementary Material for "Geometric constraints on the architecture of mammalian cortical connectomes"

#### Supplementary materials

Materials and Methods

Figs. S1 to S45

### Supplementary Materials for
### Geometric constraints on the architecture of mammalian cortical connectomes

Francis Normand*, Mehul Gajwani, Trang Cao, Jace Cruddas,

Arshiya Sangchooli, Stuart Oldham, Alexander Holmes, Peter Robinson,

James C. Pang†, Alex Fornito†

†These authors contributed equally to this work.

**This PDF file includes:**

Materials and Methods

Supplementary Figures

Figures S1 to S45

#### 0.1 Materials and Methods

##### 0.1.1 Generating human connectomes

Our analysis begins with individual human connectomes and a neocortical surface mesh of the left hemisphere initially mapped in a left–right symmetric version of FreeSurfer's fsaverage population-averaged template space at a high resolution of 32,492 vertices (fsLR-32k) (*1*). For computational purposes, we downsampled the midthickness and white surfaces onto a mesh of 4,842 vertices. which became 4,386 vertices after excluding the medial wall ('fsLR-5k'), using tools in the Connectome Workbench `https://www.humanconnectome.org/software/connectome-workbench`. Specifically, we generated a regularly spaced spherical surface mesh of 4,842 vertices (fsLR-5k) using the *surface-create-sphere* command with an input of 5,000 desired number of vertices (to achieve regular spacing between vertices. We then used this spherical surface to downsample the fsLR-32k midthickness and white surfaces with the command *surface-resample*. The downsampled surface was then parcellated using different cortical atlases (see below) using Connectome Workbench's *label-resample* command.

Individual structural connectivity matrices were generated for 339 healthy, unrelated young adult participants (ages 29 ± 4 years) in the Human Connectome Project (HCP) (*2*). Minimally-processed T1-weighted and multi-shell diffusion MRI data and derivatives provided by the HCP were used for connectome construction. The MRI data were acquired on a custom Siemens 3T Connectome Skyra scanner, with diffusion MRI data acquired along 270 directions at three b-value shells (b = 1000, 2000, 3000 s/mm$^2$, 90 volumes per b value). Minimal processing of diffusion MRI data by the HCP included intensity normalization, distortion corrections, motion correction, and gradient nonlinearity correction, before the derivation of a brain mask (*3, 4*). We adapted a previously validated pipeline to construct structural connectomes with quantitative streamline tractography (*5, 6*). Briefly, white matter, gray matter, and cerebrospinal fluid response functions were estimated for each individual (*7*). These response functions were then used to estimate multi-tissue orientation distribution functions from all available b-value shells using constrained spherical deconvolution (*8*), the results of which were normalized in the log-domain (*9*). White matter fibre orientation distribution images were used for whole-brain tractography with a probabilistic algorithm (*10*), which was constrained by a tissue-segmentation image generated from high-resolution structural images (*11*). Ten million streamlines were generated, with random seeding from the gray-white matter boundary in the tissue segmentation image to mitigate gyral bias (*6*). Streamlines were then weighted with the SIFT2 (Spherical-deconvolution Informed Filtering of Tractograms-version 2) algorithm (*12*). Weighted streamlines were then assigned to each individual's surface mesh vertices to construct their high-resolution connectome, with each streamline endpoint assigned to the closest vertex and streamlines discarded if no surface vertices were present within a 2-mm radius of either endpoint (*13*). Surface files used for streamline assignment were in MNI space, and thus tractogram endpoints were warped to MNI space before assignment (surfaces and non-linear warps were provided by the HCP). All diffusion MRI analysis and tractography steps were implemented using MRtrix3, version 3.0.4 (*14*).

To construct group-average connectomes in the fsLR-5k surface space and for atlas-based analyses, we first generated a group-average connectome in the fsLR-32k space by summing individual structural connectivity matrices. Aggregating the connectivity matrices at this resolution yields a more accurate representation of finer details of individual connectomes that can be lost if the connectomes are first downsampled prior to being combined. To project this group-average connectome from the fsLR-32k onto the fsLR-5k mesh, we established a vertex-wise correspondence by assigning each vertex in the fsLR-32k to its nearest neighbor in Euclidean space on the fsLR-5k mesh, which allocated each of the fsLR-5k vertices at least once. This mapping was then used in the same way that a parcellation is used (see section 0.1.6).

We also applied a smoothing kernel using connectome spatial smoothing (*15*) on the high-resolution group-average connectome mapped in fsLR-32k space. This technique can enhance connectome reliability and mitigate artifacts associated with high-resolution analyses of diffusion MRI data (*15*), but can also accentuate geometric effects on connectome architecture (*16*). For these reasons, we reported results using both smoothed (Fig. 2) and non-smoothed connectomes (Fig. S5). Following prior recommendations, we applied a connectome-based smoothing kernel with full-width half-maximum (FWHM) of 8 mm and truncation of the kernel $\epsilon = 0.01$, which is defined as the fraction of the kernel integral discarded as a result of kernel truncation (*15*). We used this smoothing kernel throughout our study, applied to the fsLR-32k group-average connectome, which was then downsampled to fsLR-5k or parcellated using a predefined atlas.

Our analysis of high-resolution connectomes was performed at the intrinsic resolution of the fsLR-5k mesh. As the vast majority of human connectome analyses are performed after the data have been parcellated with some regional atlas, we additionally created a series of atlas-based connectomes by parcellating the high-resolution connectomes using atlases of varying resolutions, as defined in Ref (*17*), and which have been extensively validated (*17–19*). Specifically, we used the Schaefer 17 networks 100, 200, 300, 400 and 500 region atlases, corresponding to 50, 100, 150, 200 and 250 regions in the left hemisphere to parcellate the high-resolution connectivity data using the procedure detailed in section 0.1.6. We therefore obtain five atlas-based group-averaged connectomes spanning different parcellation granularities, in addition to the high-resolution mesh-based connectome.

A common procedure in the analysis of human connectomes mapped with diffusion MRI involves thresholding the connectivity matrix to remove potentially spurious connections and thus improve the signal-to-noise ratio of the data. In the absence of any gold standard criteria for selecting a particular threshold, investigators rely on various heuristics (*20–24*). In our analysis, we estimated the expected edge density of the group-average connectome in fsLR-5k by projecting the 339 individual connectomes from fsLR-32k to fsLR-5k. Individual edge densities ranged from 4% and 6%, with a mean of 4.6%. We used this mean as our target density and thresholded the group-average connectome to keep the top 4.6% of edge weights, setting all other matrix entries to zero.

For the group-average atlas-based connectomes, our main analyses focused on connectomes with empirical edge densities set to 15%, 12.5%, 10%, 7.5%, and 5% when mapped at resolutions of $N$=50, 100, 150, 200, and 250 regions, respectively (Fig. 3). In secondary analyses, we also examined model performance for a range of 15 equally spaced empirical connectome densities between 5% and 35% on the Schaefer300 atlas ($N$ = 150) in Fig. S20. In Fig. S20, to ensure that all high-resolution models (i.e., GEM, EDR-vertex, and Random) retained enough edges to match the empirical edge density after being parcellated, we used a high-resolution model edge density threshold of 4.6% for the first four empirical edge densities (i.e., 5% to 11%), 15% for the next seven (13.5% to 26%) and 20% for the last four (28.5% to 35%).

For the 100 high-resolution connectomes mapped at the level of individual participants in Fig. S9, we simply smoothed the individual fsLR-32k connectomes and then projected them onto fsLR-5k surface space. We used the same smoothing kernel parameters and edge density (i.e., 4.6%)

for all individual connectomes as in the group-average connectome.

In all human analyses, we focused on the cortico-cortical connectome of the left hemisphere. This focus aligns with past modelling work (*25–29*), which has generally only considered intra-hemispheric cortical connectivity, due to the known difficulties of diffusion MRI in reconstructing long-range inter-hemisphere tracts (*30–33*).

##### 0.1.2 Non-human primate and mouse connectomes

Human connectomes are generally mapped with diffusion MRI, which has known limitations in accurately reconstructing axonal connectivity (*30–33*). To ensure that our results were not simply due to artifacts related to such limitations, we additionally evaluated model performance with respect to connectomes defined using gold standard invasive tract-tracing measurements in mouse, marmoset, and macaque. We also investigated the chimpanzee connectome, as mapped with diffusion MRI, to further assess the generalizability of our model across species.

Tract-tracing is an invasive technique used to map axonal projections using viral tracers. In these experiments, labeled viruses are injected within a target cell population $i$ and then transported either anterogradely or retrogradely along the axonal pathways from or to the injection site, respectively, infecting cells in distally connected population, $j$. Two-photon microscopy is then used to reconstruct these projections by counting the number of infected neurons in cell population $j$, thereby providing a quantitative measure of the directed connection strengths linking cell populations $i$ and $j$. Empirical connectomes derived from tracer injections are therefore usually directed (*34–36*), meaning that the strength of connection from region $i$ to $j$, $w_{ij}$, is not always equal to the strength of connection from region $j$ to $i$, $w_{ji}$. As our model does not try to capture the directionality of connectivity (see section 0.1.4), we symmetrized these empirical connectomes by taking the average of their upper and lower triangular entries. Diffusion MRI cannot resolve the directionality of axonal projections, so the human and chimpanzee connectomes were intrinsically undirected.

**Chimpanzee**

The chimpanzee connectome was obtained from previously published studies, specifically the group-average connectome reported in Ref. (*37*), derived from individual connectomes previously obtained via diffusion MRI in 22 female chimpanzees, as described in Refs. (*38, 39*). Each connectome was parcellated into 57 areas per hemisphere using the Desikan-Killiany atlas (*40*) (see

Fig. 4a, bottom right). We focused exclusively on the left hemispheric connectome, which had an edge density of 19% (see Figs. 4, S27, and S34). Given that the empirical connectome was already sparse, we did not conduct additional analyses across different edge densities.

**Macaque**

The macaque connectome was was obtained from previously published studies, generated through single injections of fluorescent retrograde tracers in 28 macaque monkeys (*35*). We used a subset of the left hemispheric connectome mapped between 29 of the 91 cortical areas defined by a parcellation of the entire cortical sheet (*41, 42*) (see Fig. 4a, bottom left). We focused on the $29 \times 29$ connectivity matrix where coverage was complete. The directed connection strength from area $j$ to the injected area $i$ was quantified as the extrinsic Fraction of Labeled Neurons (FLNe), which is the ratio between the number of neurons projecting to area $i$ from area $j$ and the total number of neurons projecting to area $i$ from all other areas minus the number of neurons identified in $i$ (see Ref. (*35*) for more details about the connectome generation process). After symmetrization of the edge weights, the connectome had an edge density of ~80%, which we refer to as the 'dense' connectome (see Figs. 4 and S30). The matching index (MI) and Distance-atlas models only consider binary topology, but many classical network properties such as node degree are essentially meaningless in dense networks, so we additionally considered a sparser representation of the connectome that retained the top 40% most strongly weighted edges, denoted as the 'sparse' connectome (see Figs. S26 and S33). Since this edge density of 40% was selected arbitrarily, we additionally evaluated model performances across edge densities ranging between 10% and 80% (see Fig. S37).

**Marmoset**

The marmoset connectome was obtained from previously published studies, generated by aggregating results from 143 retrograde tracer injections in 52 young adult marmosets by Ref. (*36*). The connectome was mapped between 55 of the 116 cortical areas defined by the Paxinos et al. stereotaxic atlas of the marmoset cortex (*43*) (see Fig. 4a, top right). As in the macaque, the connection strength for the marmoset was quantified with FLNe. The empirical connectome was already sparse, with an edge density of 16% after being symmetrized (see Figs. 4, S25, S35b, and S32). We therefore did not perform additional analyses using different edge densities for this species. The sparsity of the marmoset connectome may reflect the fact that this connectome was generated through the collation of multiple individual tracer studies rather than through a single concerted effort.

**Mouse**

The mouse connectome was obtained from previously published studies (*44*), derived from the data released by (*34*) and (*45*). Full details regarding the connectome generation pipeline can be found in the references cited. Briefly, 428 anterograde tracer injections were made in the right hemisphere of C57BL/6J male mice and imaged at a resolution of 100 $\mu$m (*34, 45*). The data were then resampled according to a Voronoi diagram based on the Euclidean distance between neighbouring voxels, to make the computation of the connectome computationally tractable (*44*). We used the symmetrized right hemisphere data released by (*44*). Finally, the connectivity estimates from the Voronoi neighbourhoods were aggregated into the 43-region isocortical parcellation defined by the Allen Mouse Reference Atlas (*46*) (see Fig. 4a, top left). This aggregation was done using the normalized connection density (NCD), corresponding to the same definition of edge weights used for the connectome in Ref. (*47*). The NCD is defined as the number of connections per unit volume in one region to one unit volume in the other region. The connectivity matrix was then symmetrized by taking the average of the upper and lower triangular matrix entries.

The resulting connectome was fully connected (i.e., edge density of 100%). For many of our models, we were unable to generate fully connected networks after parcellation (see below), so we thresholded the empirical data to 90%, representing the highest density that could also be matched by all models. We hereafter refer to this thresholded connectome as the 'dense' connectome (see Figs. 4 and S29). We additionally considered a sparser version of this dense connectome by retaining the edges with weights among the strongest 42%. This threshold was chosen as it represents the maximum threshold such that the network remains fully connected (no isolated nodes), as previously considered in Ref. (*44*) (see Figs. S24 and S31). We also looked at model performances across a range of edge densities between 10% and 90% (see Fig. S36). As in the analysis of macaque data, these sparser variants allowed a more meaningful consideration of binary topological properties.

##### 0.1.3 Derivation of cortical geometric eigenmodes

For humans and the other primates (i.e., chimpanzee, macaque, and marmoset), we extracted the geometric eigenmodes of their neocortical surfaces, parametrized as a 2D triangular mesh embedded within 3D Euclidean space. The Laplace-Beltrami operator (LBO) of the surface mesh fully describes the geometry of the manifold and is defined as (*48, 49*)

$$\Delta := \frac{1}{W}\sum_{i,j}\frac{\partial}{\partial x_i}\left(g^{ij}W\frac{\partial}{\partial x_j}\right), \tag{S1}$$

where $x_i$, $x_j$ are the local coordinates, $g^{ij}$ is the inverse of the inner product metric tensor $g_{ij} := \langle \frac{\partial}{\partial x_i}, \frac{\partial}{\partial x_j} \rangle$, $W := \sqrt{\det(G)}$, where det is the determinant, and $G := g_{ij}$ is the metric tensor (not to be confused with the Green's function, $\mathbf{G}$).

The geometric eigenmodes were derived by solving the eigenvalue problem associated with the LBO, also called the Helmholtz equation,

$$\Delta\psi = -\lambda\psi, \tag{S2}$$

where $\psi$ are the geometric eigenmodes, which represent the fundamental resonant patterns of excitation, or standing waves, of the neocortical surface. These modes are ordered from long to short wavelengths following the ordering of the eigenvalues ($0 \leq \lambda_1 \leq \lambda_2 ..., \leq \lambda_N$), where $N = 4,386$ is the number of vertices in the neocortical surface mesh. In the linear regime, which reasonably approximates normative, non-seizure-like macroscale dynamics (*50,51*), any spatiotemporal pattern on the neocortical surface can be represented as a weighted sum of geometric eigenmodes (*49*).

To approximate the continuous geometric eigenmodes, the LBO was discretized using finite element methods (*48*), with the discrete geometric eigenmodes computed from the neocortical surface mesh (Fig. 1a). The discrete LBO is an approximation of the negative continuous LBO; i.e., $\Delta \approx -\Delta_d$, where the subscript $d$ denotes the discretized approximation of the continuous LBO $\Delta$. We can thus write this eigenvalue problem in matrix form as

$$\mathbf{A\Psi} = \mathbf{B\Psi\Lambda} \tag{S3}$$

$$\implies \mathbf{B}^{-1}\mathbf{A\Psi} = \mathbf{\Psi\Lambda}, \tag{S4}$$

where $\mathbf{A}$ is the stiffness matrix, $\mathbf{B}$ is the mass matrix, $\mathbf{\Psi}$ is the matrix of the $N$ geometric eigenmodes along the columns, and $\mathbf{\Lambda} = \text{diag}(\lambda_N)$ is the diagonal matrix of eigenvalues. To obtain an analogous form to the continuous Helmholtz equation in Equation S2, we define the discretized LBO as

$$\Delta_d := \mathbf{B}^{-1} A = \boldsymbol{\Psi}\boldsymbol{\Lambda}\boldsymbol{\Psi}^{-1}, \tag{S5}$$

where $\mathbf{B}^{-1}$ is the matrix inverse of the mass matrix $\mathbf{B}$, $\boldsymbol{\Psi}^{-1} = \boldsymbol{\Psi}^T\mathbf{B}$ represents the matrix inverse of $\boldsymbol{\Psi}$.

In humans, we obtained the geometric eigenmodes and eigenvalues of the fsLR-5k midthickness surface (*1*) by solving the generalized eigenvalue problem in Equation S3 using the python library LaPy (*48, 52*). We repeated the same procedure for the left-hemisphere neocortical surfaces of individual human, marmoset, macaque, and chimpanzee brains. For the mouse, we used the right isocortical hemisphere because its empirical connectome was more comprehensively mapped in this hemisphere (*34*). Because the mouse data were defined in volume space, we modelled the 3D isocortex as a tetrahedral mesh, as per prior methods (*49*).

###### 0.1.4 Derivation of the Geometric Eigenmode Model (GEM)

The following general differential equation can be used to describe neuronal population dynamics on the neocortical surface by specifying how synaptic responses $\phi$ are generated from a source of activity $Q$

$$\mathcal{D}\phi(\mathbf{r}, t) = Q(\mathbf{r}, t), \tag{S6}$$

where the differential operator $\mathcal{D}$ prescribes the physical model underpinning the propagation of synaptic activity $\phi$ at location $\mathbf{r}$ and time $t$. We employ a simple NFT wave model (*49, 51*) that formalizes synaptic activity as a damped wave without regeneration (*53*)

$$\mathcal{D} = \frac{1}{\gamma_s^2}\frac{\partial^2}{\partial t^2} + \frac{2}{\gamma_s}\frac{\partial}{\partial t} + 1 - r_s^2\Delta, \tag{S7}$$

where $\Delta$ is the LBO associated with the geometry of the neocortical surface, $\gamma_s = v/r_s$ is the damping rate of the wave, $v$ is the characteristic axonal propagation velocity, and $r_s$ is the characteristic range of axons that defines the spatial length scale of wave propagation.

Equation S6 can be equivalently written in propagator form as an integral over space and time

$$\phi(\mathbf{r}, t) = \int\int G(\mathbf{r}, t, \mathbf{r}', t')Q(\mathbf{r}', t')d\mathbf{r}'dt', \tag{S8}$$

where the spatial propagator or Green's function, $G$, is identified as the structural connectome (*54*), and corresponds to the mean propagator for axons projecting to $(\mathbf{r}, t)$ from $(\mathbf{r}', t')$. The Green's function represents the response of a system to a unit impulse $\delta$, with

$$\mathcal{D}\, G(\mathbf{r}, t, \mathbf{r}', t') = \delta(\mathbf{r}' - \mathbf{r}, t - t'), \tag{S9}$$

where $\delta$ is the Dirac-delta function. In the linear regime, if the response of a system subject to a point source $\delta$ is known via the Green's function, one can obtain the system's response to a general source $Q$ as in Equation S8.

We derived a purely spatial propagator by considering the average synaptic field $\phi$ over time, i.e., $\phi(\mathbf{r}, t) \to \bar{\phi}(\mathbf{r})$, corresponding to the zero-frequency component ($\omega = 0$) in the Fourier temporal domain. In doing so, the differential operator $\mathcal{D}$ in Equation S7 can be simplified to

$$\mathcal{D} = 1 + r_s^2 \Delta_d, \tag{S10}$$

with a sign flip because we consider the discrete approximation of the LBO. Hence, the propagator equation becomes

$$\bar{\phi}(\mathbf{r}) = \int G(\mathbf{r}, \mathbf{r}') Q(\mathbf{r}') d\mathbf{r}'. \tag{S11}$$

Using eigendecomposition of the discretized LBO in Equation S5, we can write the definition of a Green's function in Equation S9 in matrix form as

$$\mathcal{D}\mathbf{G} = \mathbf{I}, \tag{S12}$$

$$\left[\mathbf{I} + r_s^2 \Delta_d\right]\mathbf{G} = \mathbf{I}, \tag{S13}$$

$$\left[\mathbf{I} + r_s^2 \mathbf{\Psi}\mathbf{\Lambda}\mathbf{\Psi}^{-1}\right]\mathbf{G} = \mathbf{I}, \tag{S14}$$

where $\mathbf{I}$ is the identity matrix. We can use the completeness relationship $\mathbf{\Psi}\mathbf{\Psi}^{-1} = \mathbf{\Psi}\mathbf{\Psi}^T\mathbf{B} = \mathbf{I}$ of the basis set formed by the eigenmodes to simplify the expression and factor the identity matrix

$$\left[\mathbf{\Psi I \Psi}^{-1} + r_s^2 \mathbf{\Psi \Lambda \Psi}^{-1}\right]\mathbf{G} = \mathbf{I}, \tag{S15}$$

$$\left[\mathbf{\Psi}[\mathbf{I} + r_s^2 \mathbf{\Lambda}]\mathbf{\Psi}^{-1}\right]\mathbf{G} = \mathbf{I}, \tag{S16}$$

$$\implies \mathbf{G} = \left[\mathbf{\Psi}[\mathbf{I} + r_s^2 \mathbf{\Lambda}]\mathbf{\Psi}^{-1}\right]^{-1}. \tag{S17}$$

Finally, the solution for the Green's function is (*54, 55*)

$$\mathbf{G} = \mathbf{\Psi}[I + r_s^2 \mathbf{\Lambda}]^{-1}\mathbf{\Psi}^{-1}. \tag{S18}$$

We may rewrite the equation above in local coordinate notation as

$$G_{ij} = \sum_{m=1}^{N} \frac{\psi_m(i)\psi_m^{-1}(j)}{1 + r_s^2 \lambda_m}, \tag{S19}$$

where $G_{ij}$ is an entry in the matrix $\mathbf{G}$ and $(i, j)$ refer to spatial locations $(\mathbf{r}, \mathbf{r}')$ introduced earlier. The summation runs over all $N$ geometric eigenmodes of the full basis set, including the first mode $\psi_1$ with constant amplitude. The denominator $(1 + r_s^2 \lambda_m)$ in Equation S19 attenuates the contribution of shorter-wavelength modes relative to longer-wavelength modes (see also Fig. S1a).

The Green's function in Equation S19 can only be derived when considering the full basis set $\mathbf{\Psi}$, because using a subset of the basis yields non-invertible matrices and dimension discrepancies. To relax this constraint on the modelling, we applied a low-rank approximation of the Green's function in Equation S19 and used only a subset of $k$ modes out of the total $N$ modes. This low-rank formulation is justified by three considerations. First, empirical connectomes themselves are likely of low rank, as suggested by recent work indicating that the bulk of the spectral power of many complex systems is contained within the first few leading eigenvectors (*55, 56*) (see Fig. S12 for a visualization of this effect in the high-resolution human connectome). Second, we expect that not all geometric modes will contribute to the formation and reinforcement of axonal connections, since shorter-wavelength modes are more difficult to excite (*53*). Finally, the eigenvalues and associated geometric modes obtained with the finite element method progressively diverge from their true values for shorter-wavelength modes, as the discrete nature of the surface mesh leads to numerical imprecision in the computations. As such, only a subset of all the geometric modes and eigenvalues can be considered a fair approximation of the true modes (*48*). As an example of this imprecision, we show in Fig. S11 the eigenvalues obtained with finite element modelling from three different spherical surface meshes with different resolutions, each containing about ~1,000 ('fsLR-1k'), 5,000 ('fsLR-5k'), and 10,000 ('fsLR-10k') vertices, as estimated using Equation S3. These numerical eigenvalues are compared to the ground truth that can be analytically derived from continuous spherical harmonics. The figure shows three key findings: (1) numerical eigenvalues obtained from higher-resolution meshes better approximate the ground-truth eigenvalues compared to lower-resolution meshes; (2) the first 200 eigenvalues obtained from the fsLR-5k mesh used for our analyses represent a good approximation of the ground-truth spherical harmonics; and (3) there is a marked divergence between analytic and numerical estimates beyond the first ~400 eigenvalues. Although ground-truth estimates are not available for the cortex, the topological and geometric similarity between the cortex and sphere allow us to use this analysis as a rough indication of the numerical error one can expect when estimating eigenmodes on the neocortical surface (*51*).

These considerations motivate a low-rank approximation as a more accurate and parsimonious account of connectome architecture. We therefore defined our connectome model, termed the Geometric Eigenmode Model (GEM), as

$$G_{ij} = \sum_{m=1}^{k} \frac{\psi_m(i)\psi_m^+(j)}{1 + r_s^2 \lambda_m}, \tag{S20}$$

or in matrix form as

$$\mathbf{G} = \mathbf{\Psi}_k [\mathbf{I} + r_s^2 \mathbf{\Lambda}]^{-1} \mathbf{\Psi}_k^+, \tag{S21}$$

where $\psi_m^+$ is the $m$th column vector of the Moore-Penrose matrix inverse (pseudoinverse) of $\mathbf{\Psi}_k$, which is used to preserve the orthonormality of the first $k$ geometric modes. The pseudoinverse $\mathbf{\Psi}_k^+$ is used in Equation S21 instead of $\mathbf{\Psi}_k^T \mathbf{B}$, because the latter orthonormalizes the modes with respect to the full basis set of $N$ modes and is not restricted to the subspace spanned by the $k$ modes selected.

The matrix **G** is generally not exactly symmetric, showing a correlation of ~ 0.99 between the upper and lower triangular entries (see Fig. S10a). We assigned no biological relevance to this negligible asymmetry and focused exclusively on undirected representations of connectomes, as per prior work (*25, 28, 57*). We thus symmetrized $G_{ij}$ by taking the average of its lower and upper triangular entries. The quantity $G_{ij}$ corresponds to the connection weight between locations $i$ and $j$ in our model. Accordingly, only positive weights are retained and negative weights are set to zero (see Fig. S10b for a visualization of the proportion of positive/negative entries). By construction, the GEM enhances the connection strength between pairs of cortical locations according to the expression profile of each mode $m$, given by the factor $(1 + r_s^2 \lambda_m)^{-1}$ in Equation S20.

The GEM in Equation S20 has two free parameters: the spatial scale of wave propagation, $r_s$ (mm), and the number of geometric modes, $k$. The specific values of these parameters were fitted to the data using an objective function that quantifies the similarity between model and empirical connectomes with respect to key elements of connectome topology and topography, as detailed in section 0.1.8.

For the human connectomes, we generated models for 50 values of $r_s$ equally spaced in $r_s = [1, 20]$ mm. In a preliminary search of the parameter space for values of $r_s$ between 1 and 80 mm (*53, 58*), we found that meaningful variations in fit statistics only resided between values of 1 and 20 mm. We performed the same parameter search for every other species, and found that optimal values spanned the ranges $r_s = [0, 3]$, $[0, 5]$, $[0, 9]$, and $[0, 15]$ mm for the mouse, marmoset, macaque, and chimpanzee, respectively.

For each value of $r_s$, we generated connectome models for $1 \leq k \leq 200$, representing a total of 10,000 sampled parameter combinations. Our connectome model can be computed via matrix multiplications with Equation S21, which considerably speeds up the connectome generation process compared to classical graph-based growth models, such as the matching index model, in which edges are added probabilistically one a time, with probabilities updating at each iteration (*28*) (see section 0.1.5). This computational efficiency allows us to apply the GEM to high-resolution networks.

##### 0.1.5 Benchmark models

We compared the performance of the GEM to several benchmark models. For the high-resolution connectome analysis, the benchmark models were 'LBO', 'Permuted', 'EDR-vertex', and 'Random'. These models were used to assess the specific contributions of the geometric and other constraints implied by NFT that are embedded within the GEM. For the atlas-based connectomes, the benchmarks were the matching index ('MI'), 'Distance-atlas', 'EDR-vertex', and 'Random'. In the analysis of atlas-based connectomes, the EDR-vertex and Random benchmarks were defined at vertex resolution and then parcellated using the same atlas as the empirical data according to the procedure outlined in section 0.1.6, thus indexing the level of performance induced simply by parcellating mesh-resolution connectivity matrices. The MI and Distance-atlas models were defined at atlas resolution and used as points of reference for state-of-the-art models widely used in the literature. The following presents the details and rationale for each model.

**Random**

Our simplest benchmark model involved forming connections at random according to an Erdős–Rényi-like process (*59*) with a connection density chosen to match the empirical edge density. This model serves as a simple baseline for the high-resolution analysis and a quantification of the bias associated with parcellating high-resolution connectomes in the analysis of atlas-based networks. This bias arises because larger parcels are likely to have more connections than smaller ones simply because they have more vertices, which may artificially inflate connection weights and nodal binary/weighted degrees.

We defined random networks with exactly the same number of connections as the empirical data, with all possible pairs of nodes having a uniform probability of connection. To have a 'random' benchmark for weighted network features, we also assigned weights sampled from a uniform distribution spanning the range $[0.01, 1]$. We assigned weights in the Random model only for the analysis of the high-resolution human connectome (Figs. 2 and S5). When analyzing atlas-based connectomes, the parcellation process already gives an estimate of connection weights, which captures biases imparted on the connectivity of parcels due to their surface areas and irregularities in the mesh (*60*).

The only parameter of the Random model is the number of edges, which is matched exactly to the data. As such, this model did not require any optimization process. We generated 100 realizations of the model and reported the distribution of model fit statistics across iterations.

**EDR-vertex**

The Random model is a simple yet unrealistic benchmark, given the well-known distance-dependence of cortico-cortical connectivity (*61–64*). We therefore evaluated a second benchmark that relied on a stochastic process constrained by an exponential distance rule (EDR). Under this model, the probability of connecting two vertices, $p_{ij}$, given their Euclidean distance $d_{ij}$ is

$$p_{ij} = \exp(-\eta_p d_{ij}), \tag{S22}$$

where $\eta_p$ controls how fast the connection probability decays with distance and is a free parameter fitted to the data. Binary connections are first established probabilistically, after which EDR connection weights are also added deterministically with $w_{ij} = \exp(-\eta_w d_{ij})$, where $\eta_w$ represents how fast connection weights decays with distance and is also fitted to the data.

We instantiated an EDR benchmark model at the vertex resolution of the neocortical surface mesh, called 'EDR-vertex', such that all distances and connections were estimated between vertices. For the high-resolution human connectome, we first explored 100 decay values in $-1 \leq \eta_p \leq 2$ and 100 values in $-1 \leq \eta_w \leq 2$. We identified the regions where fit statistics were optimal, and then refined this parameter space for optimization to 100 equally spaced values for the decay constants in $0.4 \leq \eta_p \leq 0.7$ and 100 values in $-0.1 \leq \eta_w \leq 0.3$. This model served as a benchmark for the analysis of high-resolution connectomes and allowed us to determine the specific contributions of the geometrically constrained mechanisms implied by the GEM as distinct from a generic EDR connectivity kernel.

The EDR-vertex benchmark was also used in the analysis of atlas-based connectomes. In this case, the model was again defined at vertex resolution. We performed the same preliminary search of parameter space mentioned above and then refined this space to 100 values in $0.05 \leq \eta_p \leq 0.2$ and 100 values in $-0.1 \leq \eta_w \leq 0.1$ where meaningful variation of fit statistics were observed.

The resulting high-resolution model was then parcellated using a predetermined atlas (see section 0.1.6). This benchmark allowed us to determine the level of model performance expected simply from parcellating a high-resolution EDR model, which is important given that the GEM underwent the same parcellation procedure. The fitted parameter $\eta_w$ was $\approx$0 (uniform weights) for both high-resolution and atlas-based human connectomes, so we did not include this additional free parameter in the EDR-vertex model for other species (these models only included the parameter $\eta_p$), consistent with previous EDR models in the literature (*25, 28, 57*). For all non-human species, we used 10,000 equally spaced values in $0 \leq \eta_p \leq 15$. This parameter space was chosen since it covered all meaningful variations in fit statistics across all species, given that 10,000 parameters were explored in that range.

During the optimization, each model was generated 10 times and the average of their fit statistics (see section 0.1.8 for definitions) was used to identify optimal parameters in order to account for the inherent stochasticity of the model. We generated 100 realizations of the optimized model and reported the distribution of model fit statistics across iterations.

**LBO**

The GEM rests on specific assumptions about how geometry and other constraints implied by the NFT dynamical theory might sculpt connectome wiring. To assess the specificity of these effects, which are encapsulated by the denominator in Equation S20, we compared the GEM against an additional benchmark that accounts for geometry alone, independent of NFT. This benchmark, which we name 'LBO' (see Equation S5), was constructed as a low-rank approximation of the LBO

$$\mathrm{LBO}_{ij} = \sum_{m=1}^{k} \lambda_m \psi_m(i) \psi_m^{+}(j). \tag{S23}$$

The model accounts for the short-range connectivity of the mesh itself and the geometric properties captured by the LBO, without the transformation of the eigenvalues via the Green's function derived from the NFT formalism (see Equation S18 and Ref. (*53*)). We used this benchmark to evaluate the predictive value of the GEM beyond low-order effects of geometry and mesh connectivity. The benchmark was only employed in the analysis of the high-resolution human connectome as the coarse-graining effect of a parcellation atlas obscures fine-scale differences between the GEM and LBO models. Note that the spatial patterns of the eigenmodes used in the LBO model and the GEM are identical; it is only their relative weighting that differ.

**Permuted**

As a second benchmark assessing the specificity of our GEM formulation, we defined a model to assess the importance of the specific ordering of the eigenvalues. For given optimal parameters $r_s$ and $k$, we specifically compared the GEM against 100 realizations of the same model, but with the order of the eigenvalues randomly permuted. This Permuted benchmark was generated only for the high-resolution human connectome for the same reason as the LBO benchmark.

**Matching Index (MI)**

The matching index (MI) model probabilistically adds binary connections one at a time, with connection probabilities being constantly updated during the process. The updates are determined by a rule that attempts to quantify a trade-off between a distance penalty on long-range connectivity and a bias towards connections that link nodes with similar topological neighbourhoods (i.e., the bias favours connections between nodes with links to similar sets of other nodes) (*25, 28, 57*). This rule is defined as

$$p_{ij} = (d_{ij})^{\eta} \times (t_{ij})^{\gamma}, \tag{S24}$$

where $p_{ij}$ is the probability of connection between node $i$ and node $j$ and $d_{ij}$ represents their Euclidean distance. The term $t_{ij}$ corresponds to the matching index, defined as

$$t_{ij} = \frac{\Gamma_{i\backslash j} \cap \Gamma_{j\backslash i}}{\Gamma_{i\backslash j} \cup \Gamma_{j\backslash i}}, \tag{S25}$$

where $\Gamma_i$ represents the set of nodes connected to $i$ and $\Gamma_{i\backslash j}$ is simply $\Gamma_i$ but with $j$ excluded from the set.

The free parameters $\eta$ and $\gamma$ control the putative trade-off between network wiring cost (operationalized in terms of edge lengths) and the topological complexity induced by forming connections with high MI. Prior work indicates that the MI model performs better than models that rely on the distance penalty alone, or models that substitute other topological properties for $t_{ij}$, in head-to-head comparisons (*25, 28, 57*). As the MI model can only generate binary networks, we only evaluated its performance in relation to binary properties and used a distinct objective function for parameter optimization (see section 0.1.8).

The computational burden imposed by the need to update wiring probabilities at each iteration means that the MI model is only practical for lower-resolution atlas-based connectomes (i.e, networks with fewer than around 300 nodes), which is a key limitation of the model. In the analysis of human atlas-based connectomes, we first explored 10,000 parameters combinations using a uniform grid bounded by $\eta = [-20, 0]$ and $\gamma = [-2, 2]$. After visualizing the similarity landscape (i.e., the landscape of model performance across the various parameter combinations), the range of parameters was further refined to 10,000 points uniformly spaced in a grid bounded by $\eta = [-11, -3]$ and $\gamma = [0.1, 0.7]$ where most meaningful variations in model performance exist. During optimization, the model was run 10 times for each parameter combination and the performance metrics obtained for these 10 runs were averaged to account for model stochasticity. For atlas-based connectomes in non-human species, the same initial parameter space was explored and then refined to 10,000 points contained in: $\eta = [-9, -2], \gamma = [-0.05, 0.8]$ for the mouse connectome, $\eta = [-10, -1.2], \gamma = [-0.5, 0.4]$ for the marmoset connectome, $\eta = [-5, -0.4], \gamma = [-0.4, 0.5]$ for the macaque connectome, and $\eta = [-15, -1.5], \gamma = [-0.5, 1]$ for the chimpanzee connectome. We generated 100 realizations of the optimized model and reported the distribution of model fit statistics across iterations.

**Distance-atlas**

The 'Distance-atlas' model was defined at the atlas resolution, such that all distances were estimated and connections were formed between the centroids of each parcellated region. This benchmark mimics the common implementation of the distant-dependent connectivity rule in the literature and was used as a point of comparison with past work (*25–28, 65*). For the atlas-based human connectome, we sampled 10,000 equally spaced values in $3 \leq \eta_p \leq 11$. For the mammalian connectomes, we sampled the following ranges: $-2 \leq \eta_p \leq 20$, $-2 \leq \eta_p \leq 10$, $-2 \leq \eta_p \leq 15$, and $-2 \leq \eta_p \leq 15$ for the chimpanzee, macaque, marmoset, and mouse, respectively. All these range limits for parameter optimization correspond to a refined space following a preliminary search in: $-5 \leq \eta_p \leq 20$, where the refined parameter space showed meaningful variation in fit statistics for each connectome.

Following prior work (*26, 28, 57*), we used the power law formulation (instead of an exponential as in Equation S22) where the probability of connection is given by

$$p_{ij} = (d_{ij})^{-\eta_p}. \tag{S26}$$

We included the Distance-atlas model because it has been widely used as a benchmark in prior cortical connectome modelling work (*25, 26, 28*). (Note that prior work has shown that model performance is not significantly affected by the choice of an exponential distance rule versus a power law formulation (*25*).)

By construction, this model can only be evaluated against atlas-based connectomes. The distances calculated using this approach are less accurate, and our findings indicate that the performance of the Distance-atlas model is therefore considerably lower than the performance of the

EDR-vertex model in capturing the properties of atlas-based connectomes (Figs. 3 and S22). Indeed, the EDR-vertex model often performed better than the MI model, suggesting that a more realistic, high-resolution EDR process which is then parcellated onto an atlas is sufficient to account for many aspects of connectome organization. Prior work evaluating distant-dependent processes defined purely in atlas space may thus underestimate the predictive power of this mechanism. During the optimization, each model was generated 10 times and the average of their similarity measures was used to identify optimal parameters in order to account for the inherent stochasticity of the model. We generated 100 realizations of the optimized model and reported the distribution of model fit statistics across iterations.

###### 0.1.6 Parcellating model connectomes

The GEM and other benchmarks (i.e., EDR-vertex, LBO, and Permuted) define connectomes at the intrinsic resolution of the surface mesh. The resulting high-resolution connectomes can then be parcellated with any arbitrary atlas to allow comparison to extant connectomes normally studied at coarse resolutions (*25, 26, 28, 57*). This procedure involves parcellating a high-resolution matrix $\mathbf{A}$ into a lower-resolution matrix $\mathbf{A_p}$ using a parcellation matrix $\mathbf{M}$, which has dimensions $p \times N$, where $p$ is the number of nodes (i.e., parcellated atlas regions) in the lower-resolution connectome and $N$ is the number of vertices in the high-resolution connectome, such that

$$\mathbf{A_p} = \mathbf{MAM}^T. \tag{S27}$$

The matrix $\mathbf{M}$ was constructed so that each vertex is assigned to one parcel and $M_{ij} = 1$ if vertex $i$ belongs to region $i$; otherwise, $M_{ij} = 0$. This matrix can be normalized to mitigate biases due to non-uniform parcel surface areas. In such cases, larger parcels will contain more vertices, which may artificially inflate connection weights and nodal degrees. A normalization is achieved by dividing each row of $\mathbf{M}$ by its sum, effectively dividing the connection weights $w_{uv}$ by the total number of vertices contained in parcels $u$ and $v$. We used this normalization step only for the three connectomes derived from tract-tracing studies (i.e., for mouse, marmoset, and macaque) to mirror the process used in generating the empirical data (see sections 0.1.2 and 0.1.7).

##### 0.1.7 Thresholding model connectomes

Each of the empirical connectomes was generated using different techniques that ultimately resulted in varying edge densities. Because the number of edges in a network can influence its properties, it is important to match the edge densities of model and empirical connectomes. For each empirical dataset, we applied a thresholding procedure to our models that was intended to mimic the data generation process, as detailed below.

**Humans**

The GEM and the high-resolution benchmarks (i.e., EDR-vertex, LBO, Permuted, and Random) were generated at the resolution of the fsLR-5k surface (i.e., 4,386 vertices, excluding the medial wall). When comparing with the high-resolution empirical connectome mapped onto the fsLR-5k mesh, we simply retained the top 4.6% of edge weights in each model and set the remaining entries to 0. The edge density of 4.6% corresponds to the average density of the 339 individual connectomes mapped at this resolution (see section 0.1.1).

When comparing these models with atlas-based connectomes, the same edge density threshold of 4.6% was first applied to the models generated on the fsLR-5k mesh. The thresholded high-resolution model networks were then parcellated using the Schaefer300 17-network atlas (corresponding to the 150-region atlas reported in the main text, since we only considered one hemisphere) (*17*) using Equation S27 and the procedure outlined in section 0.1.6. This parcellation of the high-resolution model networks invariably resulted in a larger number of edges than the empirical atlas-based connectome. We therefore applied a second threshold to the parcellated model connectomes, retaining the same number of strongest edges present in the empirical data. In the main text, this density was fixed to 10% for the Schaefer300 atlas-based human connectome (Fig. 3). We additionally show model performance for a range of connectome densities between 5% and 35% in Fig. S20. In Fig. S21, we show the performances of the GEM, EDR-vertex, and Random models for different pre-parcellation edge densities, ranging between 5% and 20%. We also evaluated the GEM's performance on different Schaefer atlas resolutions, which subdivide a hemisphere into 50, 100, 200, and 250 regions in Fig. 3d, where the empirical edge densities were set to 15%, 12.5%, 7.5%, and 5%, respectively.

**Chimpanzee**

As in humans, the chimpanzee neocortical surface was initially mapped onto the fsLR-32k space. We then downsampled it to fsLR-5k (4,842 vertices) using the same approach used for the human neocortical surface (see section 0.1.1). The chimpanzee connectome derived from diffusion MRI has full brain coverage and is mapped between 55 nodes, corresponding to a subset of 4,374 vertices. Before parcellating the high-resolution models (i.e., the GEM, EDR-vertex, and Random), we used an edge density threshold of 5%. This threshold was chosen to be close to that used in humans, given that both connectomes were derived from diffusion MRI. We also evaluated the GEM's performance for a range of values between 4% and 10% edge density (Fig. S40). The MI and Distance-atlas models were generated at the atlas resolution using the Euclidean distances between parcel centroids. We generated the atlas-based models with the same number of binary connections present in the empirical connectomes.

**Macaque**

The macaque cortical midthickness surface was modeled using the fsLR-10k template (*66*) and comprised 10,242 vertices, of which 4,137 were associated with the partial empirical connectome spanning 29 nodes (*35*). Since the atlas-based connectome does not have full hemispheric coverage, we modified the GEM, since it intrinsically produces connectomes that span all vertices of the neocortical surface mesh. Specifically, we computed the pseudo-inverse of $\mathbf{\Psi}_k$ and selected the row indices of $\mathbf{\Psi}_k$ and $\mathbf{\Psi}_k^+$ that correspond to the 4,137 vertex indices contained within the partial connectome. This operation is equivalent to constructing the full connectivity matrix and then subsampling its rows and columns, but our approach limits computational cost by reducing the matrix before multiplication. Before parcellating the high-resolution models, we used an edge density threshold of 60% when analyzing both the dense and sparse macaque connectomes. For the GEM, this threshold had no impact on the performance; it was used only to control the high-resolution edge density of the EDR-vertex and Random models. As a result, we did not perform additional analysis using different high-resolution edge densities for the GEM.

To make the weights in our model comparable with those used to estimate connectivity weights in tracer-derived empirical connectomes, we normalized the rows of the parcellation matrix $\mathbf{M}$ in Equation S27 (see section 0.1.6), i.e., we divided each row by its sum, so that every row sums to 1. This effectively scales the total connection weight between two parcels by the number of vertices they contain. Such normalization of the parcellation matrix better reflects how connection weights are defined by the FLNe in the macaque and marmoset connectomes, where connection weights are normalized in a similar way (see section 0.1.2). For the MI and Distance-atlas models, we used the same approach described for the chimpanzee.

**Marmoset**

The marmoset neocortical surface mesh comprised 37,974 vertices, of which 21,250 were associated with the partial empirical connectome spanning 55 nodes (*36, 67*). Before parcellating the high-resolution models, we applied an edge density threshold of 3%. While this threshold is relatively stringent, it is consistent with the sparsity of the atlas-based empirical connectome, which has an edge density of 16%. We also show the GEM's performance for high-resolution model edge densities between 3% and 15% (see Fig. S39). To construct the connectome models, we applied the same approach as described for the macaque to account for the partial coverage of the neocortical surface. We also normalized the parcellation matrix **M**, since the empirical connections weights were also estimated as the FLNe (*36*). For the MI and Distance-atlas models, we used the same approach described for the chimpanzee.

**Mouse**

Geometric modes of the right hemisphere of the mouse isocortex were computed using volumetric data from (*44*). Note that the voxels allotted to the right isocortex differ from those of other releases (such as (*45*)) due to the symmetrization procedure used by (*44*). We parameterized this isocortical volume was using a tetrahedral mesh comprising 7,364 vertices. The faces of the tetrahedral mesh were given by the volumetric alpha shape with the smallest alpha radius that enclosed all points.

The GEM was run using volumetric geometric eigenmodes computed using the same approach as in (*49*). Before parcellating the high-resolution models, we used an edge density threshold of 60% when comparing with the dense empirical connectome and 20% when comparing with the sparse connectome (see section 0.1.2 for details). We also evaluated the GEM's performance using different values between 20% and 100% for the GEM's edge density (see Fig. S38). As in the macaque and marmoset, we normalized the parcellation matrix **M**, since empirical connections weights were quantified using the normalized connection density (NCD) (see section 0.1.2). For the MI and Distance-atlas models, we used the same approach described for the chimpanzee.

##### 0.1.8 Objective functions

The free parameters for each model were fitted to the empirical data to maximize an objective function quantifying similarity between model and data with respect to several key topological and topographical properties. These properties were computed using different measures of undirected graph-based representations of model and data connectomes, with the graphs comprising $N$ nodes interconnected by $E$ undirected edges. If a network is binary, $A_{ij} = A_{ji} = 1$ if a connection exists between $i$ and $j$, and $A_{ij} = 0$ otherwise. For a weighted network, the strength of a connection is denoted as $w_{ij}$, where $w_{ij} \geq 0$. We used two different objective functions depending on the models being compared.

In the analysis of high-resolution models (i.e., the GEM, LBO, and EDR-vertex), we used an objective function defined as the sum of three rank correlations, $\rho$: (1) the correlation between model and empirical edge weights, taken over the union of edges (hereafter referred to as edge weights union); (2) the correlation between model and empirical nodal degree sequences; and (3) the correlation between model and empirical rank-based nodal strength sequences. We used rank correlations as they provide a nonparametric measure that assesses the linear monotonic relationship between two vectors without assuming any specific underlying distribution of values, and because our model is largely intended to reproduce the rank ordering of network properties without capturing their specific values (see below).

The 'edge weights union' measure corresponds to the Spearman rank correlation of the union of all the edge weights that exist (i.e., $w_{ij} > 0$) in *either* the model or the empirical connectome. By considering the union rather than only the intersection, this metric provides a more comprehensive assessment of how well the model captures the relative strength of connections and also of its ability to predict the empirical binary network topology (i.e., the presence/absence of specific edges).

The nodal degree correlation was computed using the binary node degree of each node, $k_i$, which corresponds to the number of binary connections incident upon it

$$k_i = \sum_j A_{ij}, \tag{S28}$$

where the sum runs over all column indices $j$. The resulting map of degree values encodes the extent to which each network node is connected to all others. The resulting map is an important topographical property of empirical connectomes that has thus far been difficult to capture with extant generative network models (*25, 26*).

The rank-based strength correlation was computed using a weighted variant of node degree, also called node strength, $s_i$, which corresponds to the sum of all edge weights attached to each node

$$s_i = \sum_j w_{ij}. \tag{S29}$$

A challenge arises when comparing empirical node strength maps with those derived from models. The empirical range of edge weights can span up to six orders of magnitude but the product of pairwise location-specific geometric modes amplitude values used to define the edge weights in the GEM (see Equation S20) is not defined to be proportional to empirical edge weights; rather, it is intended to capture the *rank ordering* of empirical edge weights. We therefore defined an alternative strength measure called the rank-based strength $s_i^{\mathrm{r}}$, in which we replace the actual edge weights $w_{ij}$ with their ranks, assigning values such that the weighted network has edge weights in the range $1 \le w_{ij}^r \le E$, where $E$ is the total number of edges. The rank-based strength is thus calculated as

$$s_i^{\mathrm{r}} = \sum_j w_{ij}^r. \tag{S30}$$

Figures S6, S23, S41, S42, S43, and S44 show that there is generally good agreement between Equations S29 and S30, along with the nodal strength obtained after resampling the edge weights according to the method proposed in Ref. (*68*), which is an alternative approach for mitigating the wide range of edge weights encountered empirically. The two exceptions are the high-resolution human and atlas-based macaque connectomes, where original and rank-based strength maps diverge, but weight resampling increases their convergence suggesting that the raw strength measure, $s_i$, is highly susceptible to outlying weight values. Together, these results indicate that, as intended, that GEM accurately captures the rank ordering of edge weights, but does not necessarily reproduce the full range of edge weight values.

Given these considerations, we defined our weighted objective function as

$$\arg\max(\rho_{\text{degree}} + \rho_{\text{rank-based strength}} + \rho_{\text{edge weights union}}). \tag{S31}$$

This objective function was used to optimize the GEM, LBO, and EDR-vertex in all analyses, unless otherwise specified (see below). When evaluating models on the dense mouse and macaque connectomes in Figs. 4, S29, and S30, we focused exclusively on weighted properties and therefore excluded the binary degree from the objective function, since binary properties carry little information in dense networks. In these instances, the objective function was

$$\arg\max(\rho_{\text{rank-based strength}} + \rho_{\text{edge weights union}}). \tag{S32}$$

The MI and Distance-atlas models can only generate binary networks. Hence, the objective function in Equation S31 is inappropriate. For these binary models, we defined a different objective function that relied only on two binary properties: the binary degree correlation and the edge-wise true positive rate, defined as the percentage of empirical edges recovered by the model:

$$\arg\max(\rho_{\text{degree}} + \text{edge-wise true positive rate}). \tag{S33}$$

The edge-wise true positive rate offers an intuitive assessment of the model's ability to capture the presence of connections (i.e., the binary topology) in the data. As such, it was also used to evaluate the performance of models optimized with Equation S31, even though it was not directly embedded within the objective function. We additionally examined the edge-wise true positive rate within different connection length ranges, defined using 10 bins comprising the same number of connections, across all connectomes analyzed (Figs. S7, S19, and S45).

Figures 3, S27, S26, S25, and S24 compare the GEM and EDR-vertex models optimized with Equation S31 and the MI and Distance-atlas models optimized with Equation S33. In Fig. S16, we provide comparisons between the MI, Distance-atlas, and other models on the atlas-based human connectome, in which all models were optimized using the two-property binary objective function in Equation S33.

Previous studies have fitted models to data using only topological properties, relying on the Kolmogorov-Smirnov (KS) statistic to quantify the similarity between nodal or edge distributions of properties such as node degree, clustering, betweenness centrality, and connection distance (*25, 28, 57*). However, the KS statistic is agnostic to topographical properties; as such, models optimized using this statistic generally fail to capture the way in which topological features are spatially embedded within the brain (*25, 26*). Recent analyses have documented further limitations with KS-based optimization (*69*). We therefore used the specific form defined in Equation S31 to capture both binary and weighted aspects of network topology and topography.

###### 0.1.9 Other metrics for model evaluation

To determine model generalizability, it is important to assess performance in capturing network properties for which parameter values were not directly optimized in the objective function. We therefore evaluated several additional network properties capturing distinct aspects of weighted and binary topology and topography (*24, 70*). The network features were extracted from both model and empirical connectomes and, unless otherwise specified, the similarity between model and data was quantified using Spearman rank correlations ($\rho$) across either nodes or edges, as appropriate.

**Edge intersection weight correlation**

This measure corresponds to the Spearman rank correlation between edge weights that simultaneously exist in both the model and empirical connectomes (i.e., $w_{ij} > 0$); i.e., it is the rank correlation computed over the intersection of non-zero edge weights found in model and empirical data. This metric was not considered during model optimization, but was used in scatter plots showing the correspondence between model and empirical edge weights.

**Average node connection distance**

The average connection distance of a node corresponds to the average length of all connections incident on that node,

$$\ell_i = \sum_j \frac{\sum_j A_{ij} d_{ij}}{\sum_j A_{ij}}, \tag{S34}$$

where $d_{ij}$ is the Euclidean distance between nodes $i$ and $j$. These distances represent a key aspect of the way in which the connectomes are spatially embedded and their recapitulation is an important goal for connectome models.

**Clustering coefficient**

The clustering coefficient of a node $i$ quantifies the probability that any two of its neighbours are also connected to each other. While random networks typically exhibit low clustering, complex networks such as brain networks tend to display high clustering. This property has been linked to key functional advantages, including support for locally specialized processing and increased robustness to failure (*24, 70*). It is a widely used metric for characterizing network architecture and is defined as

$$c_i = \frac{T_i}{k_i(k_i - 1)/2}\,, \tag{S35}$$

where $T_i$ is the number of triangles attached to node $i$. The value of $c_i$ is set to 0 when $k_i < 2$ (i.e., when node $i$ has less than two connections). The resulting spatial map of nodal clustering coefficients captures the spatial embedding of this important topological property.

**Modularity (community structure)**

Modular organization is a fundamental feature of empirical brain networks (*71–81*) and is characterized by groups of nodes forming densely interconnected communities (or modules). More specifically, a network with high modularity has dense connections within its communities and sparse connections between communities. These modules are thought to support specialized cognitive functions by enabling efficient local processing through dense intra-community connections, while supporting global integration via sparse inter-community links (*82, 83*). The modularity $Q$ of a network is quantified as (*84*)

$$Q = \frac{1}{2E}\sum_{ij}\left(A_{ij} - \frac{k_i k_j}{2m}\right)\delta(m_i, m_j)\,, \tag{S36}$$

where $E$ is the total number of edges in the network, $\delta(m_i, m_j) = 1$ if $i$ and $j$ belong to the same community, and 0 otherwise, and $k_i$ represents the binary degree of node $i$ introduced earlier.

The problem of finding an optimal partition to maximize $Q$ can be approached using many techniques (*84–87*). For our purpose, we wanted to systematically compare the optimal partitions of the model and empirical connectomes across different numbers of communities. To do so, we performed a $k$-means clustering of the $K$ leading eigenvectors of the modularity matrix $\mathbf{H}$, where the number of communities is equal to the number of leading eigenvectors selected (i.e., the eigenvectors associated with the largest positive eigenvalues) (*87–89*).

The expression for the modularity matrix is given by

$$H_{ij} = A_{ij} - \frac{k_i k_j}{2m}. \tag{S37}$$

We used the normalized variation of information (NVI), an information-theoretic distance measure that quantifies the similarity between the model and empirical partitions, where smaller values indicate a better match with the empirical partition (*82, 90, 91*). This measure thus allows us to quantify the similarity between model and data in the spatial topography of their community structures. For the high-resolution and atlas-based human connectomes, we compared community structures across different numbers of communities ranging from 3 to 15 (Figs. S3 and S17) and computed the area under the curve (AUC) for each model.

For the non-human species, the range of the numbers of communities was fixed from 3 to 10, given the smaller number of nodes of these connectomes (Figs. S31, S32, S33, and S34).

**Spectral distance**

The spectral distance between the model and empirical connectomes was computed as the cosine similarity between the eigenspectra of their respective normalized graph Laplacian (*92–95*). This measure is purely topological since the eigenspectrum of a graph is independent of the way in which the nodes are labeled. To allow comparison with extant binary models, we considered the normalized graph Laplacian of the binary network representation of both model and empirical connectomes.

The eigenspectrum of the graph Laplacian captures diverse aspects of the connectome's topology such as the degree distribution, community structure, path lengths, motifs architecture, and dynamic interactions between the constituents of the network (*93–97*). The spectrum thus provides a succinct summary of multiple features of network topology, and has been used to compare the connectomes of different species (*96, 98*) and for classifying distinct classes of networks (*92, 99, 100*).

The normalized graph Laplacian matrix $\mathbf{L}$ is defined as

$$\mathbf{L} = \mathbf{I} - \mathbf{D}^{-1}\mathbf{A}, \tag{S38}$$

where $\mathbf{I}$ is the identity matrix, $\mathbf{D}$ is the diagonal matrix of the binary degree of each node, and $\mathbf{A}$ is the binary adjacency matrix (i.e., the binary connectome). The entries of the graph Laplacian matrix are thus given by

$$L_{ij} = \begin{cases} 1 & \text{if } i = j \text{ and } k_i \neq 0 \\ -\frac{1}{k_i} & \text{if } i \neq j \text{ and } i \text{ is connected to } j \\ 0 & \text{otherwise.} \end{cases} \tag{S39}$$

We took the absolute value of the eigenspectra of $\mathbf{L}$ calculated for both model and empirical connectomes and quantified their distance using the cosine similarity (see Figs. S4 and S17 for high-resolution and parcellated results in humans, respectively, and Fig. S35 for non-human species). This approach effectively treats each network as a low-dimensional embedding, represented as a feature vector (i.e., the eigenspectrum). However, for partial connectomes with a small number of nodes (such as the macaque with only 29 nodes), the extent to which this spectral representation captures meaningful topological properties becomes less certain, potentially limiting its interpretability and sensitivity. For the mouse and macaque connectomes, we used the sparse connectomes (see section 0.1.2) to allow comparison with extant binary models (i.e., the MI and Distance-atlas).

**Row-wise Pearson correlations versus distance**

We obtained a weighted metric of edge reconstruction versus distance by adapting the similarity metric previously used in Ref. (*13, 15*). The similarity between two networks was calculated as follows: (1) we computed the row-wise Pearson correlation between weighted model and empirical connectomes, yielding $N$ correlations indicating the similarity between each node of the network; and (2) we averaged these correlations resulting in a single similarity measure between the two networks. We repeated this procedure across a series of connection-length bins, where, for each bin, only edges whose lengths fell within that range were included, and other connection weights were set to 0. This metric thus defines the success of each model in capturing edge weights variations within distinct distance bins and can only be calculated for weighted network models (i.e., the GEM, LBO, Permuted, EDR-vertex, and Random) (Figs. S7, S19, and S45).

##### 0.1.10 Spin tests

To assess the statistical significance of the rank correlations between the GEM and empirical nodal and edge properties (i.e., degree, rank-based strength, average node connection distance, clustering, edge weights, and edge true positive rate) we compared each observed correlation against a null distribution of 10,000 values generated using spatially constrained randomization methods. For the surface-based connectomes, i.e., human, chimpanzee, macaque, and marmoset), we used the Spin test (*101, 102*), which generates surrogate maps by applying random spatial rotations to the vertices (or parcel centroids) on a spherical surface, thereby preserving the spatial autocorrelation between vertices/parcels. The null correlation values were computed as the rank correlation between the surrogate maps and the empirical map. For the high-resolution human data, we applied the method introduced by Ref. (*101*), while for the atlas-based connectomes, we used the approach proposed by Ref. (*102*), following recommendations outlined in Ref. (*103*). We generated the surrogate maps using the Python module neuromaps (*104*). For the mouse data, which is mapped in volumetric space, we generated the surrogate maps using the BrainSMASH approach proposed by Ref. (*105*). The one-sided $P$ values, $P_{\mathrm{spin}}$ (surface-based surrogate maps) and $P_{\mathrm{SMASH}}$ (volume-based surrogate maps), were computed as the number of null correlation values greater or equal to the empirical correlation value.

For the high-resolution human connectome, we handled medial wall vertices following the approach of Refs. (*101, 102*); i.e., for each rotation of the vertex indices, we masked the union of the original medial wall vertices and any medial vertices that were rotated into the cortex, in both the model, empirical, and surrogate maps. Because the number of medial wall vertices rotated into the cortex varied across rotations, the resulting 10,000 comparisons contained maps of different sizes. To minimize this variability, we first generated 20,000 rotations and then selected the 10,000 that had the fewest medial wall vertices leaking into the cortex. Finally, $P_{\mathrm{spin}}$ was estimated as the proportion of null correlation values greater than or equal to the empirical correlation value (with the appropriate indices masked), using pairwise comparisons between the empirical and null correlations. This process therefore yielded a distribution of empirical correlations, resulting from different vertex indices being masked for each rotation. We opted for this pairwise comparison across rotations to guarantee that both empirical and null correlations were derived from maps of identical size, thereby ensuring a fair comparison (see Fig. S13 for the distribution of correlations obtained with this procedure on the smoothed and non-smoothed connectomes).

###### 0.1.11 Cross-validation

To avoid over-fitting and enable fair comparison between models with different numbers of parameters, we performed split-half cross-validation with 10 iterations for the analysis of human data (Figs. 2c and 3 on smoothed connectomes and Figs. S22c and S22c on non-smoothed connectomes). We randomly separated the 339 individual HCP connectomes into two (almost equal-sized) groups and generated a train and test group-average connectome from these two sub-groups. All models were first optimized on the average connectome of the training set and the optimal parameters were then used to evaluate model performance on the unseen test set.

The results in Figs. 2a, b, c, Figs. S5a, b, c, Figs. 3a, b, c, e, and Figs. S22a, b, c are for the analyses of the entire sample. The results presented in panel d of these figures show the cross-validated fit statistics. We found a strong agreement between both analyses, supporting the generalizability of the GEM.

#### 0.2 Supplementary Figures

##### 0.2.1 High-resolution human connectome

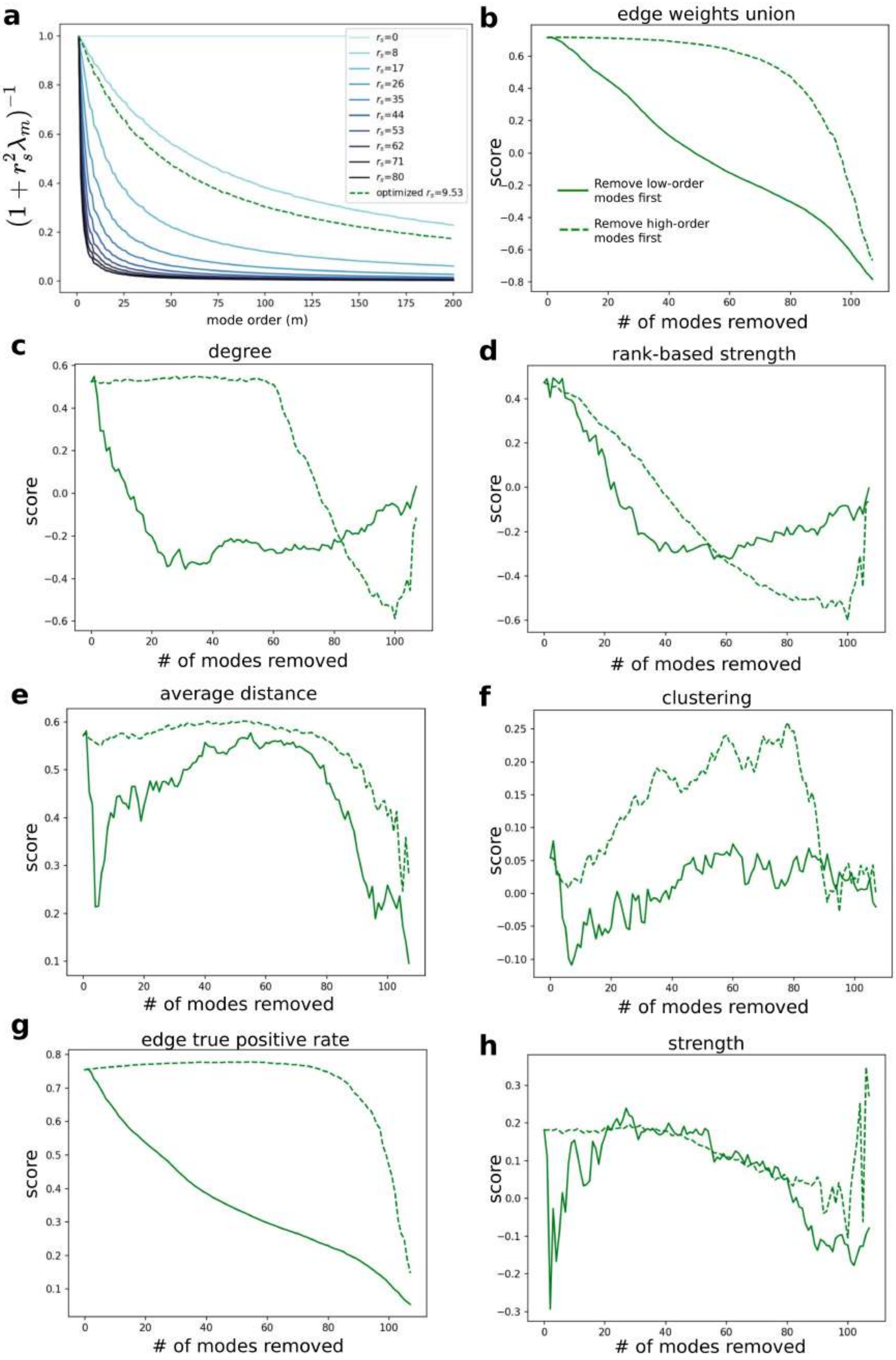
a
b
edge weights union
c
degree
d
rank-based strength
e
average distance
f
clustering
g
edge true positive rate
h
strength
mode order (m)
### of modes removed
score
Remove low-order modes first
Remove high-order modes first
optimized $r_s$=9.53

**Figure S1**: **Long-wavelength modes make a dominant contribution to the GEM's performance.** **a:** The dependence of the denominator term $(1+r_s^2\lambda_m)^{-1}$ in Equation 3 on mode order $m$ for various values of $r_s$, along with the optimal $r_s = 9.53$ mm found for the smoothed high-resolution human connectome. Higher values of $r_s$ lead to a sharper decline in the weighting that the GEM assigned to contributions from short-wavelength modes. **b:** Edge weights union **c:** binary nodal degree **d:** rank-based nodal strength **e:** average node connection distance **f:** nodal clustering **g:** edge true positive rate **h:** nodal strength. See sections 0.1.8 and 0.1.9 for details about each network metric. For panels b to h, the optimal parameters $r_s = 9.53$ mm and $k = 108$ modes were used as the starting point, i.e., corresponding to the case where no modes are removed. Except for the edge true positive rate, all scores in the remaining plots are rank correlations, $\rho$, between the GEM and empirical smoothed connectome as a function of the number of modes that have been removed from the optimal model. Solid and dashed lines correspond to the removal of modes starting from the first (long-wavelength) and last (short-wavelength) modes, respectively.

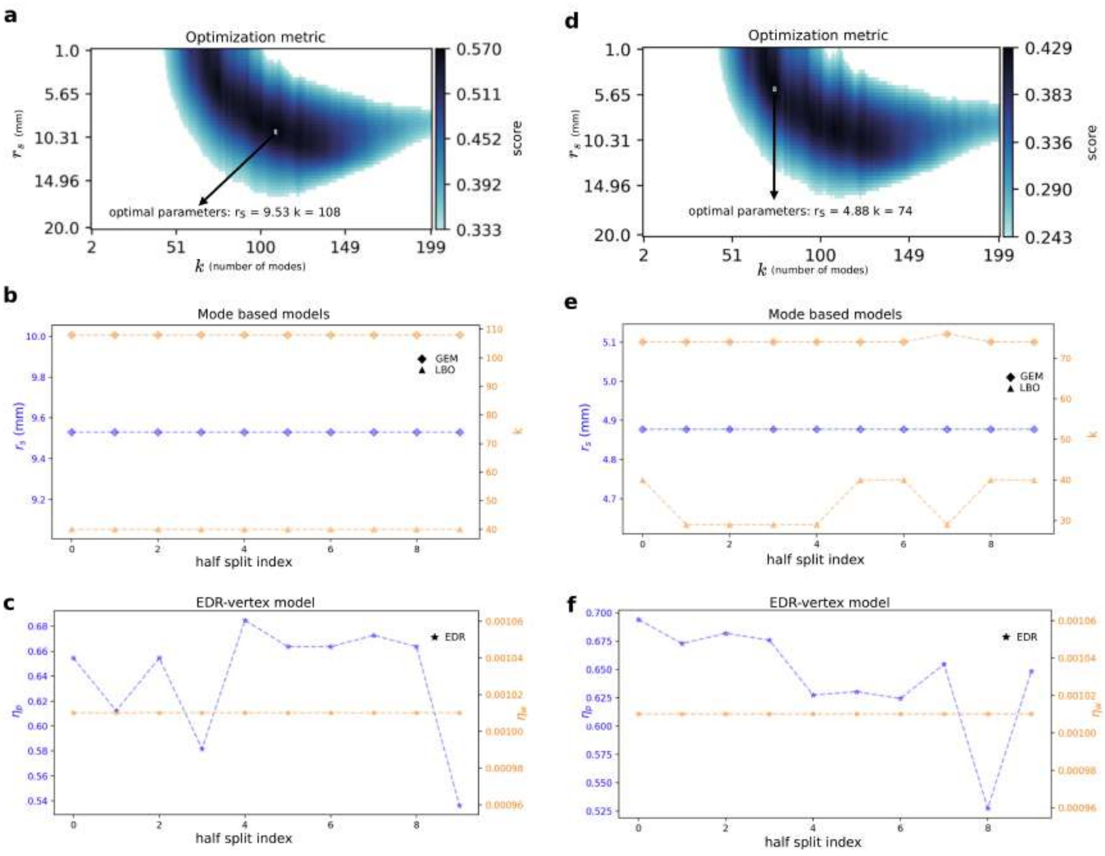


**Figure S2**: **Consistency of model parameters. a:** GEM optimization landscape using the full sample to generate the smoothed high-resolution human connectome. The GEM has two free parameters: the length scale parameter $r_s$ (mm) and the number of geometric modes $k$. The optimal parameter combination is $r_s = 9.53$ mm and $k = 108$ modes. The score represents the average of the three correlation metrics comprising our objective function, as defined in Equation S31. For a clearer visualization, the landscape shows only the scores among the top 60%, coloured from light blue to black. **b:** Optimal parameters for the GEM and LBO model for each half split cross-validation on the smoothed connectome. The LBO model has only one free parameter $k$. The Permuted model is not shown here because it uses the same optimal parameters as the GEM. **c:** Optimal decay constants of the EDR-vertex model for the probability of connection ($\eta_p$) and connection weight ($\eta_w$) for the smoothed connectome. Panels b and c show that the optimal GEM parameters are consistent across folds, whereas the EDR parameter shows much greater variation. **d:** Same as **a**, but using the non-smoothed connectome. The optimal parameter combination is $r_s = 4.88$ mm and $k = 74$ modes. **e** and **f** correspond to panels **b** and **c**, respectively, but using the non-smoothed connectome.

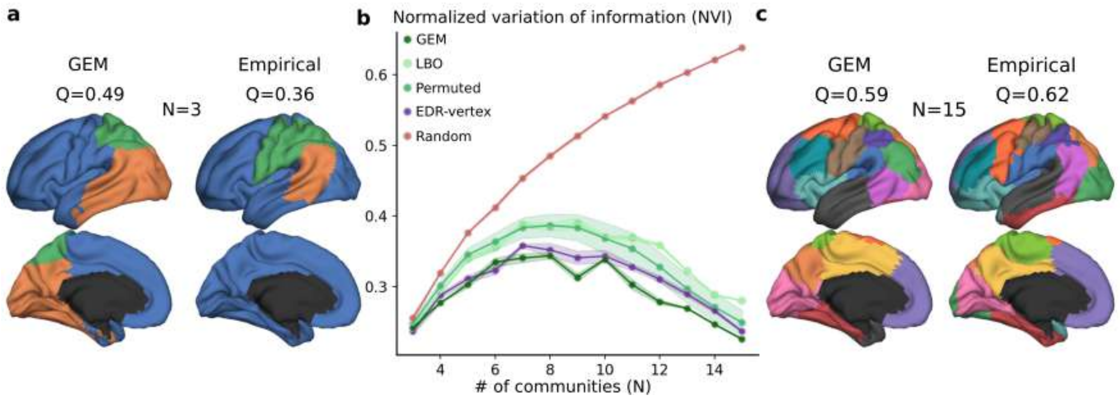


**Figure S3**: **The GEM captures the multiscale topography of the empirical human connectome's community structure better than benchmark models.** Using cross-validated fits, we quantified the agreement between the modular architecture of model and empirical smoothed high-resolution human connectomes at different community resolutions using the normalized variation of information (NVI; see section 0.1.9), for which lower values indicate better agreement. We calculated the NVI for a range of numbers of communities between $N = 3$ and $N = 15$. **a:** Visualization on the neocortical surface of the GEM and empirical communities for a solution comprising 3 communities, with the modularity $Q$. Each colour represents a set of vertices within a community, with each vertex assigned to a single community. **b:** NVI across different numbers of communities for every model using cross-validated fits: GEM (AUC = 3.58), LBO (AUC = 4.12), Permuted (AUC = 4.03), EDR-vertex (AUC = 3.75), and Random (AUC = 5.92), where AUC is the area under the curve, with lower values indicating greater similarity to the empirical community structure. The shaded areas represent the standard deviation for each model through cross-validated splits. **c:** Model and empirical communities for a solution comprising 15 communities, with the modularity $Q$.

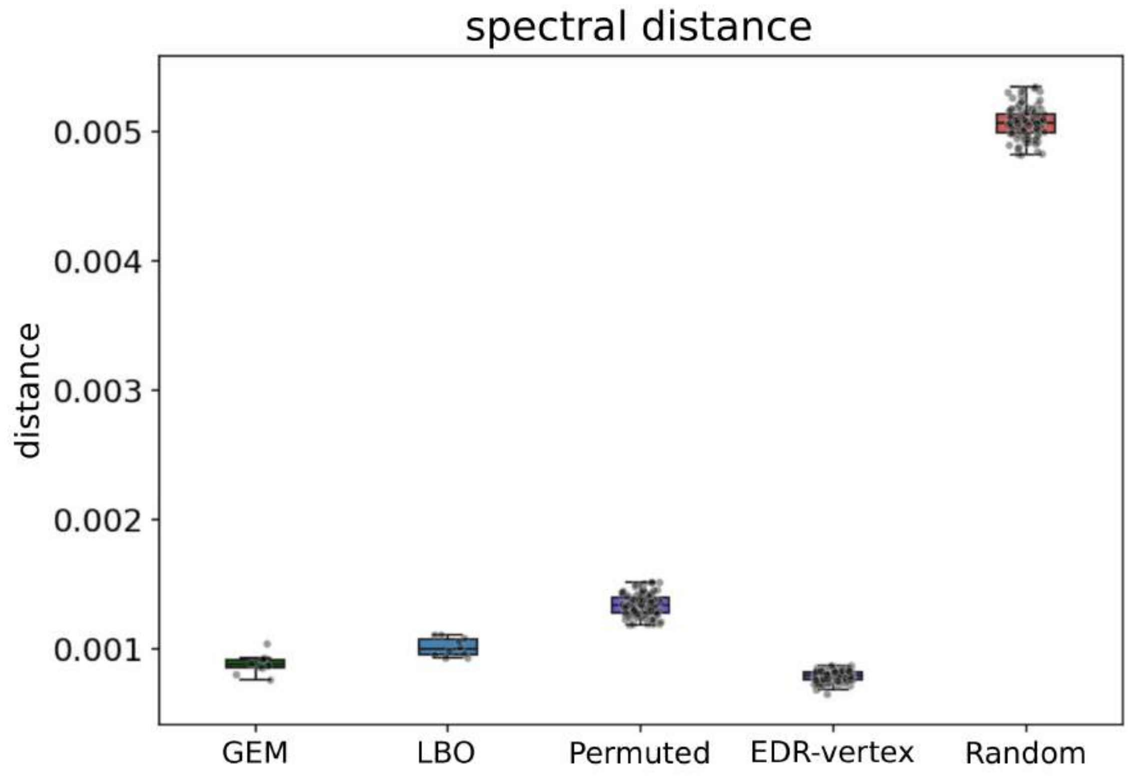


**Figure S4**: **Spectral distances between model and data for the smoothed high-resolution human connectome using cross-validated fits.** We assessed the topological similarity between the GEM and empirical connectomes using their spectral distance, with lower values indicating better agreement; see section 0.1.9.

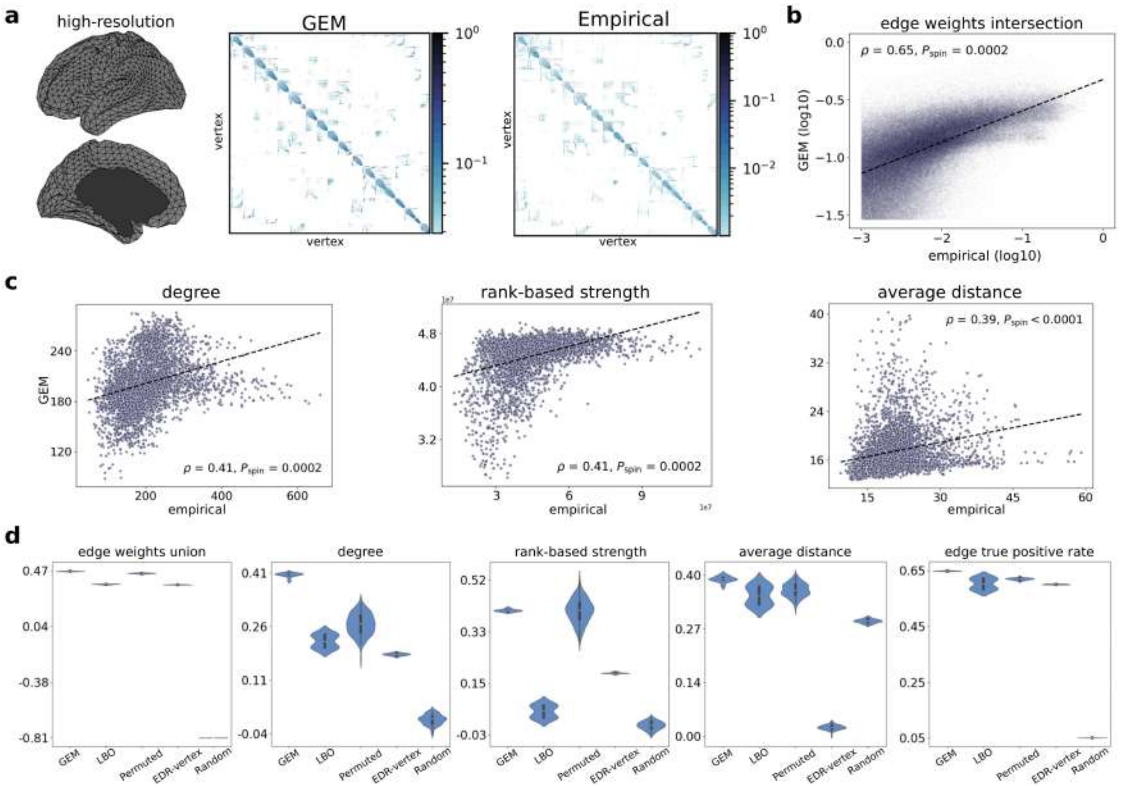


**Figure S5**: **Modelling high-resolution human connectomes.** Same as Fig. 2, but without smoothing the high-resolution empirical connectome; see section 0.1.1. **a:** Neocortical surface mesh and optimized GEM and empirical high-resolution connectivity matrices mapped with diffusion MRI. The optimized model was generated using parameters $r_s = 4.88$ mm and $k = 74$ modes, obtained by maximizing an objective function that quantifies similarities in binary and weighted topological and topographical properties of model and empirical connectomes (see Methods). **b:** Scatter plot of the common (intersection) edge weights between the GEM and empirical connectomes using the full dataset during optimization and testing. $P_{\text{spin}}$ is the one-sided p-value estimated from 10,000 spin permutations. **c:** Scatter plots showing the topographical agreement between the optimized model and empirical data using the full dataset, quantified using Spearman rank correlations ($\rho$). Panels from left to right show results for the degree (binary degree), rank-based strength (weighted degree), and average node connection distance. **d:** Comparison of the performance of the GEM relative to other models, as evaluated using split-half cross-validation. GEM=Geometric Eigenmode Model; LBO=Laplace-Beltrami Operator model approximating the connectivity of the mesh; Permuted=GEM computed with optimal parameters after randomizing the ordering of the geometric eigenvalues; EDR-vertex=Exponential Distance Rule model defined at the vertex level; Random=Random network defined at vertex resolution (see Methods for details).

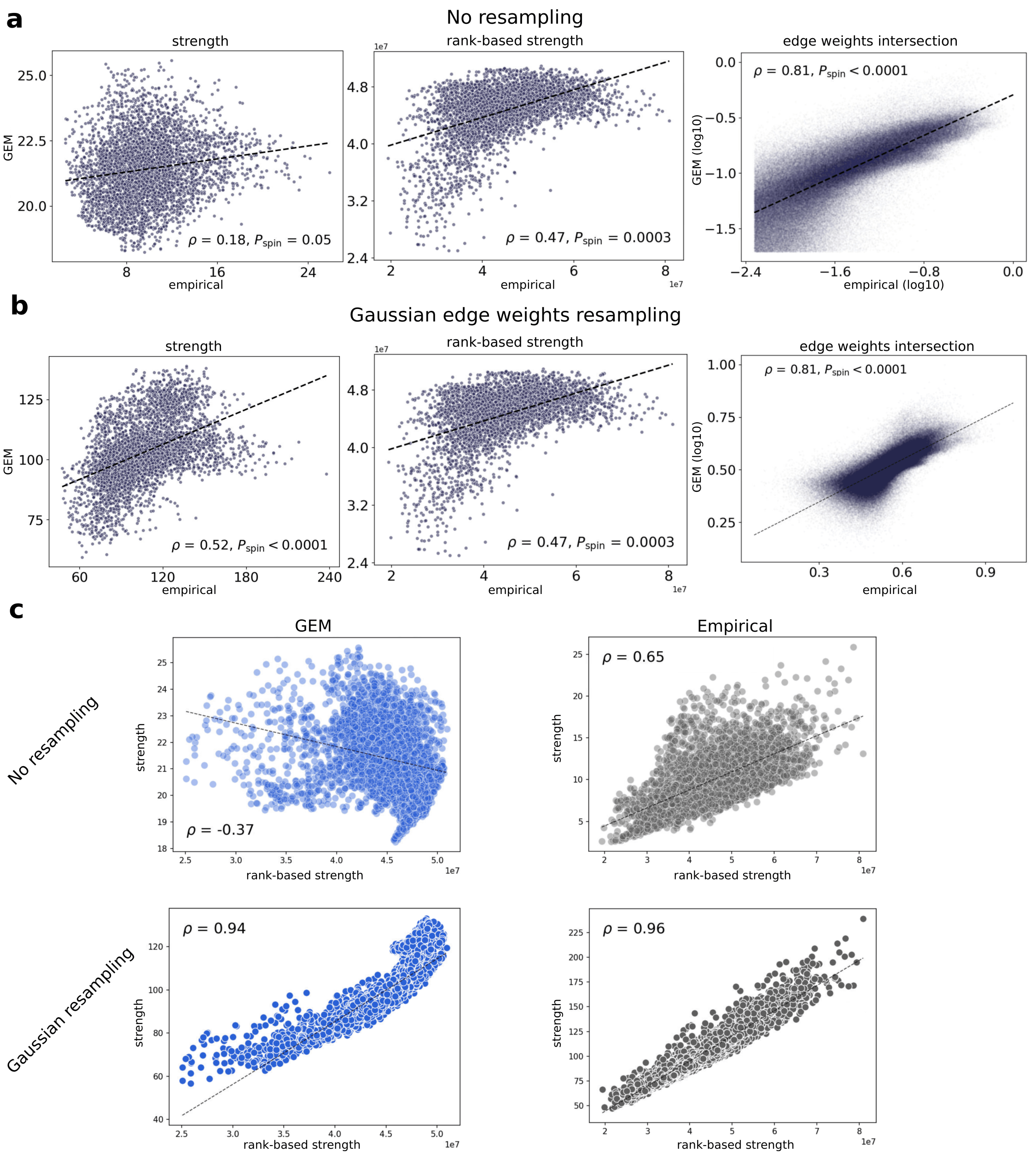
a
No resampling
strength
GEM
empirical
ρ = 0.18, P_spin = 0.05
rank-based strength
empirical
ρ = 0.47, P_spin = 0.0003
edge weights intersection
ρ = 0.81, P_spin < 0.0001
GEM (log10)
empirical (log10)
b
Gaussian edge weights resampling
strength
GEM
empirical
ρ = 0.52, P_spin < 0.0001
rank-based strength
empirical
ρ = 0.47, P_spin = 0.0003
edge weights intersection
ρ = 0.81, P_spin < 0.0001
GEM (log10)
empirical
c
GEM
Empirical
No resampling
strength
rank-based strength
ρ = -0.37
ρ = 0.65
Gaussian resampling
ρ = 0.94
ρ = 0.96

**Figure S6**: **Agreement between different definitions of nodal strengths and effect of edge weights resampling in the human connectome. a:** Panels from left to right show scatter plots of nodal strength, rank-based strength, and the edge weights intersection between GEM and smoothed high-resolution empirical human connectome. **b:** Same as **a**, but after resampling the edge weights. **c:** Scatter plot of node strength versus rank-based strength in the GEM (top left), node strength versus rank-based strength in the empirical connectome (top right), node strength versus rank-based strength after resampling the edge weights in the GEM (bottom left), and node strength versus rank-based strength after resampling the edge weights in the empirical connectome (bottom right). Here, we compared different definitions of nodal strength before and after resampling the edge weights using the approach in (*68*). Since the resampling of edge weights preserves their ranking, it has no effect on rank correlations at the edge level. We found that the two definitions of nodal strength given in Equations S29 and S30 differ in the GEM, whereas the two definitions show better agreement in the high-resolution empirical connectome, implying that smaller edge weights contribute more to the nodal strength in the GEM compared to the empirical connectome, resulting from the difference in magnitudes spanned by the edge weights in both cases. We also observed that resampling the edge weights increases the agreement between the two definitions.

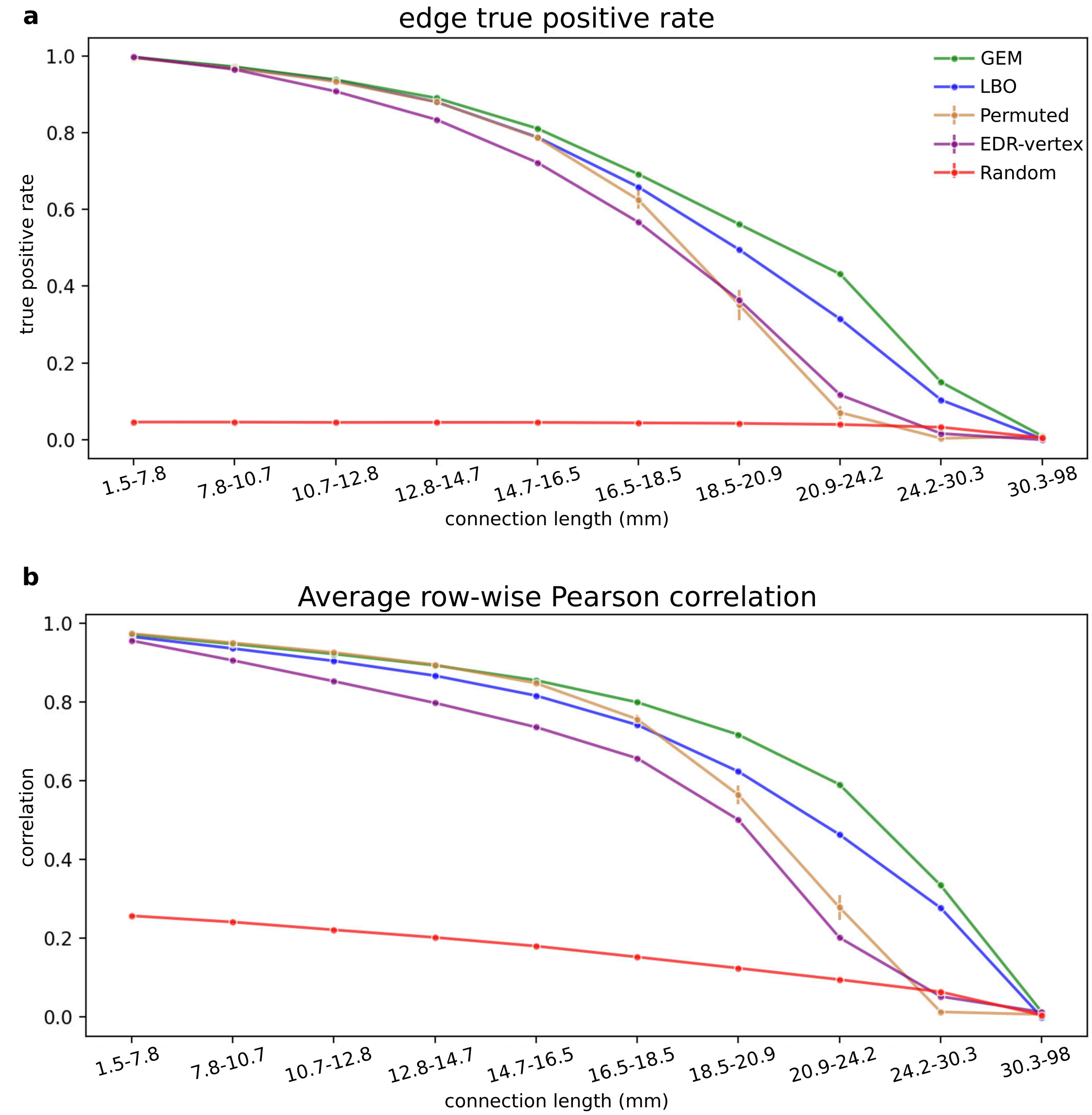


**Figure S7**: **The GEM better captures empirical edges than benchmark models across all connection lengths in the human connectome.** We assessed the GEM's ability to predict empirical edges in the high-resolution smoothed connectome across 10 connection-length ranges, each containing an equal number of edges. **a:** Edge-wise true positive rate (binary) as a function of connection length for the GEM, LBO, Permuted, EDR-vertex, and Random. **b:** Average row-wise Pearson correlation across connection lengths for the same models in **a**. To quantify the correlations, we adapted the approach of (*15*): for each length range, we excluded edges outside the range, computed row-wise Pearson correlations between the model and empirical connectomes (preserving the weights), and averaged these values to obtain a single similarity score per range of connection lengths.

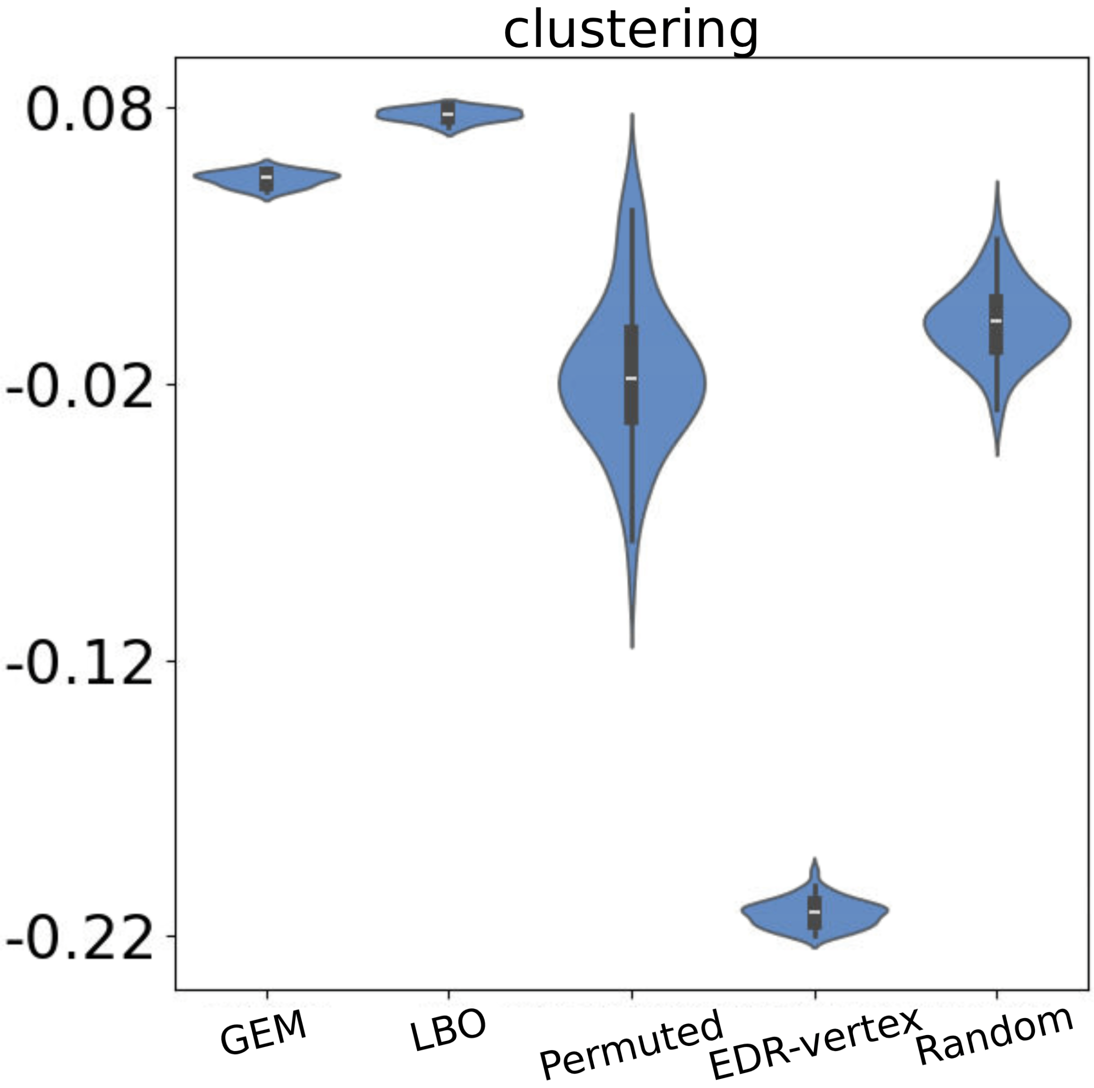


**Figure S8**: **High-resolution model and benchmarks fail to capture the spatial embedding of nodal clustering in the smoothed high-resolution human connectome.** Rank correlation $\rho$ between the empirical and model clustering in the GEM and benchmark models, i.e., LBO, Permuted, EDR-vertex, and Random, using cross-validated fits.

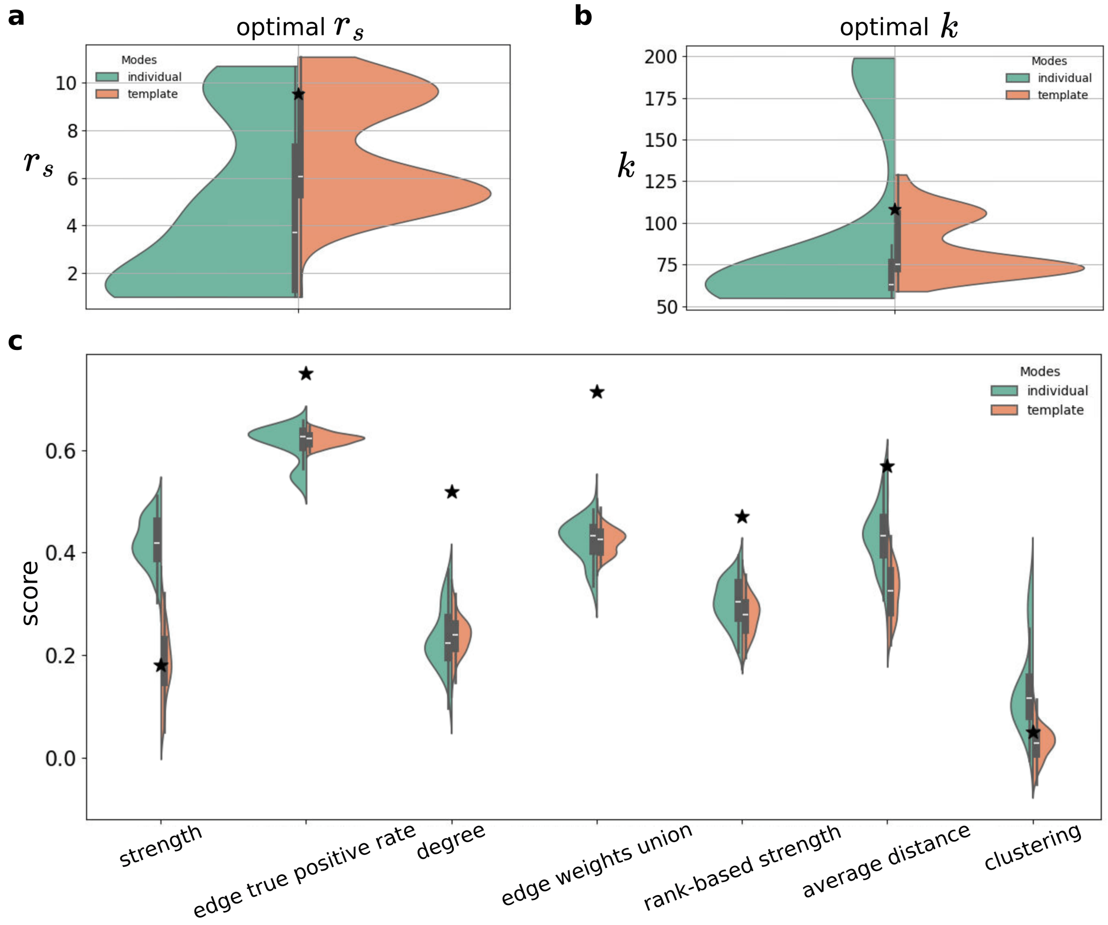


**Figure S9**: **GEM's performance on individual smoothed high-resolution connectomes from a subset of 100 participants.** For each participant, we optimized the GEM using (1) the modes derived from the group-average template neocortical surface (coloured orange); or (2) the modes derived from the individual's neocortical surface. In each plot, the black star corresponds to the performance obtained on the group-average connectome in Fig. 2 with optimal parameters identified in Fig. S2. **a:** Distribution of optimal length scale parameter $r_s$. **b:** Distribution of optimal number of modes $k$. **c:** GEM's performance across network metrics: from left to right, nodal strength, edge true positive rate, nodal (binary) degree, edge weights union, nodal rank-based strength, average node connection distance, and clustering.

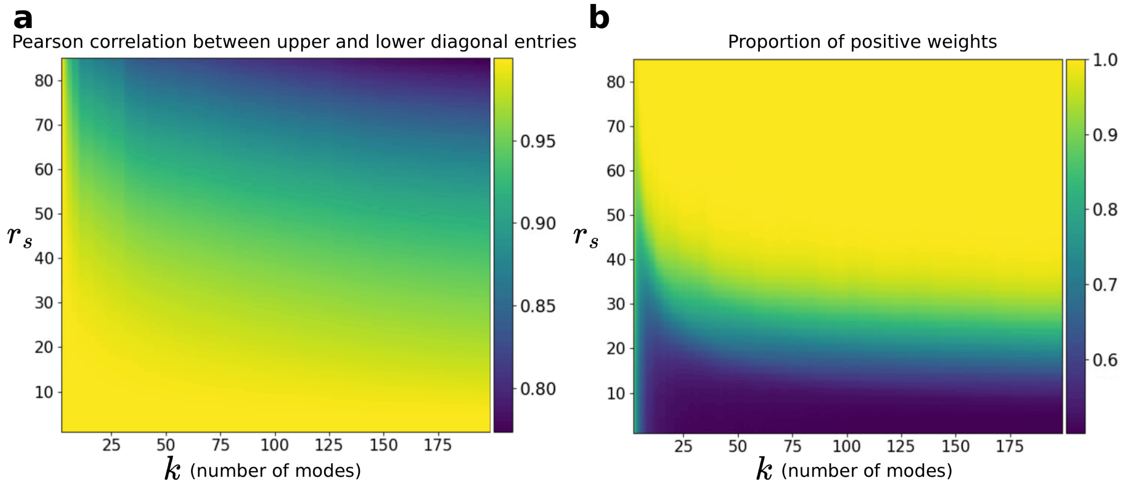


**Figure S10**: **Asymmetries and negative weights in the low-rank approximation of the Green's function. a:** We quantified the similarity between the entries $G_{ij}$ and $G_{ji}$ produced by Equation S20, without applying any threshold to the weighted matrix. Across a range of parameters between $1 \leq r_s \leq 85$ and $2 \leq k \leq 200$, we computed the Pearson correlation coefficient between the upper ($G_{ij}$) and lower triangular entries ($G_{ji}$). **b:** Proportion of positive weights (versus negative weights) produced by Equation S20 for the same range of parameters in **a**.

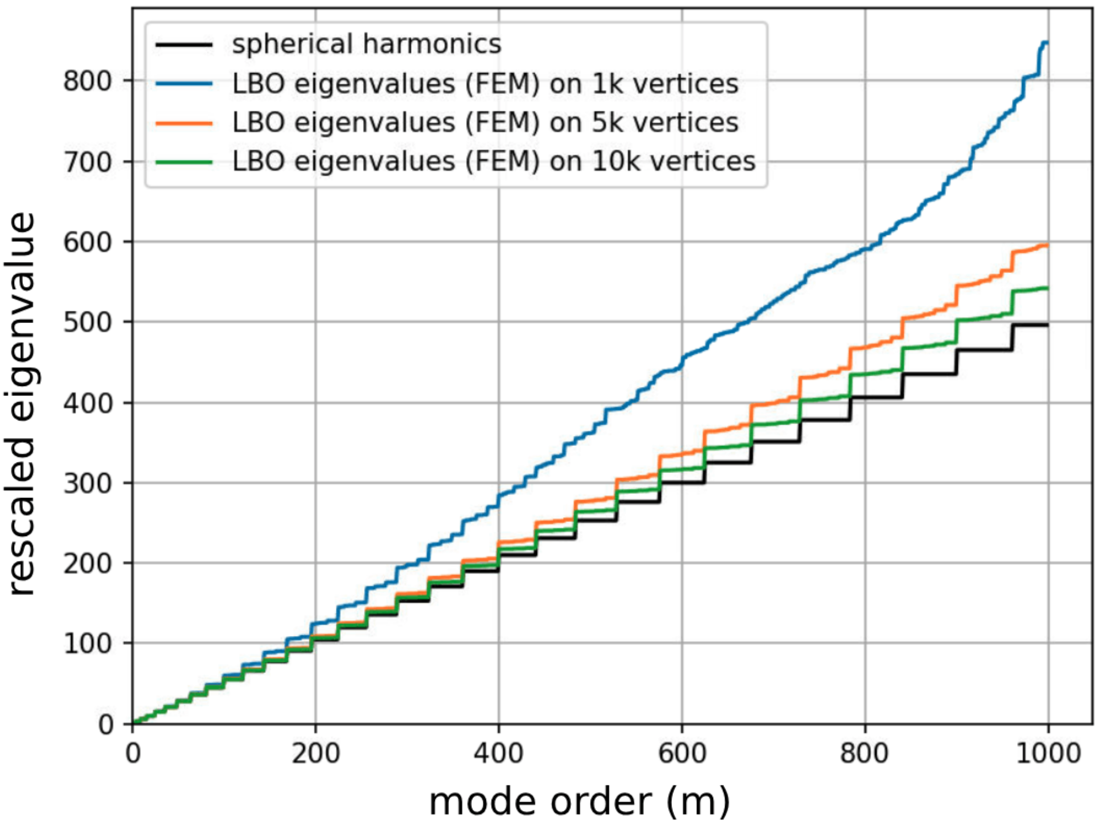


**Figure S11**: **Comparison between numerical and analytic estimates of spherical harmonics at different mesh resolutions.** Comparison of the eigenvalues of a spherical surface at different surface mesh resolutions (1000, 5000, and 10000 vertices) obtained through the finite-element method with the eigenvalues of the analytical spherical harmonics. Specifically, the spherical harmonics eigenvalues are given by $\ell(\ell + 1)$, with multiplicity $2\ell + 1$ due to the symmetries of the sphere, with $\ell = 0, 1, 2, 3, \ldots$, where $\ell$ corresponds to the degree of the polynomial of Laplace's spherical harmonics, or the angular momentum quantum number, and relates to the number of nodal lines (i.e., spatial frequencies) in the associated eigenvectors. Each eigenvalue set has been rescaled by dividing each value by the smallest non zero value.

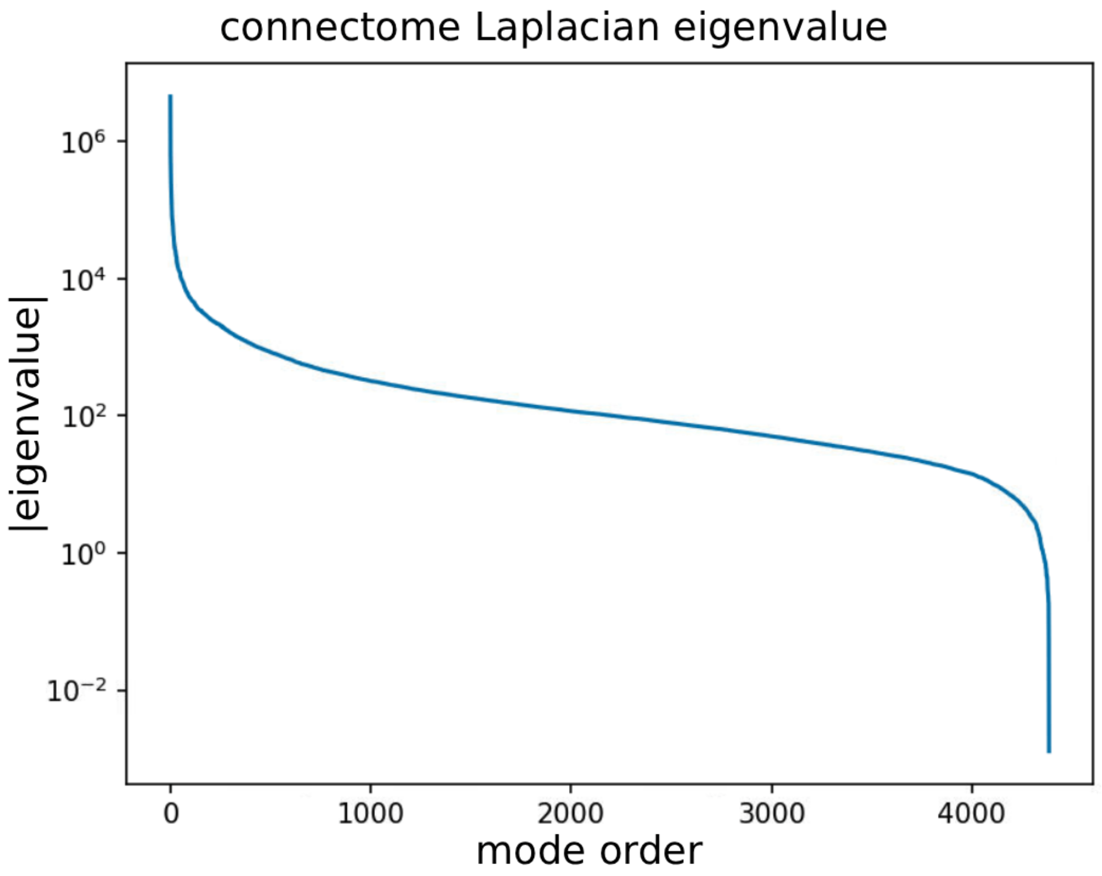


**Figure S12**: **Empirical human connectomes are effectively low-rank.** Absolute value of the graph Laplacian eigenvalues of the empirical smoothed high-resolution connectome of 4,386 vertices. The eigenvalues span 8 orders of magnitude, with the bulk of spectral energy concentrated in the earliest graph Laplacian eigenvectors, suggesting that empirical connectomes are relatively low-rank (see also Ref. (*56*)).

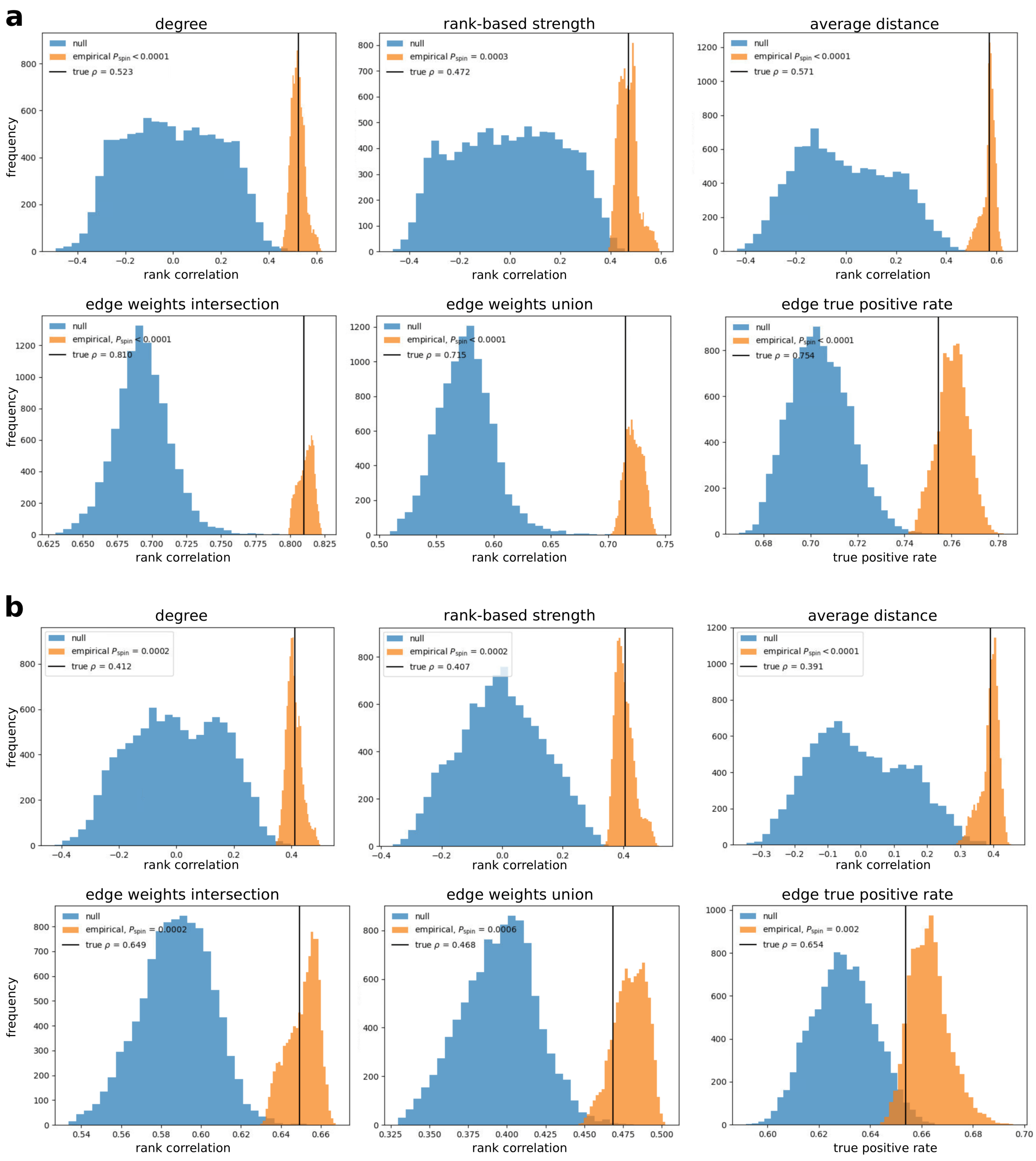


**Figure S13**: **Spin tests of the metrics derived from the high-resolution human connectome.** We performed pair-wise comparisons between the empirical and rotated spatial maps obtained from spin tests to calculate the $P$ values, $P_{\text{spin}}$, in Figs. 2 and S5 for the smoothed and non-smoothed connectomes, respectively (see section 0.1.10). **a:** Empirical and null distributions across network metrics for the smoothed high-resolution connectome. **b:** Same as panel **a**, but using the non-smoothed connectome.

##### 0.2.2 Atlas-based human connectome

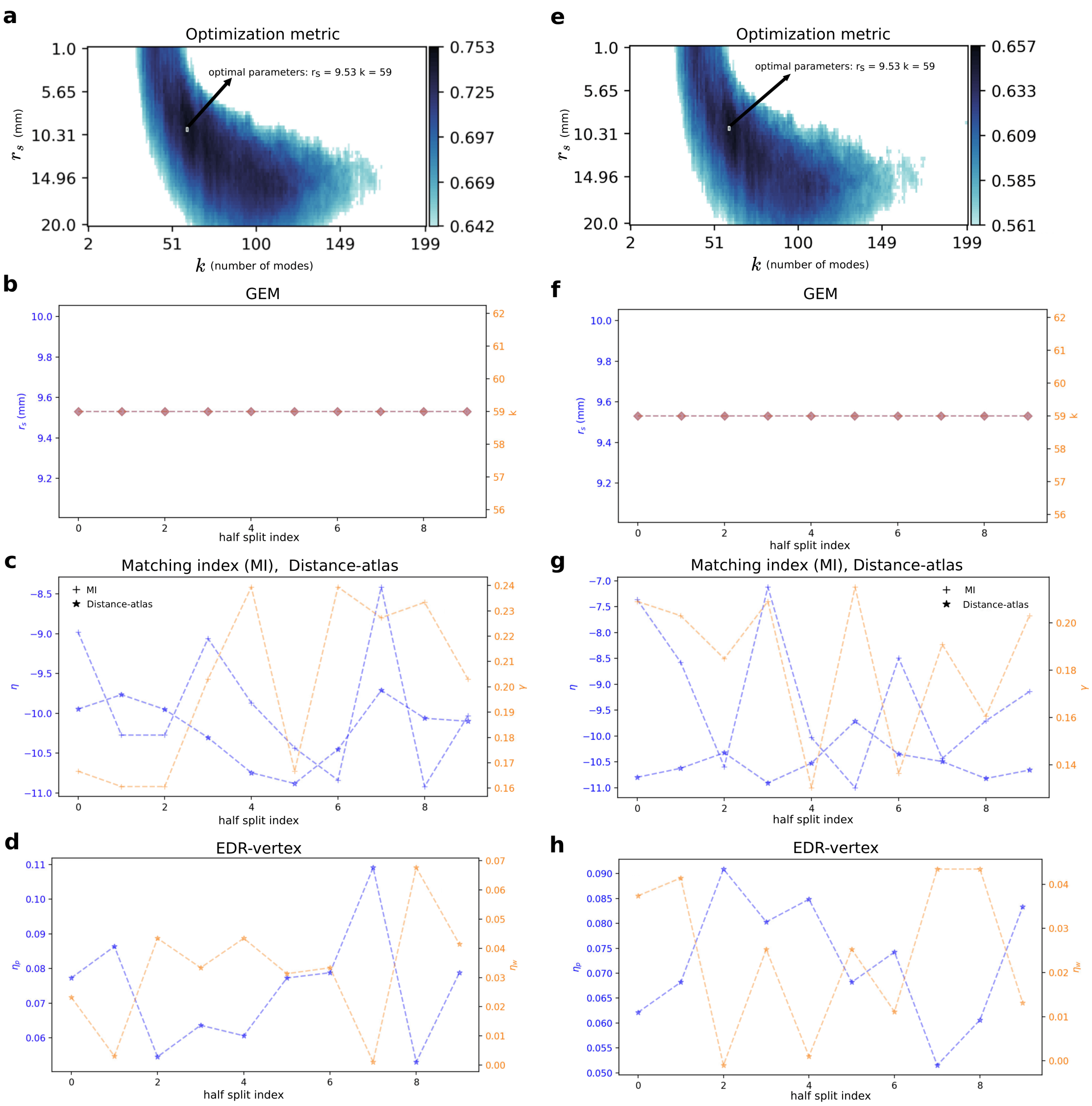

**Figure S14**: **Consistency of model parameters in the atlas-based human connectome.** High-resolution models (i.e., GEM and EDR-vertex) were optimized using the objective function in Equation S31 and atlas-based models (i.e., MI and Distance-atlas) were optimized using the objective function in Equation S33. **a:** GEM optimization landscape using the full sample to generate the smoothed high-resolution human connectome. The GEM has two free parameters: the length scale parameter $r_s$ (mm) and the number of geometric modes $k$. The optimal parameter combination is $r_s = 9.53$ mm and $k = 59$ modes. The score represents the simple mean of the three Spearman rank correlations between model and empirical network metrics: (1) the binary nodal degree; (2) the weighted nodal degree (rank-based strength); and (3) the model and empirical edge weights intersection. For clearer visualization, the landscape only shows the scores among the top 60%, coloured from light blue to black. **b:** Optimal parameters for the GEM for each half split cross-validation for the smoothed connectome. **c:** Optimal decay constant $\eta$ for the MI and Distance-atlas models and optimal parameter $\gamma$ for the MI for the smoothed connectome. **d:** Optimal decay constants of the EDR-vertex model for the probability of connection ($\eta_p$) and connection weight ($\eta_w$) for the smoothed connectome. **e:**, **f:**, **g**, and **h:** correspond to panels **a**, **b**, **c**, and **d**, respectively, but using the non-smoothed connectome. We found consistent optimal parameters across half split indices for the GEM for the smoothed and non-smoothed connectomes.

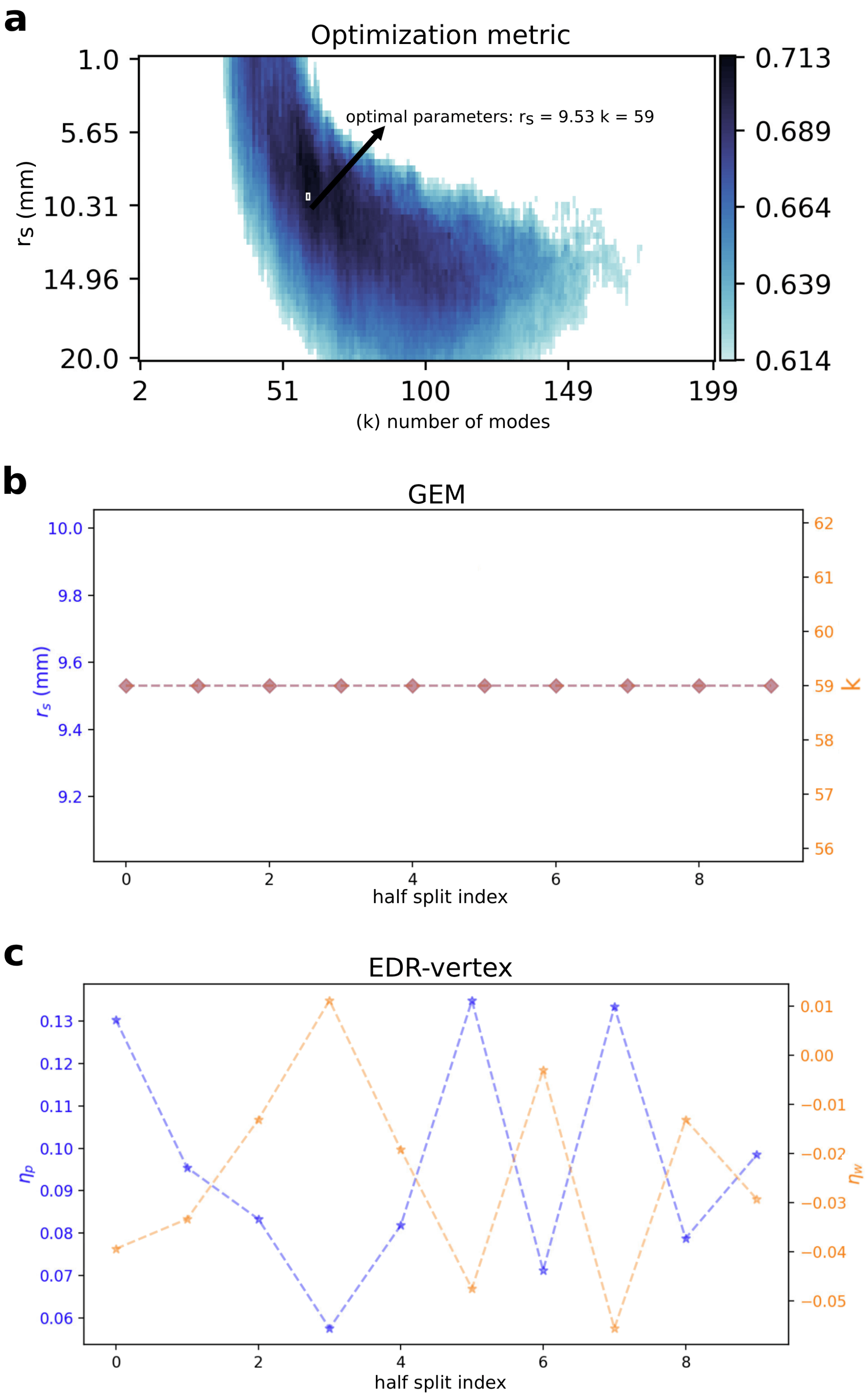
a
Optimization metric
optimal parameters: rs = 9.53 k = 59
rs (mm)
1.0
5.65
10.31
14.96
20.0
2
51
100
149
199
(k) number of modes
0.713
0.689
0.664
0.639
0.614
b
GEM
10.0
9.8
9.6
9.4
9.2
rs (mm)
62
61
60
59
58
57
56
K
0
2
4
6
8
half split index
c
EDR-vertex
0.13
0.12
0.11
0.10
0.09
0.08
0.07
0.06
ηp
0.01
0.00
−0.01
−0.02
−0.03
−0.04
−0.05
ηw
0
2
4
6
8
half split index

**Figure S15**: **GEM optimal parameters are consistent for different objective functions in the atlas-based human connectome.** We optimized the GEM and EDR-vertex models with the binary objective function given by Equation S33. We found that the optimal parameters of the GEM remained the same when optimized with the binary objective function; see Fig. S16 for model performances using this objective function. **a:** GEM optimization landscape with Equation S33 using the full sample to generate the smoothed high-resolution human connectome. The optimal parameter combination is $r_s = 9.53$ mm and $k = 59$ modes, which is identical to the one obtained with the weighted objective function in Fig. S14. The score represents the simple mean of Spearman rank correlation between model and empirical binary nodal degree and edge-wise true positive rate, corresponding to the binary objective function in Equation S33. For clearer visualization, the landscape only shows the scores among the top 60%, coloured from light blue to black. **b:** Optimal parameters for the GEM for each half split cross-validation for the smoothed connectome. **b:** Optimal decay constants of the EDR-vertex model for the probability of connection ($\eta_p$) and connection weight ($\eta_w$) for the smoothed connectome.

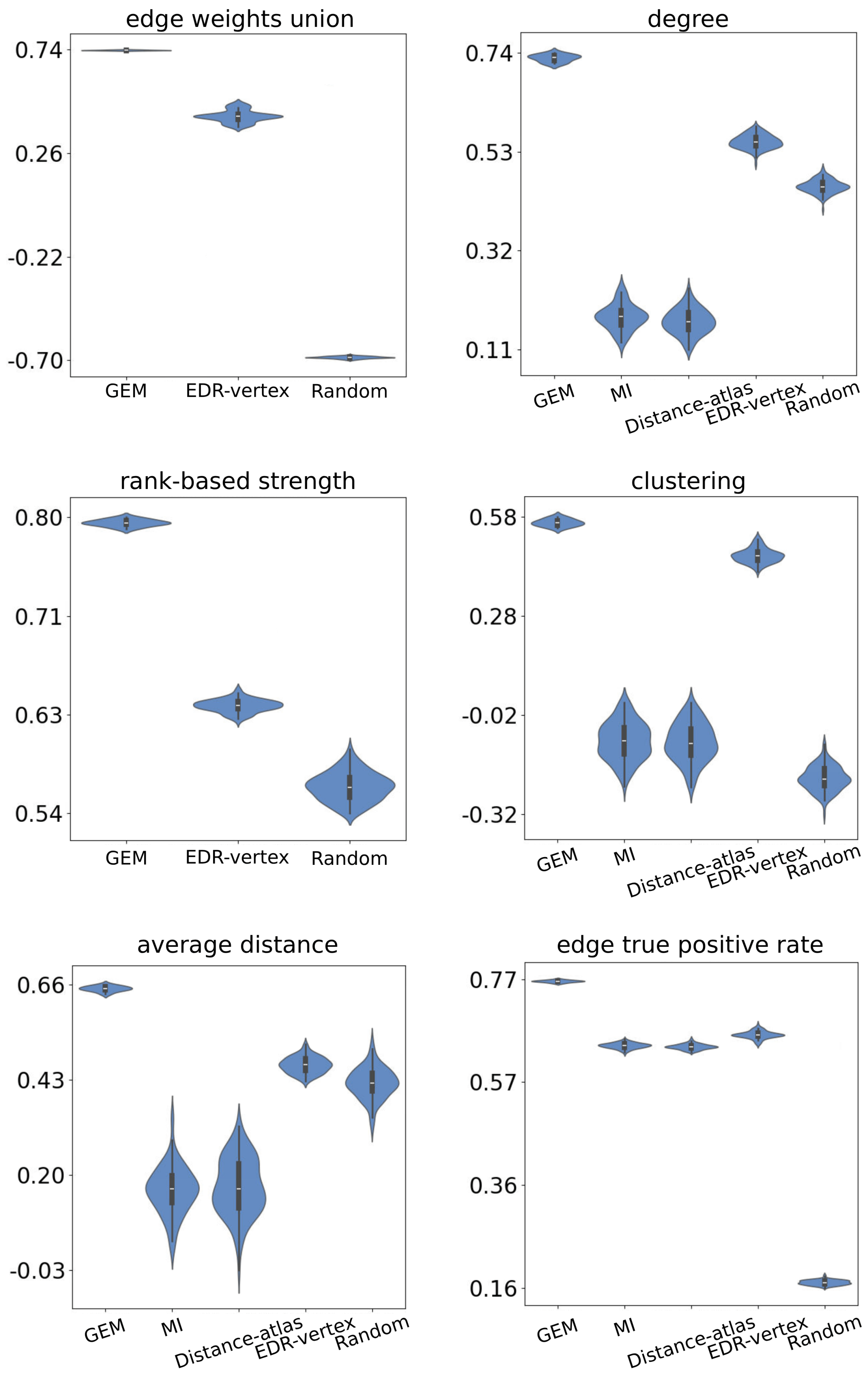
edge weights union
0.74
0.26
-0.22
-0.70
GEM
EDR-vertex
Random
degree
0.74
0.53
0.32
0.11
GEM
MI
Distance-atlas
EDR-vertex
Random
rank-based strength
0.80
0.71
0.63
0.54
GEM
EDR-vertex
Random
clustering
0.58
0.28
-0.02
-0.32
GEM
MI
Distance-atlas
EDR-vertex
Random
average distance
0.66
0.43
0.20
-0.03
GEM
MI
Distance-atlas
EDR-vertex
Random
edge true positive rate
0.77
0.57
0.36
0.16
GEM
MI
Distance-atlas
EDR-vertex
Random

**Figure S16**: **The superior performance of the GEM is not affected by the choice of the objective function in the atlas-based human connectome.** In Fig. 3, the GEM and EDR-vertex were optimized with the weighted objective function given by Equation S31. To allow fair comparison between these models and the binary-only atlas-based models (i.e., the MI and Distance-atlas), we also optimized the GEM and EDR-vertex models with the binary objective function given by Equation S33, so that all models were optimized with the same objective function; see Fig. S15 for optimal parameters. We found that the superior performance of the GEM was robust to the choice of the objective function across all network metrics; see section 0.1.8 and 0.1.9 for details about each measure. As in Fig. 3d, cross-validated fits were used to evaluate the model performances on the smoothed empirical connectome.

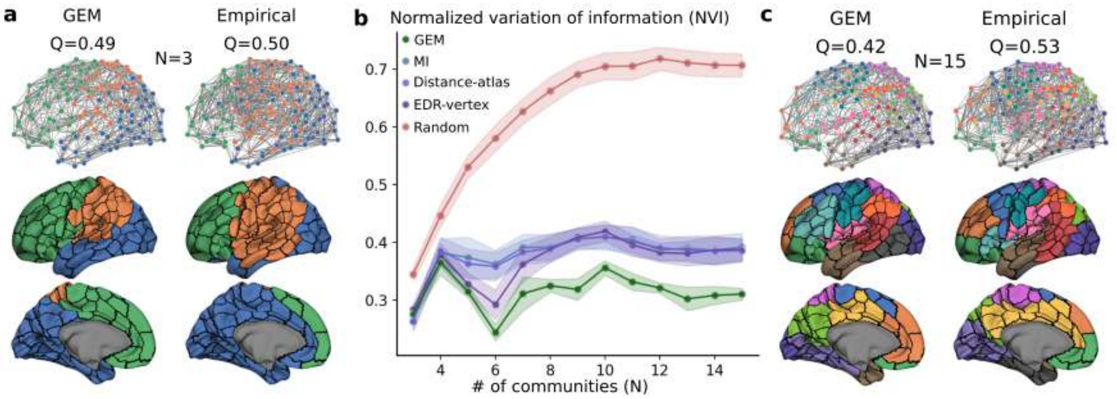


**Figure S17**: **The GEM captures the multiscale topography of empirical atlas-based human connectome's community structure better than benchmark models.** Using cross-validated fits, we quantified the agreement between the modular architecture of model and empirical smoothed atlas-based human connectomes at different community resolutions using the normalized variation of information (NVI; see section 0.1.9 for details), for which lower values indicate better agreement. We calculated the NVI for a range of numbers of communities between $N = 3$ and $N = 15$. **a:** Visualization on the $N = 150$-region cortical atlas of the GEM and empirical communities for a solution comprising 3 communities, with the modularity $Q$. Each colour represents a set of vertices within a community, with each vertex assigned to a single community. **b:** Using cross-validated fits, the NVI across different numbers of communities for every model was: GEM (AUC = 3.79), MI (AUC = 4.62), Distance-atlas (AUC = 4.59), EDR-vertex (AUC = 4.46), and Random (AUC = 7.61), where AUC is the area under the curve, with lower values indicating greater similarity to the empirical community structure. The shaded area around the curves represents the standard deviation. **c:** Model and empirical communities for a solution comprising 15 communities.

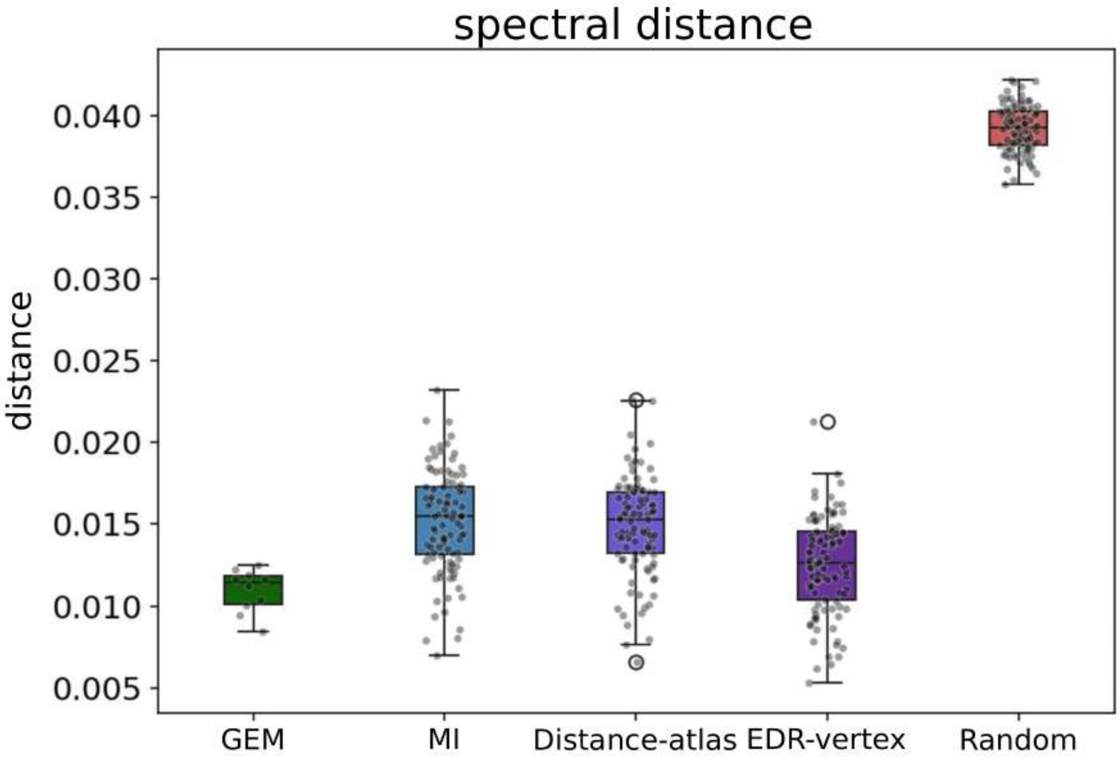


**Figure S18**: **Spectral distances between models and data for the smoothed atlas-based human connectome using cross-validated fits.** We assessed the topological similarity between the GEM and empirical connectome using their spectral distance, with lower values indicating better agreement; see section 0.1.9.

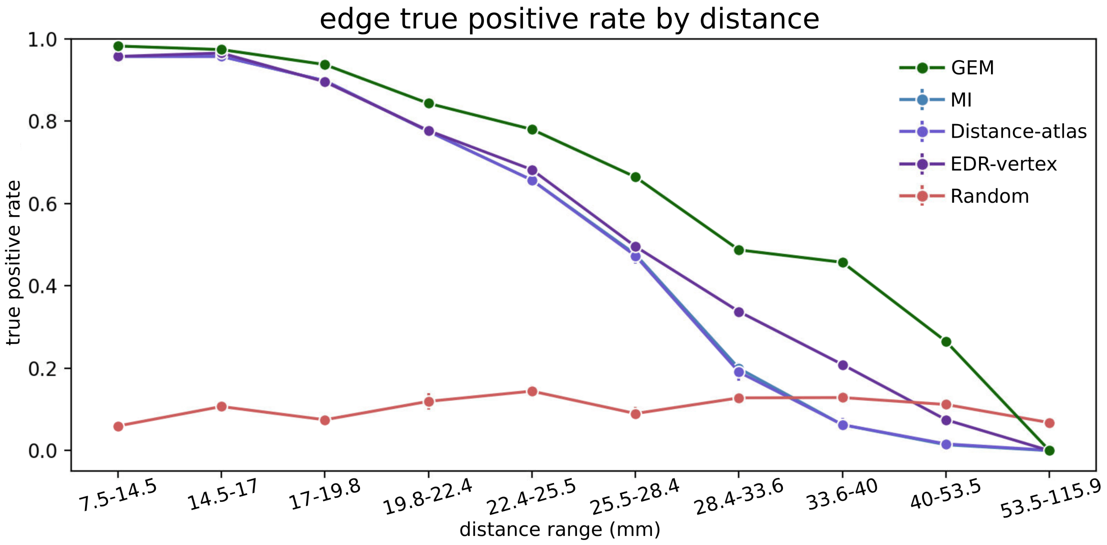
edge true positive rate by distance
true positive rate
distance range (mm)
GEM
MI
Distance-atlas
EDR-vertex
Random
1.0
0.8
0.6
0.4
0.2
0.0
7.5-14.5
14.5-17
17-19.8
19.8-22.4
22.4-25.5
25.5-28.4
28.4-33.6
33.6-40
40-53.5
53.5-115.9

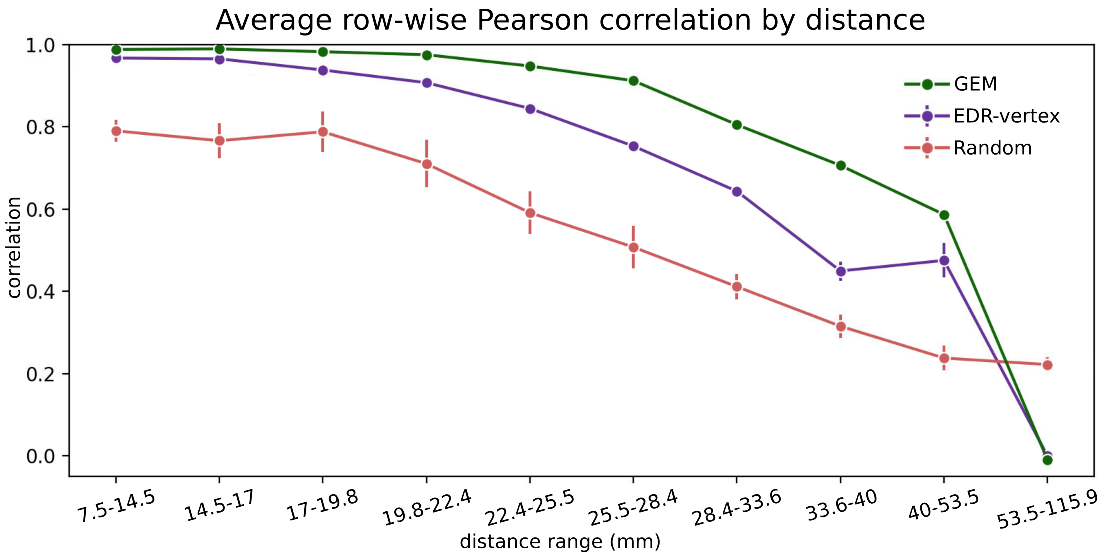
Average row-wise Pearson correlation by distance
correlation
distance range (mm)
GEM
EDR-vertex
Random
1.0
0.8
0.6
0.4
0.2
0.0
7.5-14.5
14.5-17
17-19.8
19.8-22.4
22.4-25.5
25.5-28.4
28.4-33.6
33.6-40
40-53.5
53.5-115.9

**Figure S19**: **The GEM better capture empirical edges than benchmark models across all connection lengths in the atlas-based human connectome.** We assessed the GEM's ability to predict empirical edges in the atlas-based smoothed connectome across 10 connection-length ranges, each containing an equal number of edges. **a:** Edge-wise true positive rate (binary) as a function of connection length for the GEM, MI, Distance-atlas, EDR-vertex, and Random models. **b:** Average row-wise Pearson correlation across connection lengths for the GEM, EDR-vertex, and Random models. To quantify correlations, we adapted the approach of (*15*): for each length range, we excluded edges outside the range, computed row-wise Pearson correlations between the model and empirical connectomes (preserving the weights), and averaged these correlations to obtain a single similarity score per range of connection lengths. Note that the MI and Distance-atlas model can only generate binary connections, so we could not evaluate their performance with respect to this measure.

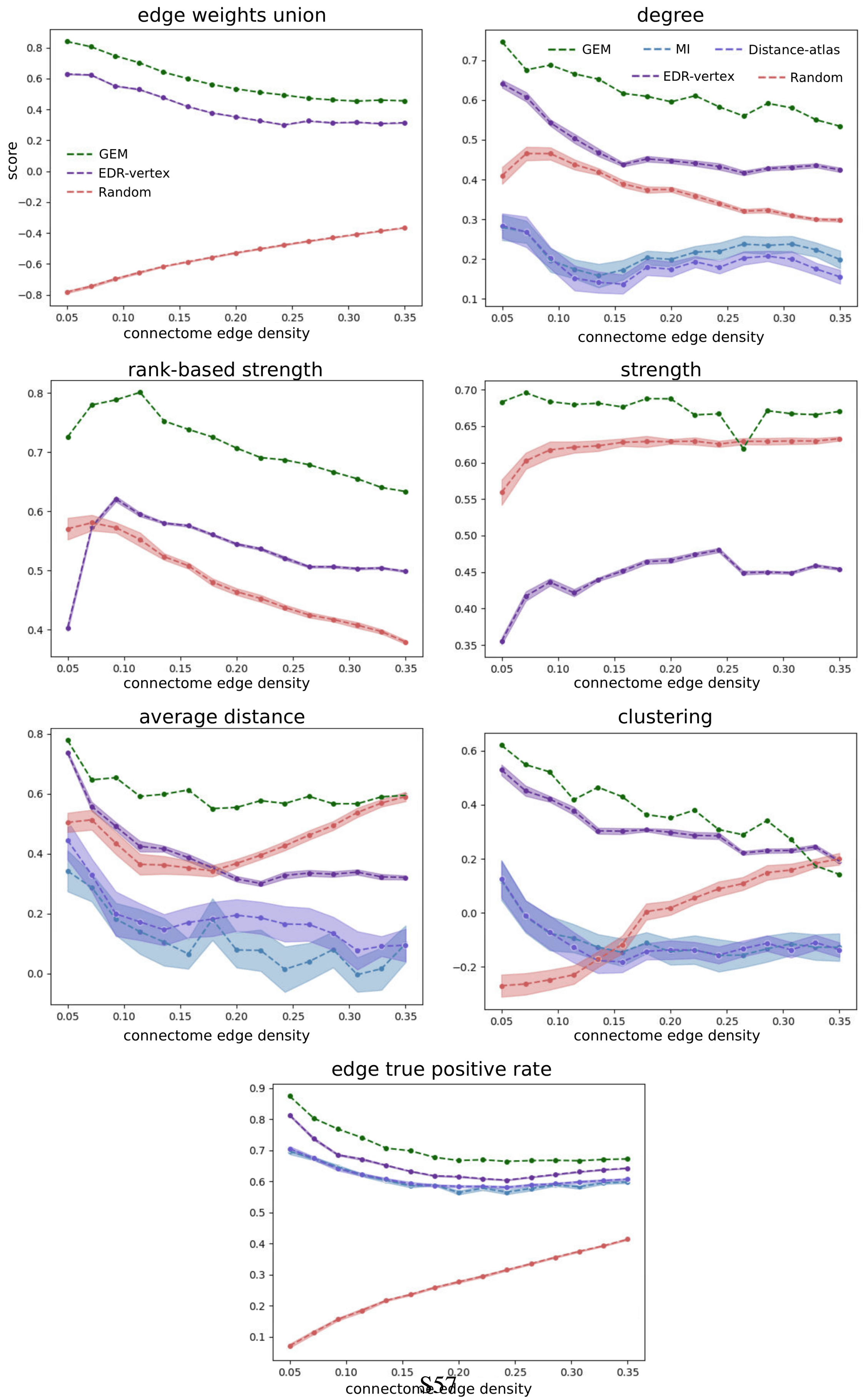
edge weights union
score
GEM
EDR-vertex
Random
connectome edge density
degree
GEM
MI
Distance-atlas
EDR-vertex
Random
rank-based strength
strength
average distance
clustering
edge true positive rate

**Figure S20**: **The GEM outperforms benchmark models across different edge densities in the atlas-based human connectome.** Using the full sample, we quantified the performance of each model across a range of empirical smoothed connectome edge densities equally spaced between 5 and 35%; see section 0.1.7 for details. We evaluated performance based on 7 edge- and node-level network metrics; see sections 0.1.8 and 0.1.9 for details about each measure. Note that the MI and Distance-atlas can only model binary connections, so we cannot evaluate their performance with respect to the strength, rank-based strength, and edge weights union correlation. High-resolution models (i.e., GEM and EDR-vertex) were optimized using the objective function in Equation S31 and atlas-based models (i.e., MI and Distance-atlas) were optimized using the objective function in Equation S33.

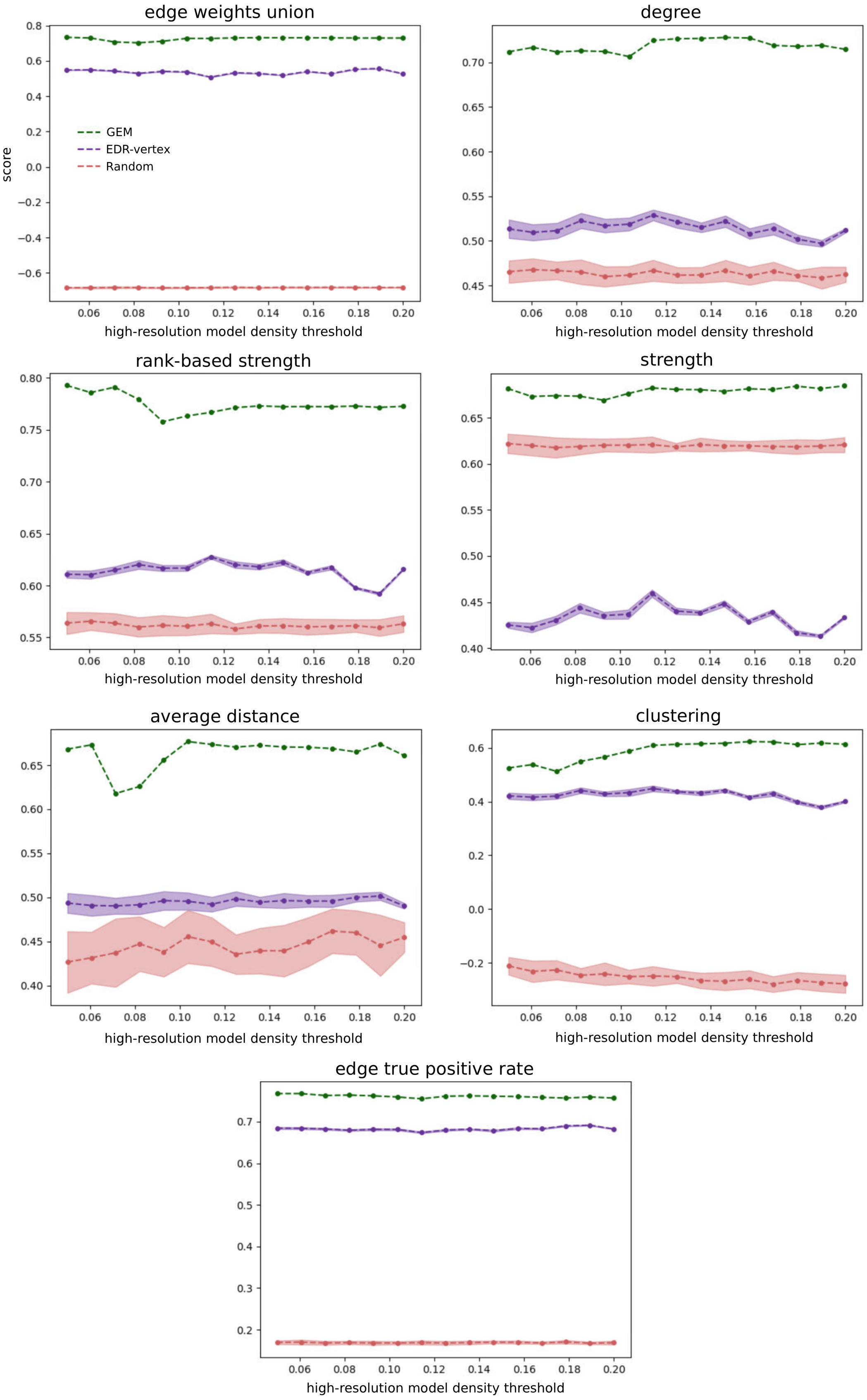
edge weights union
score
GEM
EDR-vertex
Random
high-resolution model density threshold
degree
high-resolution model density threshold
rank-based strength
high-resolution model density threshold
strength
high-resolution model density threshold
average distance
high-resolution model density threshold
clustering
high-resolution model density threshold
edge true positive rate
high-resolution model density threshold

**Figure S21**: **The GEM's superior performance in the atlas-based human connectome is robust to the high-resolution edge density threshold.** High-resolution models (i.e., GEM, EDR-vertex, and Random) were first generated at the vertex resolution, before being parcellated and compared with the smoothed atlas-based connectome. The edge density of the high-resolution models prior to applying the parcellation can affect the bias imparted to these models; see section 0.1.6. In Figs. 3 and S22, we used a density threshold of 4.6% for all high-resolution models, which corresponds to the average edge density of the high-resolution empirical connectomes; see section 0.1.1. Using the full sample, we evaluated the performance of the GEM and high-resolution benchmark models (i.e., EDR-vertex and Random) across different values of this high-resolution model edge threshold. We sampled 15 equally spaced values between 5 and 20%. We found that the GEM consistently shows superior performance across thresholds, and is therefore not sensitive to this additional parameter.

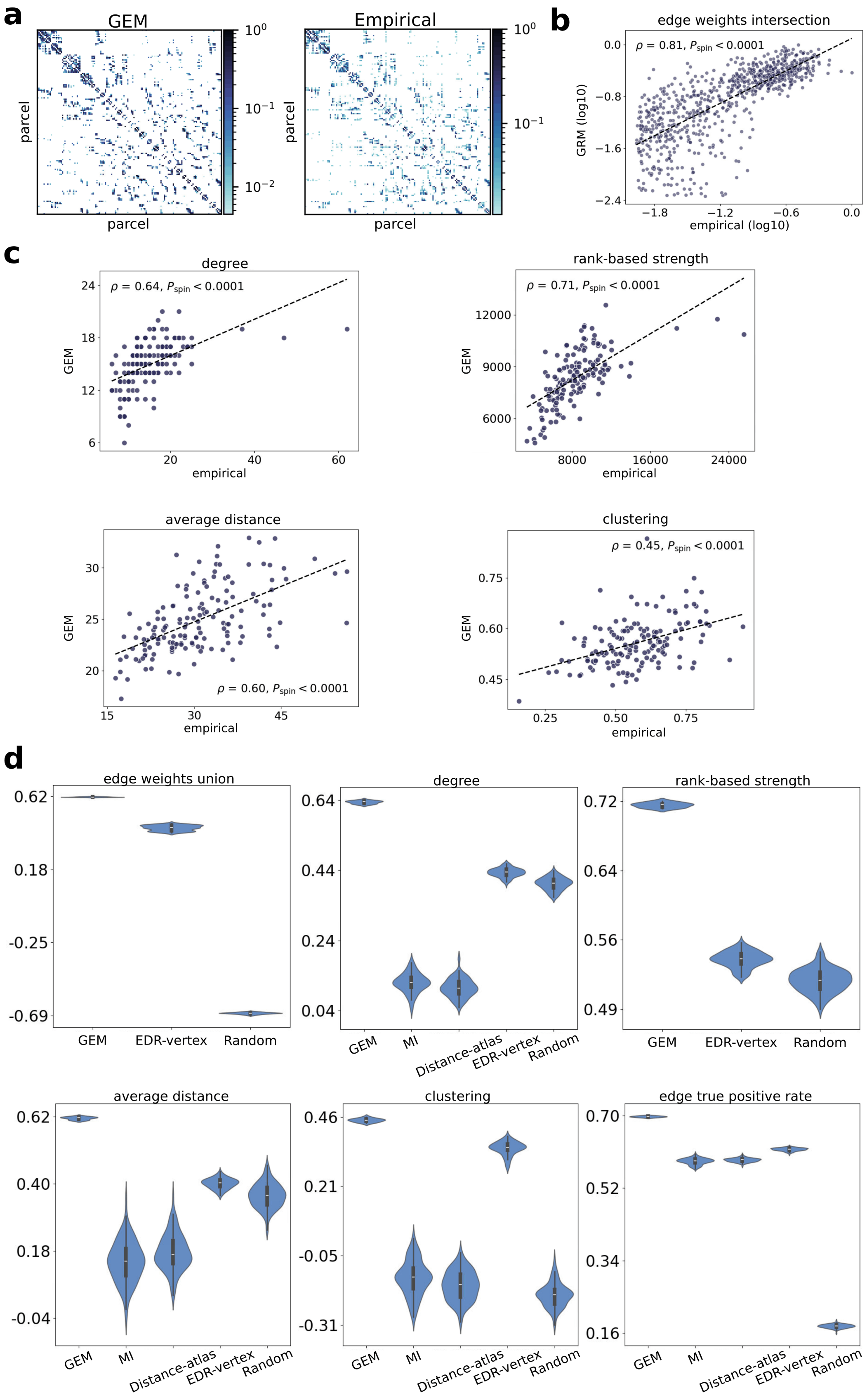
a
GEM
Empirical
parcel
parcel
parcel
parcel
b
edge weights intersection
$\rho$ = 0.81, $P_{spin}$ < 0.0001
GRM (log10)
empirical (log10)
c
degree
$\rho$ = 0.64, $P_{spin}$ < 0.0001
GEM
empirical
rank-based strength
$\rho$ = 0.71, $P_{spin}$ < 0.0001
GEM
empirical
average distance
$\rho$ = 0.60, $P_{spin}$ < 0.0001
GEM
empirical
clustering
$\rho$ = 0.45, $P_{spin}$ < 0.0001
GEM
empirical
d
edge weights union
GEM
EDR-vertex
Random
degree
GEM
MI
Distance-atlas
EDR-vertex
Random
rank-based strength
GEM
EDR-vertex
Random
average distance
GEM
MI
Distance-atlas
EDR-vertex
Random
clustering
GEM
MI
Distance-atlas
EDR-vertex
Random
edge true positive rate
GEM
MI
Distance-atlas
EDR-vertex
Random

**Figure S22**: **Modelling non-smoothed atlas-based human connectome.** We repeated the analysis shown in Fig. 3, but instead using the non-smoothed atlas-based connectome. We found that all models generally performed better on the smoothed connectome, but the strong and superior performance of the GEM compared to the benchmark models remains even without smoothing the empirical connectome. **a:** Optimized GEM and empirical atlas-based connectome matrices. The optimized model was generated using parameters $r_s = 9.53$ mm and $k = 59$ modes, which maximizes the objective function in Equation S31. **b:** Scatter plot of the common (intersection) edge weights between the GEM and empirical connectome using the full dataset. $P_{\text{spin}}$ is the one-sided p-value estimated from 10,000 spin-based permutations; see section 0.1.10 for details. **c:** Scatter plots of topographical agreement between the optimized model and empirical data using the full dataset, quantified using Spearman rank correlations ($\rho$). From left to right, top panels show results for the degree (binary degree) and rank-based strength (weighted degree), while bottom panels show results for the average node connection distance and nodal clustering coefficient. **d:** Comparison of GEM performance relative to other models, as evaluated using split-half cross-validation, for the N=150 regions atlas. Note that the MI and Distance-atlas can only model binary connections, so we cannot evaluate their performance with respect to the rank-based strength and edge weights union correlation. GEM=Geometric Eigenmode Model; MI=Matching Index model; Distance-atlas=Power law distance rule model defined at atlas resolution; EDR-vertex=Exponential Distance Rule model defined at the vertex level and then parcellated; Random=Random network defined at vertex resolution and then parcellated (see Methods for details).

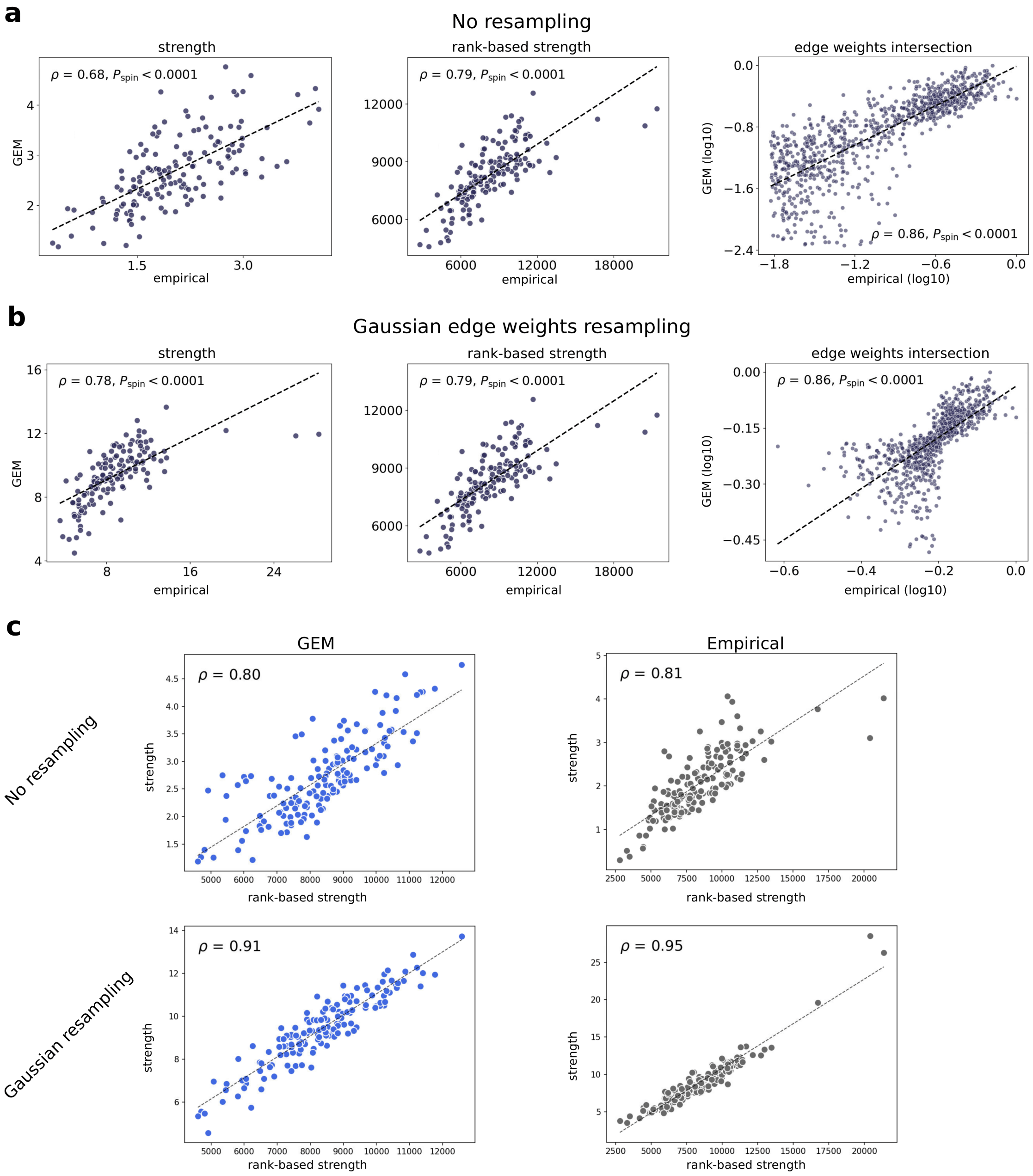
a
No resampling
strength
rank-based strength
edge weights intersection
$\rho$ = 0.68, $P_{spin} < 0.0001$
$\rho$ = 0.79, $P_{spin} < 0.0001$
$\rho$ = 0.86, $P_{spin} < 0.0001$
GEM
GEM (log10)
empirical
empirical (log10)
b
Gaussian edge weights resampling
strength
rank-based strength
edge weights intersection
$\rho$ = 0.78, $P_{spin} < 0.0001$
$\rho$ = 0.79, $P_{spin} < 0.0001$
$\rho$ = 0.86, $P_{spin} < 0.0001$
GEM
GEM (log10)
empirical
empirical (log10)
c
GEM
Empirical
No resampling
Gaussian resampling
$\rho$ = 0.80
$\rho$ = 0.81
$\rho$ = 0.91
$\rho$ = 0.95
strength
rank-based strength

**Figure S23**: **Agreement between different definitions of nodal strengths and effect of edge weights resampling in the atlas-based human connectome.** Same as Fig. S6, but in the atlas-based connectome. We compared different definitions of nodal strength before and after resampling the edge weights using the approach in (*68*). Since the resampling of edge weights preserves their ranking, it has no effect on rank correlations at the edge level. We found that the two definitions of nodal strength given in Equations S29 and S30 show good agreement in both the GEM and empirical connectome. We also observed that resampling the edge weights increases the agreement between the two definitions. **a:** Panels from left to right show scatter plots of nodal strength, rank-based strength and the edge weights intersection between GEM and atlas-based empirical smoothed human connectome. **b:** Same as panel a, but after resampling the edge weights. **c:** Scatter plot of the strength versus rank-based strength in the GEM (top left). Scatter plot of the strength versus rank-based strength in the empirical connectome (top right). Scatter plot between strength and rank-based strength after resampling the edge weights in the GEM (bottom left). Scatter plot between strength and rank-based strength after resampling the edge weights in the empirical connectome (bottom right).

##### 0.2.3 Non-human mammalian species

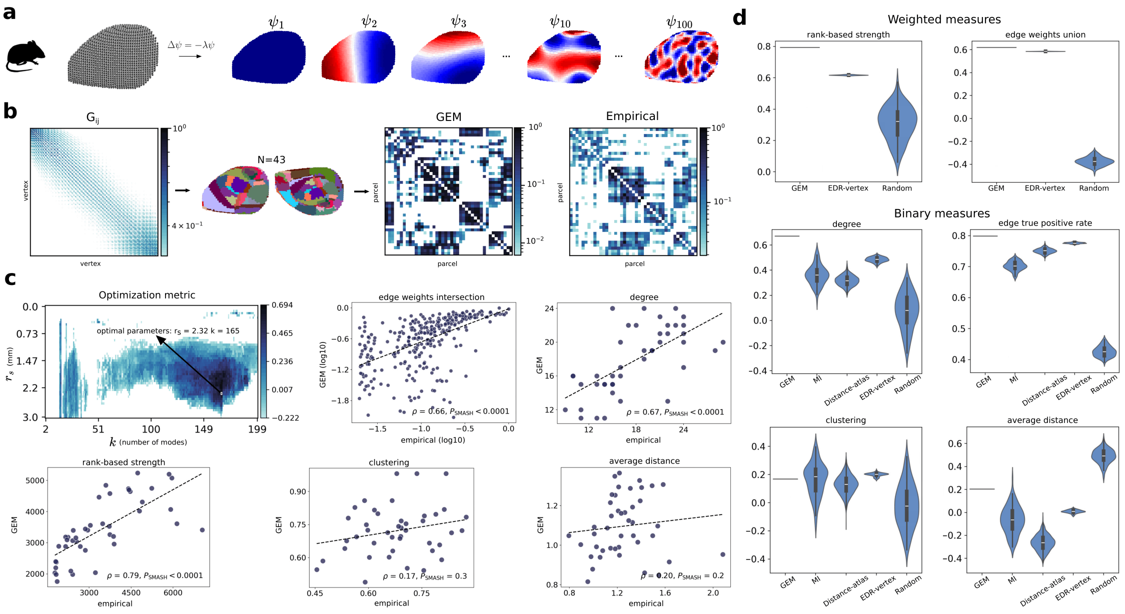


**Figure S24**: **The GEM better captures the topology and topography of the sparse empirical mouse connectome compared to benchmark models.** We compared the performance of the GEM with different benchmark models in capturing diverse topological and topographical properties of the empirical mouse connectome; see section 0.1.8 and section 0.1.9 for details about the network measures. To compare with all benchmark models available, we used the sparse connectome. The GEM and EDR-vertex model were optimized using Equation S31, while the MI and Distance-atlas models were optimized using Equation S33. **a:** We obtained the geometric eigenmodes from a tetrahedral volumetric mesh of the right hemispheric isocortex of the mouse, which are ordered from long- to short-wavelengths; see section 0.1.3 for details about obtaining the modes. **b:** The GEM was generated at the resolution of the vertices of the mesh, and then parcellated onto the $N = 43$ regions atlas, which can be compared to the atlas-based empirical mouse connectome; see section 0.1.7 for details about the generation process and section 0.1.2 for details about the connectome. **c:** Top row: GEM optimization landscape, with optimal parameters $r_S = 2.32$mm and $k = 165$ modes (left) and scatter plots of topographical agreement between model and empirical nodal binary (degree) and weighted degree (rank-based strength) (center and right). Bottom row: Scatter plots of topographical agreement of the rank-based strength, clustering and average connection length, respectively. **d:** Performance comparison between the GEM and benchmark models, i.e. the MI, Distance-atlas, EDR-vertex, and Random; see section 0.1.5 for details about each model. All benchmarks were generated 100 times using their optimal parameter(s), and the fit statistics evaluated each time. The violin plots show the distribution of these 100 fits.

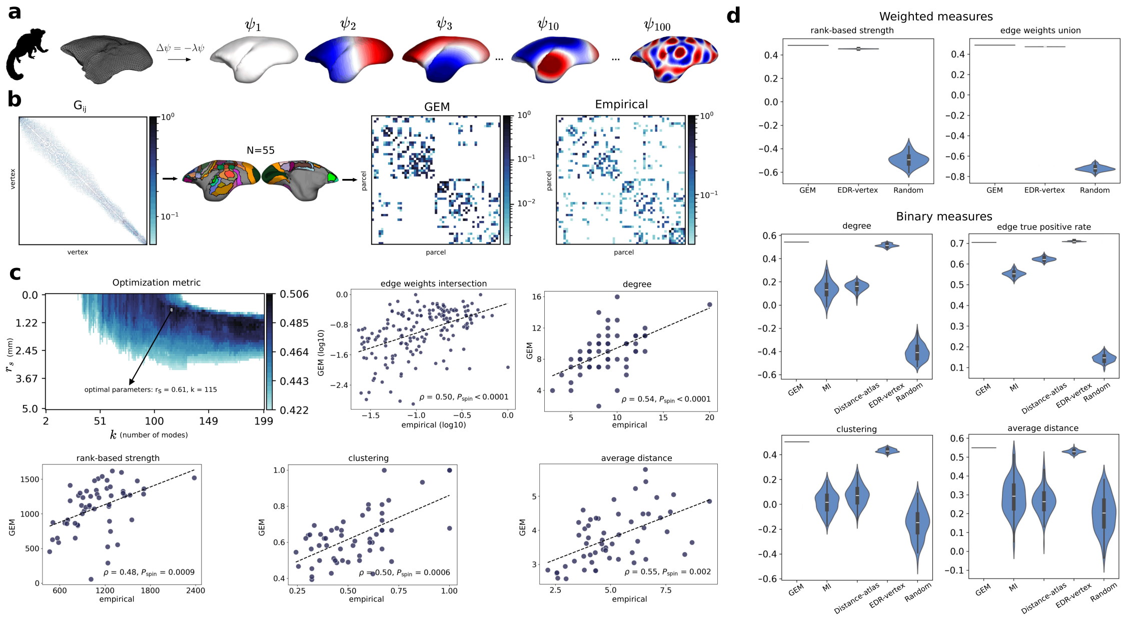

**Figure S25**: **The GEM better captures the topology and topography of the empirical marmoset connectome compared to benchmark models.** Same as Fig. S24, but for the marmoset connectome; see section 0.1.7 for details about generating the models and see section 0.1.2 for details about the connectome.

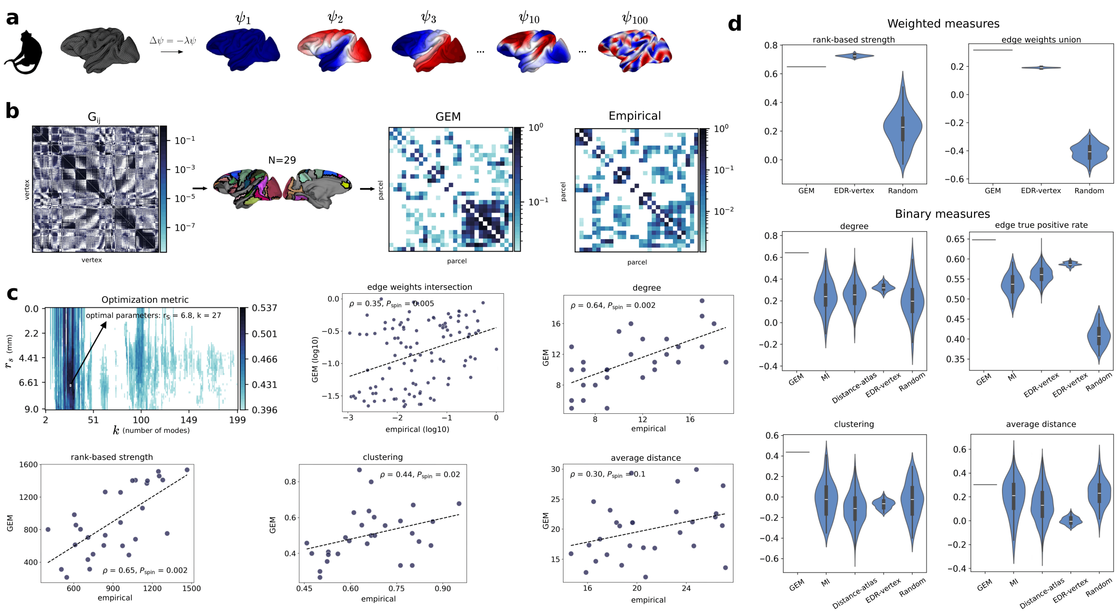


**Figure S26**: **The GEM better captures the topology and topography of the empirical macaque sparse connectome compared to benchmark models.** Same as Fig. S24, but for the sparse macaque connectome; see section 0.1.7 for details about generating the models and see section 0.1.2 for details about the connectome.

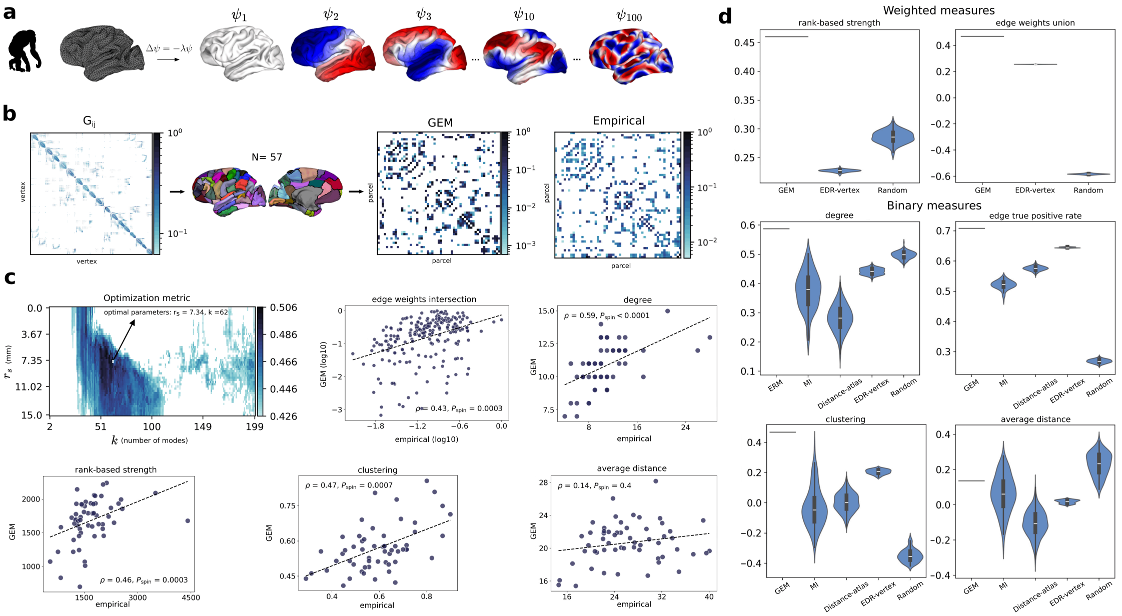


**Figure S27**: **The GEM better captures the topology and topography of the empirical chimpanzee connectome compared to benchmark models.** Same as Fig. S24, but for the chimpanzee connectome; see section 0.1.7 for details about generating the models and see section 0.1.2 for details about the connectome.

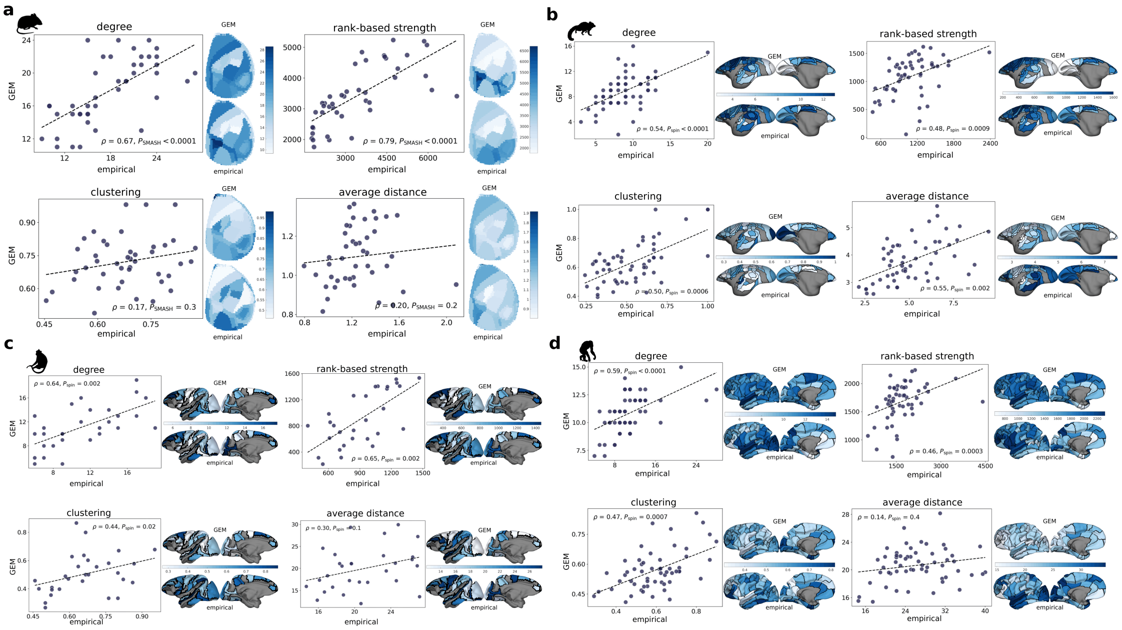

**Figure S28**: **Spatial maps of network metrics in non-human mammalian species.** All GEMs were optimized with the objective function given by Equation S31. In each species, we compared the GEM and empirical spatial maps of four network measures. From left to right, the top rows show the binary nodal degree and the rank-based nodal strength, and the bottom rows show the nodal clustering and average nodal connection length. The panels show the spatial maps for the **a:** sparse mouse connectome (see Fig. S24), **b:** marmoset connectome (see Fig. S25), **c:** sparse macaque connectome (see Fig. S26), and **d:** chimpanzee connectome (see Fig. S27).

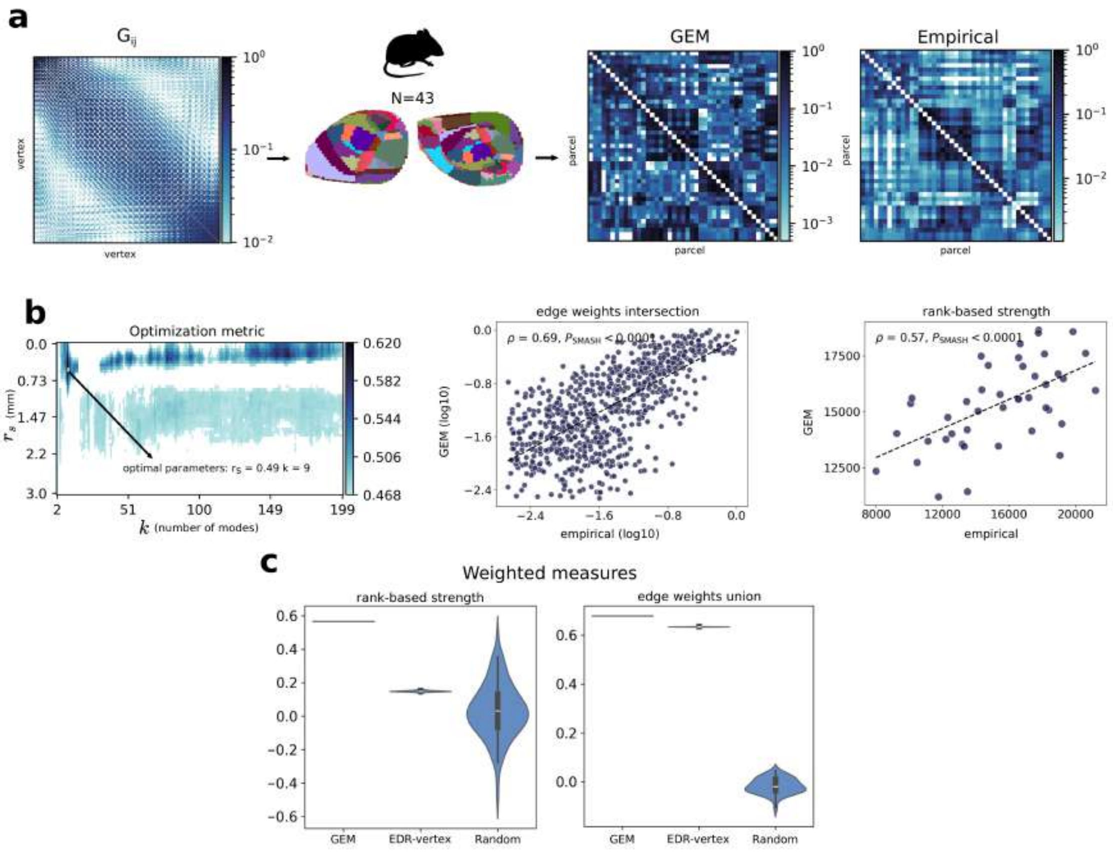


**Figure S29**: **The GEM better captures the topology and topography of the dense empirical mouse connectome compared to benchmark models.** In Fig. S24, we compared models using a sparse version of the mouse connectome. Here we used the dense connectome and compare with the high-resolution benchmark models, i.e., EDR-vertex and Random. In this case, we focused on weighted network properties and therefore optimized the models using Equation S32. **a:** The GEM was generated at the resolution of the vertices of the mesh, and then parcellated onto the $N = 43$ regions atlas, and then compared with the empirical mouse dense connectome. **b:** GEM optimization landscape, with optimal parameters $r_s = 0.49$ mm and $k = 9$ modes (left) and scatter plots of the edge weights intersection (edges common to both model and empirical) and topographical agreement between model and empirical weighted degree (rank-based strength) (center and right). **c:** Comparison of the GEM's performance relative to benchmark models.

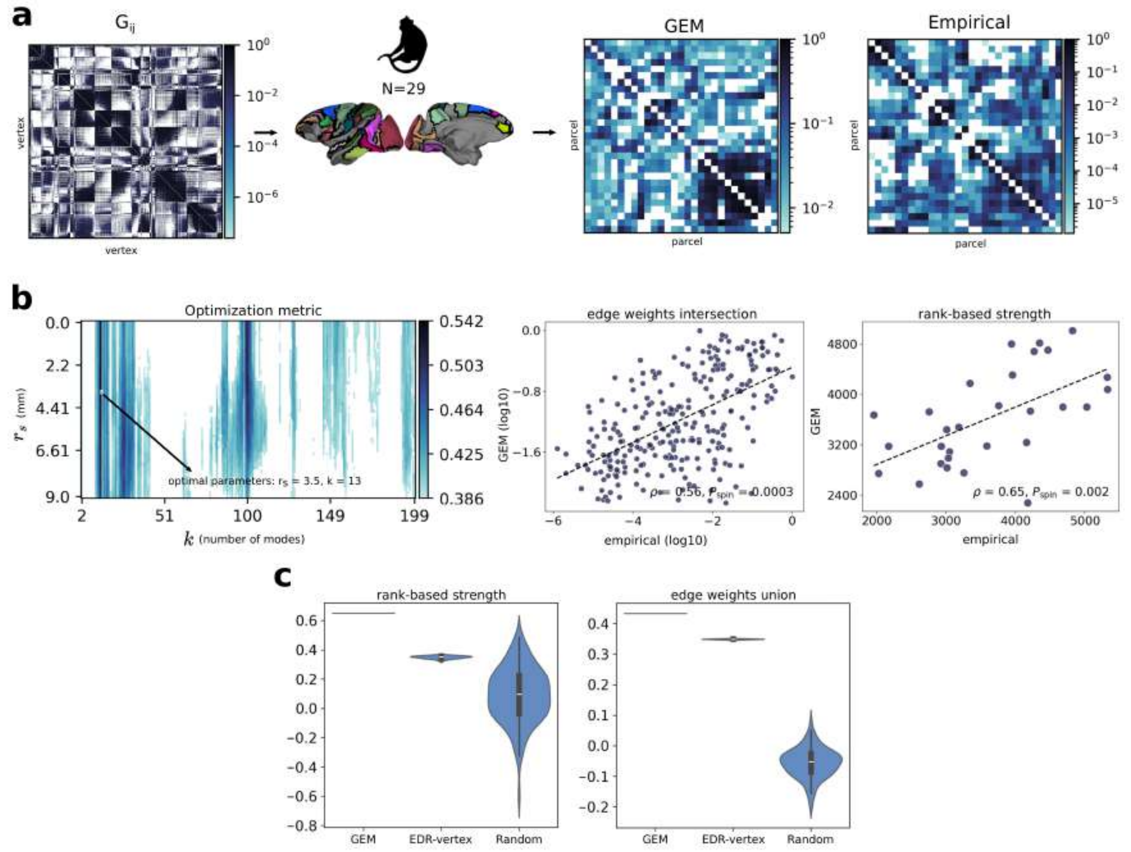


**Figure S30**: **The GEM better captures the topology and topography of the dense empirical macaque connectome compared to benchmark models.** Same as Fig. S29, but for the macaque connectome; see section 0.1.7 for details about the models and section 0.1.2 for details about the connectome.

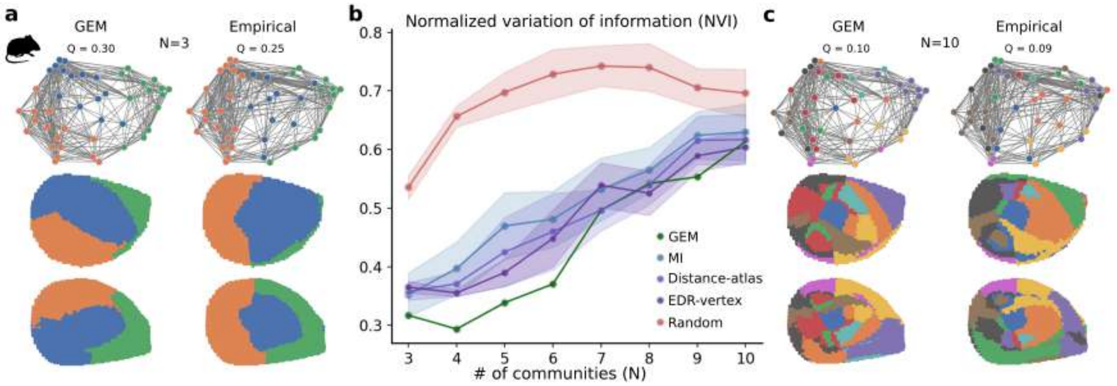


**Figure S31**: **The GEM captures the multiscale topography of the empirical atlas-based mouse connectome's community structure better than benchmark models.** We quantified the agreement between the modular architecture of model and the sparse empirical mouse connectome at different community resolutions using the normalized variation of information (NVI; see section 0.1.9), for which lower values indicate better agreement. We calculated the NVI for a range of numbers of communities between $N = 3$ and $N = 10$. **a:** Visualization on the $N = 43$ regions cortical atlas of the GEM and empirical communities for a solution comprising 3 communities, with the modularity $Q$. Each colour represents a set of vertices within a community, with each vertex assigned to a single community. **b:** NVI across different numbers of communities for every model, i.e., GEM (AUC = 3.06), MI (AUC = 3.56), Distance-atlas (AUC = 3.39), EDR-vertex (AUC = 3.33). and Random (AUC = 4.89), where AUC is the area under the curve, with lower values indicating greater similarity to the empirical community structure. The shaded area around the curve of the probabilistic models (i.e., all models except the GEM) represents the standard deviation. **c:** Model and empirical communities for a solution comprising 10 communities, with modularity $Q$.

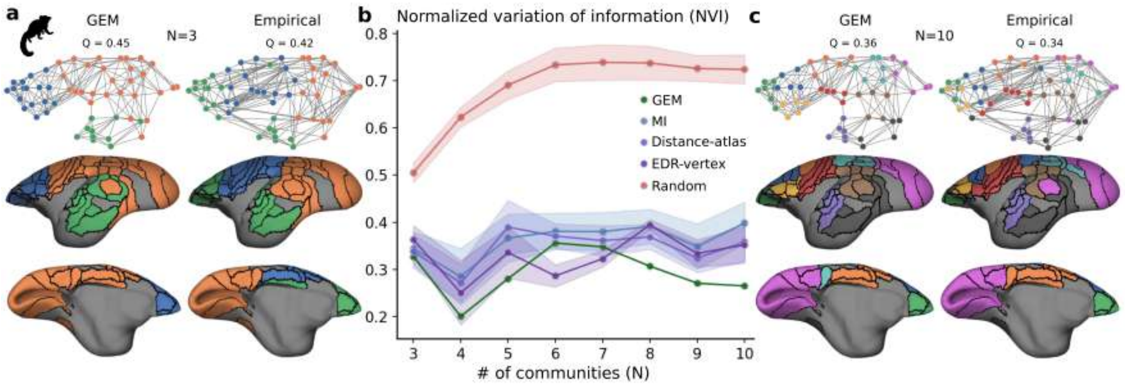


**Figure S32**: **The GEM captures the multiscale topography of the empirical atlas-based marmoset connectome's community structure better than benchmark models.** Same as Fig. S31, but for the marmoset connectome. The AUC for each model is: GEM = 2.06, MI = 2.52, Distance-atlas = 2.44, EDR-vertex = 2.28. and Random = 4.86.

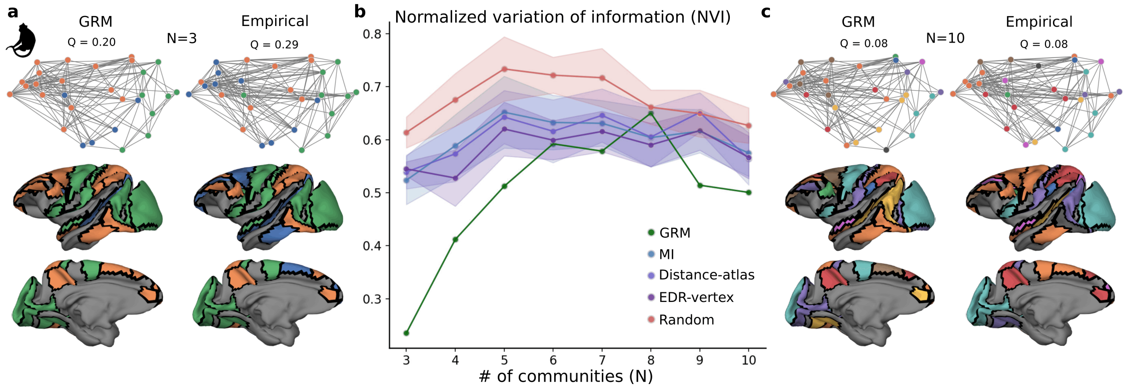


**Figure S33**: **The GEM captures the multiscale topography of the empirical atlas-based macaque connectome's community structure better than benchmark models.** Same as Fig. S31, but for the macaque sparse connectome. The AUC for each model is: GEM = 3.63, MI = 4.27, Distance-atlas = 4.29, EDR-vertex = 4.13, and Random = 4.78.

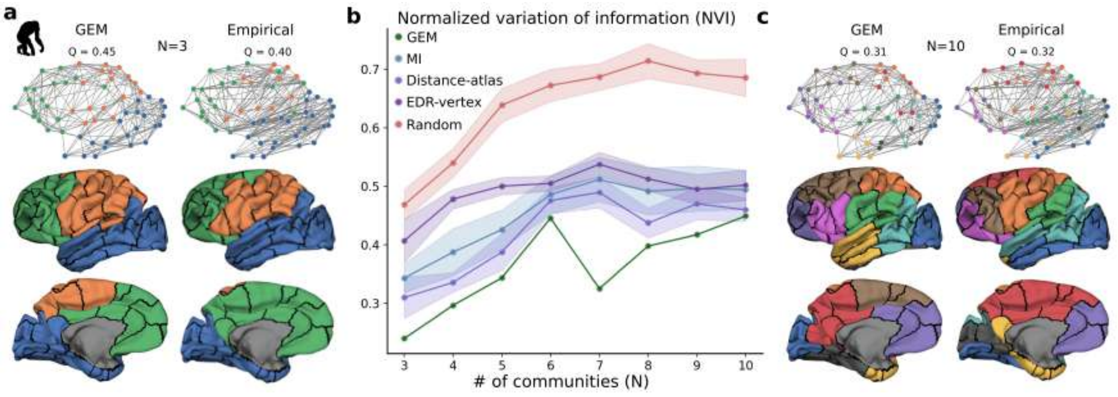


**Figure S34**: **The GEM captures the multiscale topography of the empirical atlas-based chimpanzee connectome's community structure better than benchmark models.** Same as Fig. S31, but for the chimpanzee connectome. The AUC for each model is: GEM = 2.57, MI = 3.22, Distance-atlas = 2.98, EDR-vertex = 3.48, and Random = 4.52.

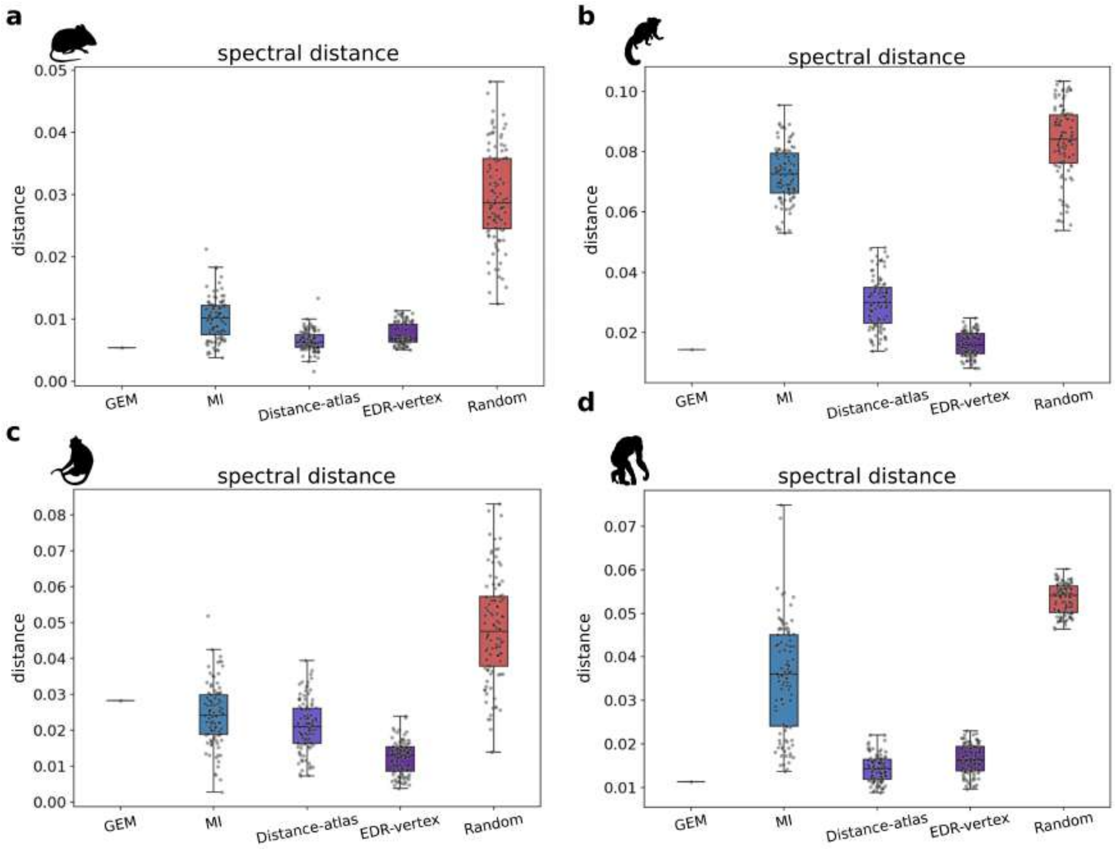


**Figure S35**: **Spectral distances between models and data for the non-human mammalian connectomes.** We assessed the topological similarity between the GEM and the atlas-based connectomes for the four mammalian species using their spectral distance, with lower values indicating better agreement; see section 0.1.9 for details. The four panels show results for the **a:** mouse, **b:** marmoset, **c:** macaque, and **d:** chimpanzee connectomes.

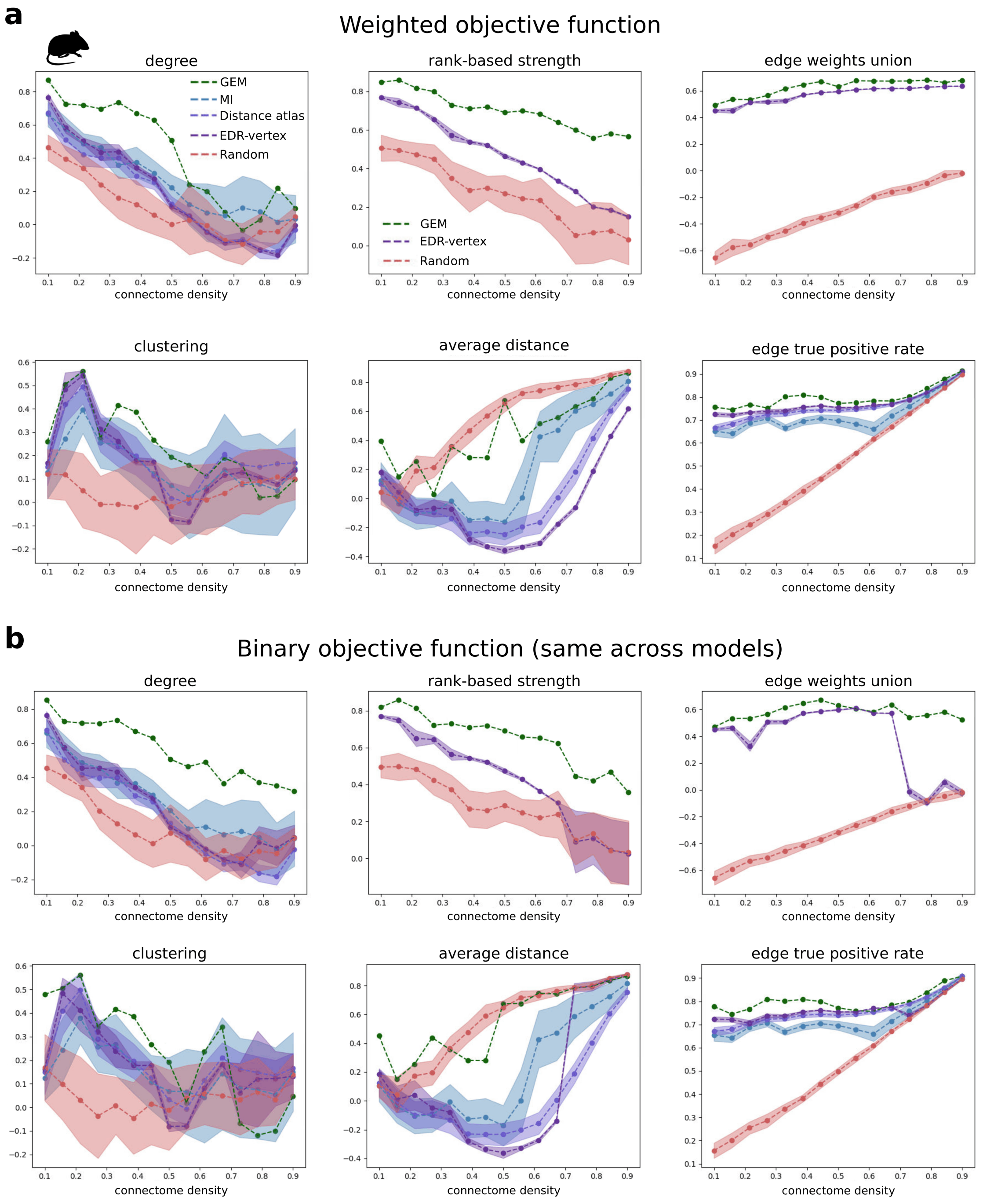
a
Weighted objective function
degree
rank-based strength
edge weights union
clustering
average distance
edge true positive rate
GEM
MI
Distance atlas
EDR-vertex
Random
connectome density
b
Binary objective function (same across models)
degree
rank-based strength
edge weights union
clustering
average distance
edge true positive rate
connectome density

**Figure S36**: **Performance comparison between the GEM and benchmark models across connection densities of the empirical mouse connectome.** In Fig. S24, we evaluated the GEM's performance relative to other benchmarks on the sparse mouse connectome. Here we assessed model performances across 15 empirical connectome densities equally spaced between 10 and 90%, where 90% corresponds to the density of the dense connectome analyzed in Fig. S29; see section 0.1.2 for details. When modeling the sparse connectome in Fig. S24, we used an edge density threshold on the high-resolution models (i.e., GEM, EDR-vertex, and Random) of 20%, and we used 60% when modeling the dense connectome in Fig. S29; see section 0.1.7. In this analysis, we kept a constant threshold value of 60% for the high-resolution models across all empirical connectome densities. The shaded areas represent the standard deviation on the probabilistic models (all the models except the GEM). **a:** GEM performance relative to benchmark models. The GEM and EDR-vertex were optimized using the objective function in Equation S31 for empirical connectome edge densities smaller than 50%, and were optimized with the weighted-only objective function in Equation S32 for higher densities; see section 0.1.8 The MI and Distance-atlas models were optimized using Equation S33 across all edge densities. **b:** Same as panel a, but all models were optimized using Equation S33 across all edge densities.

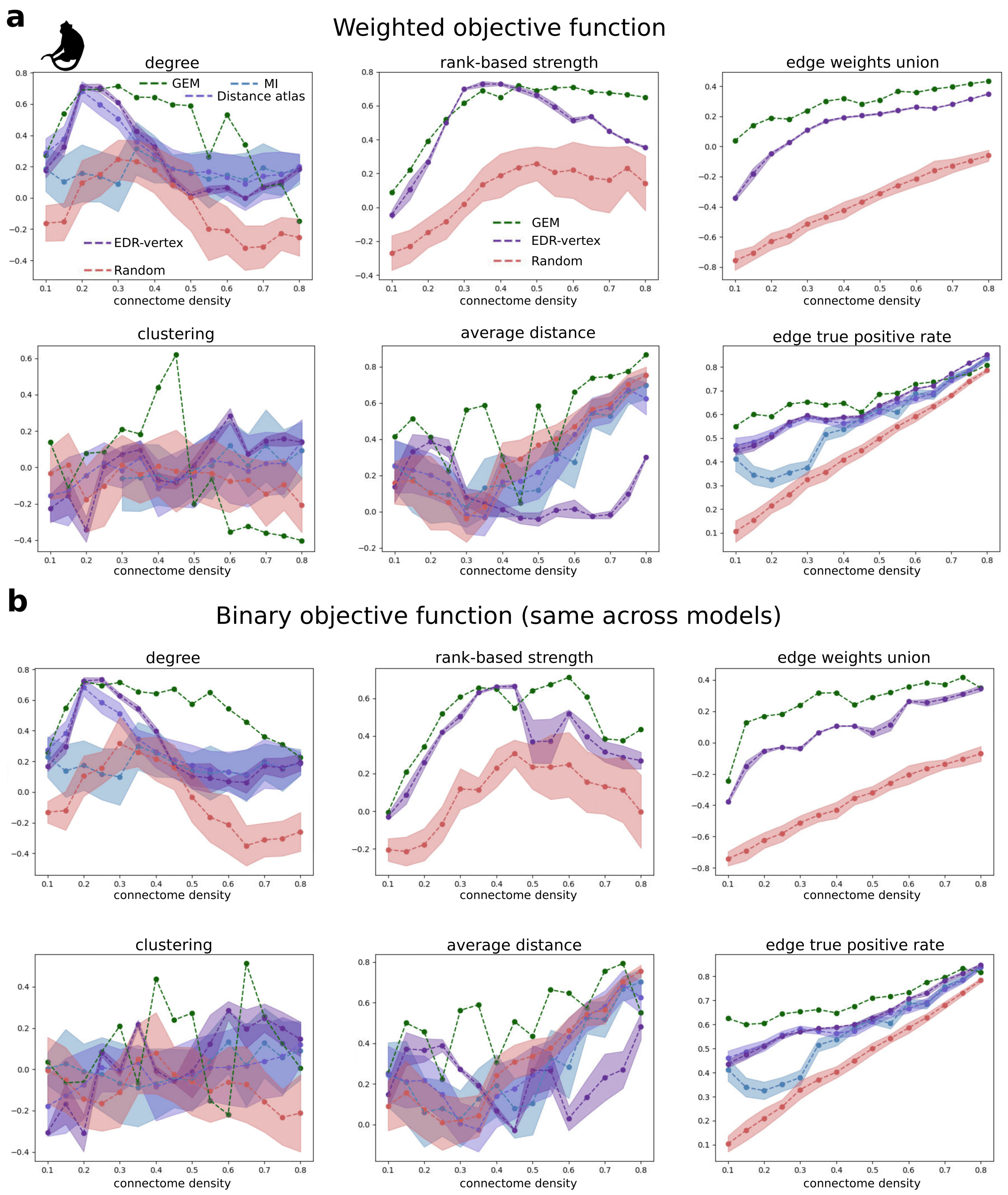


**Figure S37**: **Performance comparison between the GEM and benchmark models across connection densities of the empirical macaque connectome.** Same as Fig. S36, but for the macaque connectome, where we evaluated model performances for 15 equally spaced connectome densities ranging between 10 and 80%, where 80% corresponds to the edge density of the dense connectome analyzed in Fig. S30 and 40% corresponds to the sparse connectome analyzed in Fig. S26.

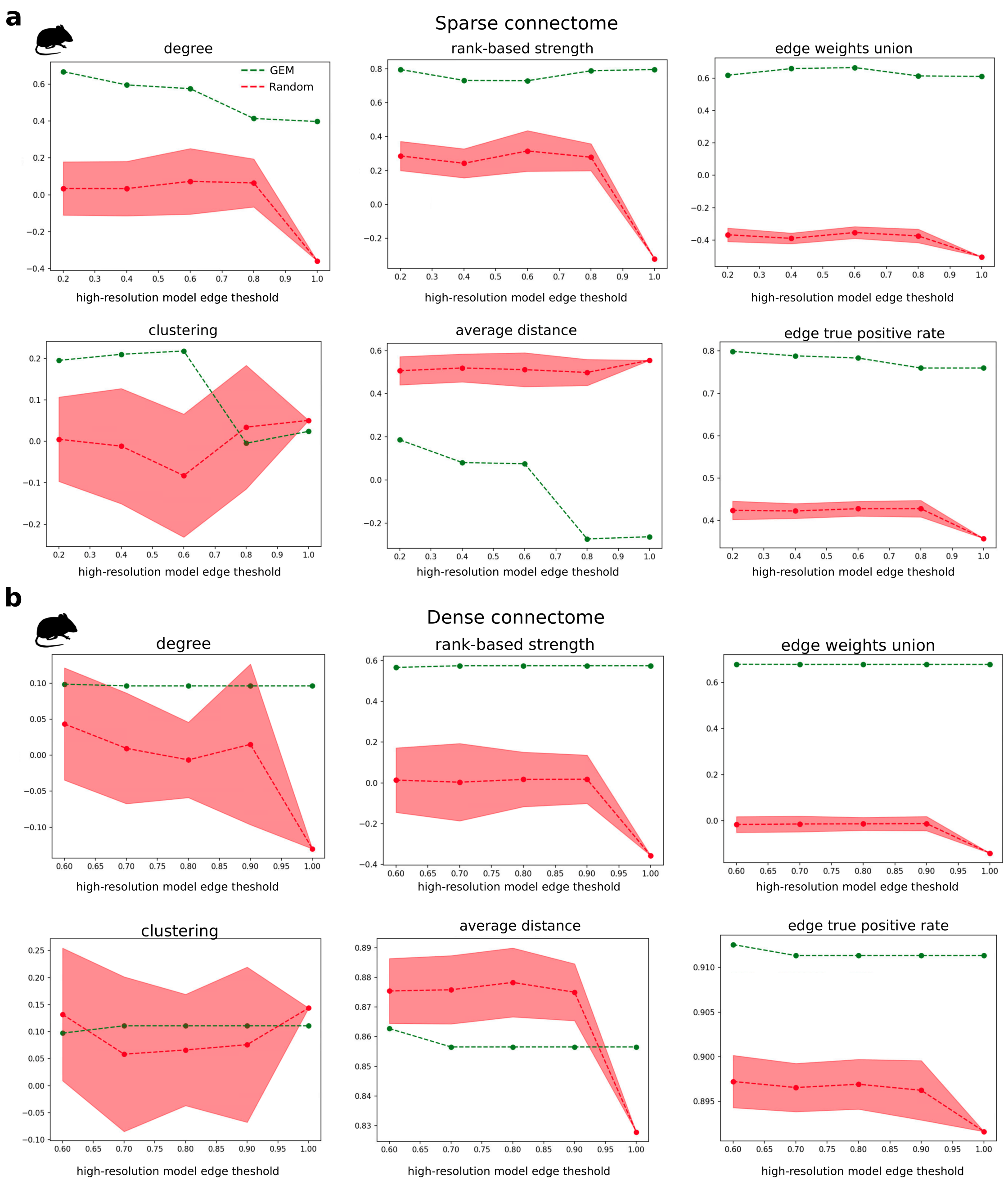
a
Sparse connectome
degree
rank-based strength
edge weights union
GEM
Random
clustering
average distance
edge true positive rate
high-resolution model edge theshold
b
Dense connectome
degree
rank-based strength
edge weights union
clustering
average distance
edge true positive rate
high-resolution model edge theshold

**Figure S38**: **Effect of the high-resolution model edge density on GEM performance in the mouse connectome.** When analyzing the sparse mouse connectome in Fig. S24, we used a high-resolution model edge density threshold of 20% and we used a threshold of 60% in Fig. S29 when analyzing the dense connectome. Here we vary this parameter when analyzing both connectomes, and compare with the Random benchmark, which quantifies the parcellation bias associated with coarse-graining a high-resolution model; see section 0.1.6 and 0.1.7 for more details. **a:** GEM and Random model performances on the sparse mouse connectome across edge density thresholds between 20 and 100%, where 100% corresponds to no threshold being applied. **b:** Same as panel a, but comparing the performance on the dense connectome, with threshold values between 60 and 100%.

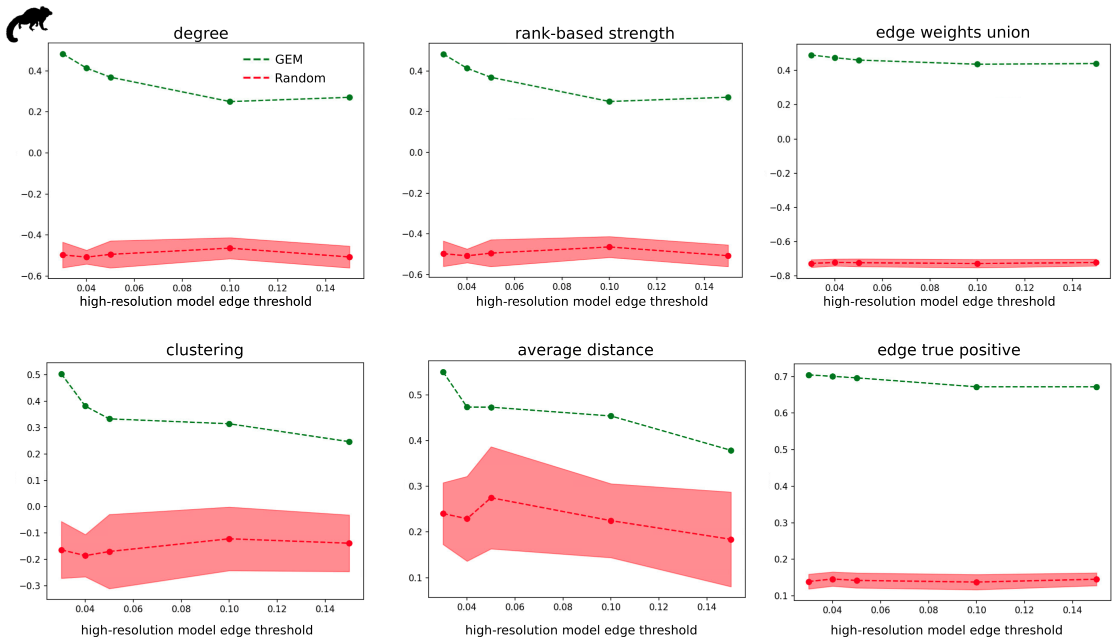


**Figure S39**: **Effect of the high-resolution model edge density on GEM performance in the marmoset connectome.** Same as Fig. S38, but for the marmosets connectome. We evaluated performances for values of the high-resolution model edge density ranging between 3 and 15%.

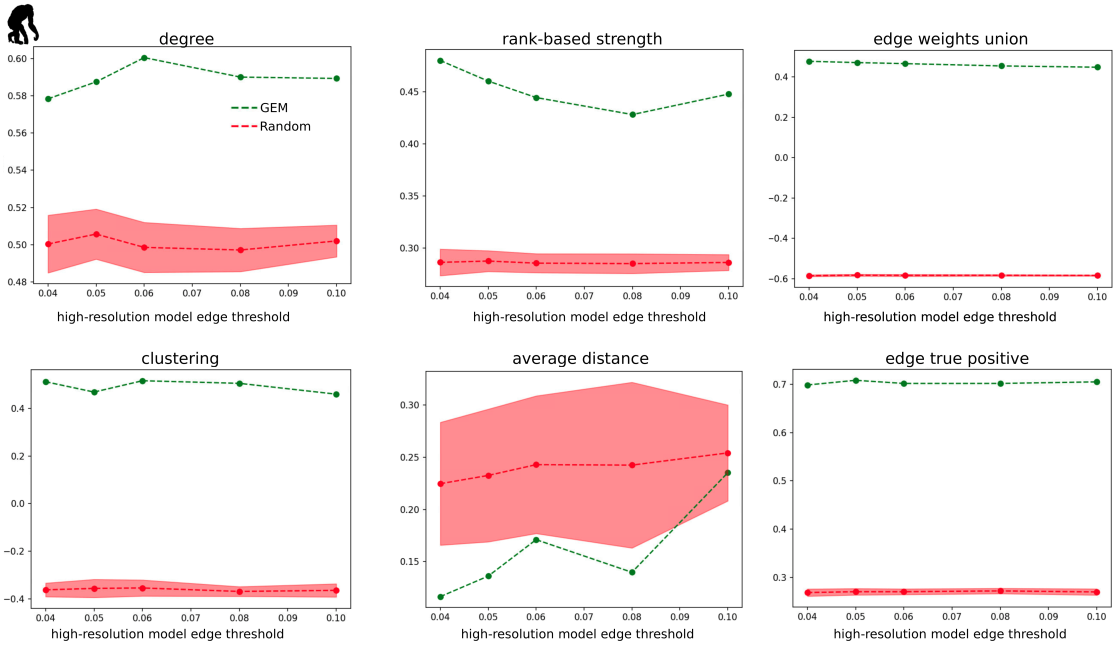


**Figure S40**: **Effect of the high-resolution model edge density on GEM performance in the chimpanzee connectome.** Same as Fig. S38, but for the chimpanzee connectome. We evaluated performances for values of the high-resolution model edge density ranging between 4 and 10%.

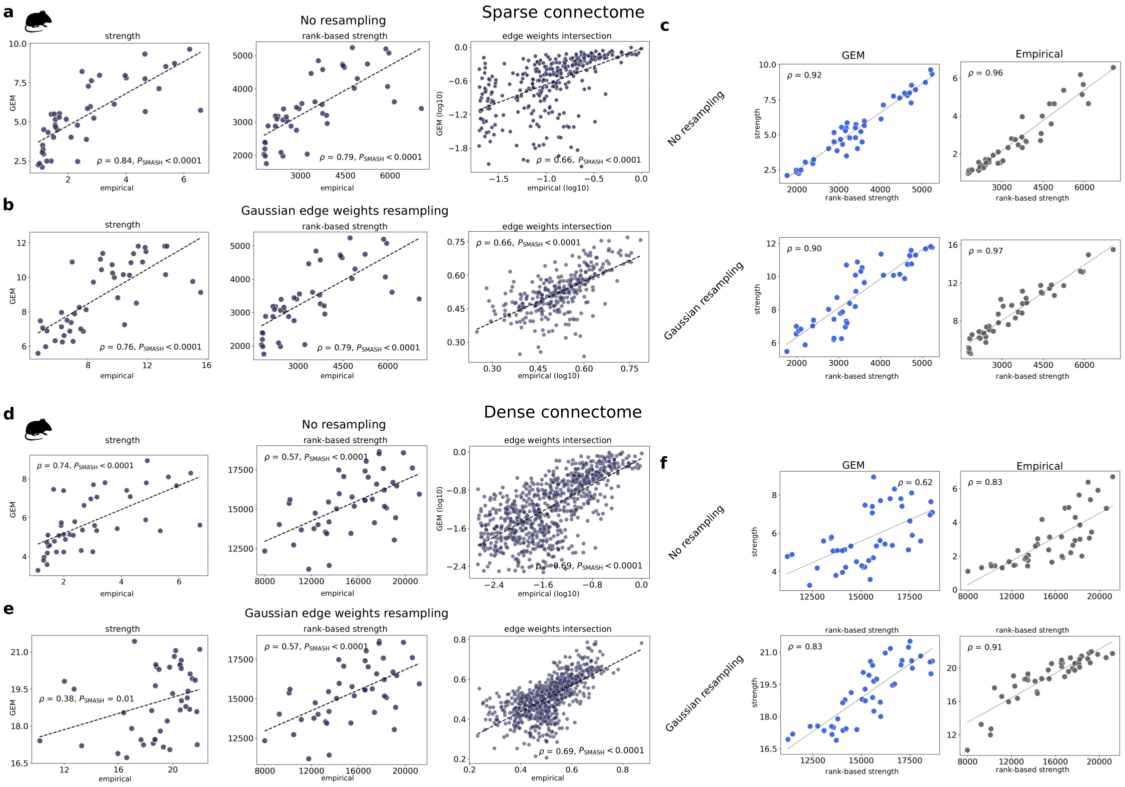


**Figure S41**: **Agreement between different definitions of nodal strengths and effect of edge weights resampling in the atlas-based mouse connectome.** Same as Fig. S23, but for the mouse connectome. Panels **a**, **b**, and **c** show the results on the sparse connectome and panels **d**, **e**, and **f** show the results on the dense connectome.

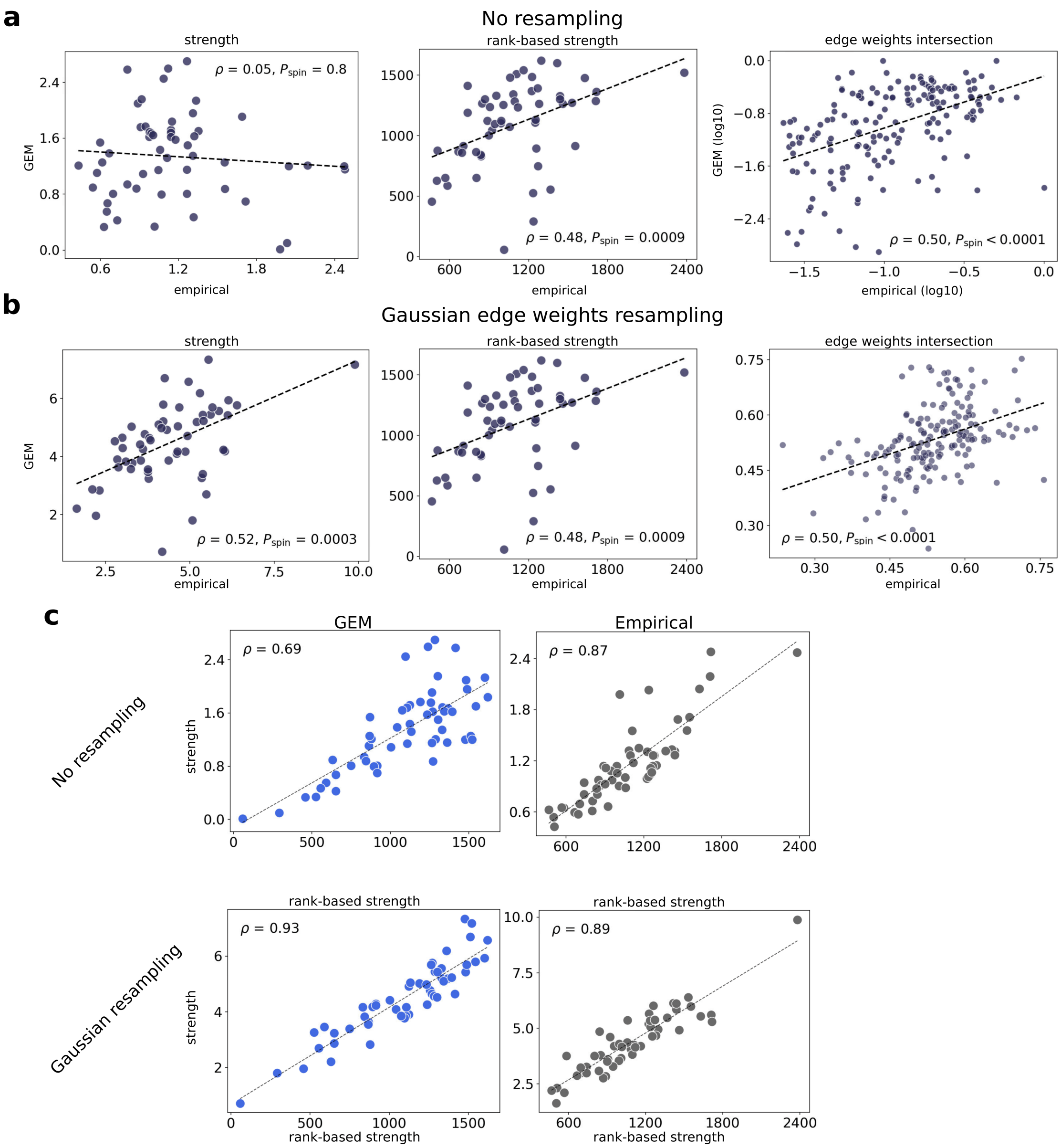


**Figure S42**: **Agreement between different definitions of nodal strengths and effect of edge weights resampling in the atlas-based macaque connectome.** Same as Fig. S41, but for the marmoset connectome.

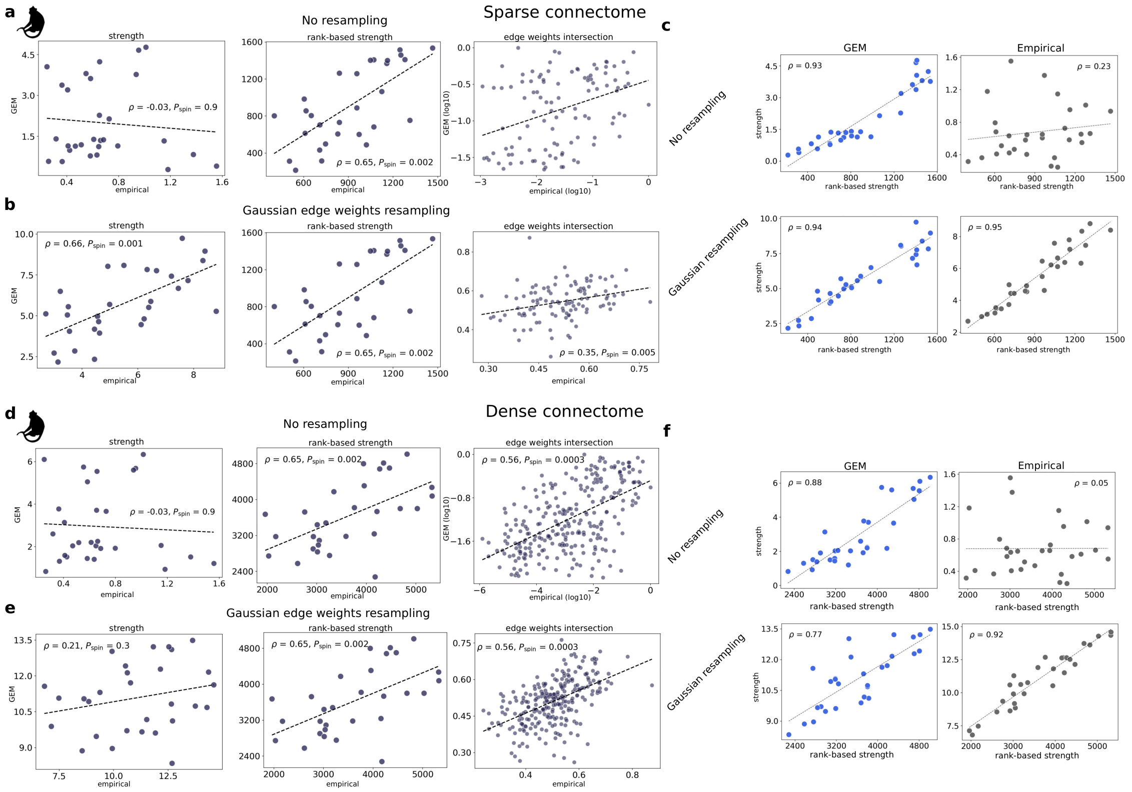


**Figure S43**: **Agreement between different definitions of nodal strengths and effect of edge weights resampling in the atlas-based macaque connectome.** Same as Fig. S41, but for the macaque connectome.

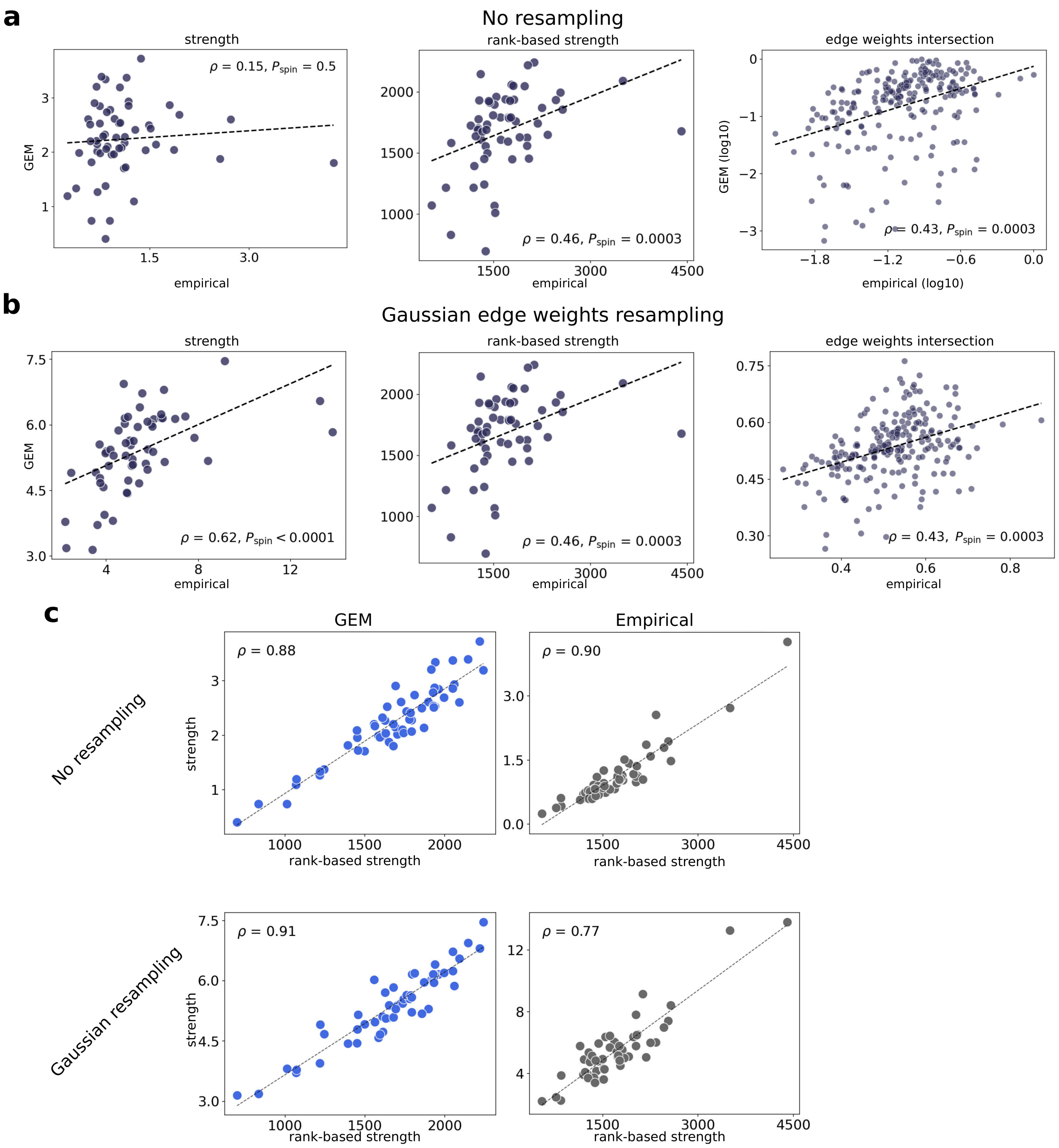


**Figure S44**: **Agreement between different definitions of nodal strengths and effect of edge weights resampling in the atlas-based chimpanzee connectome.** Same as Fig. S42, but for the chimpanzee connectome.

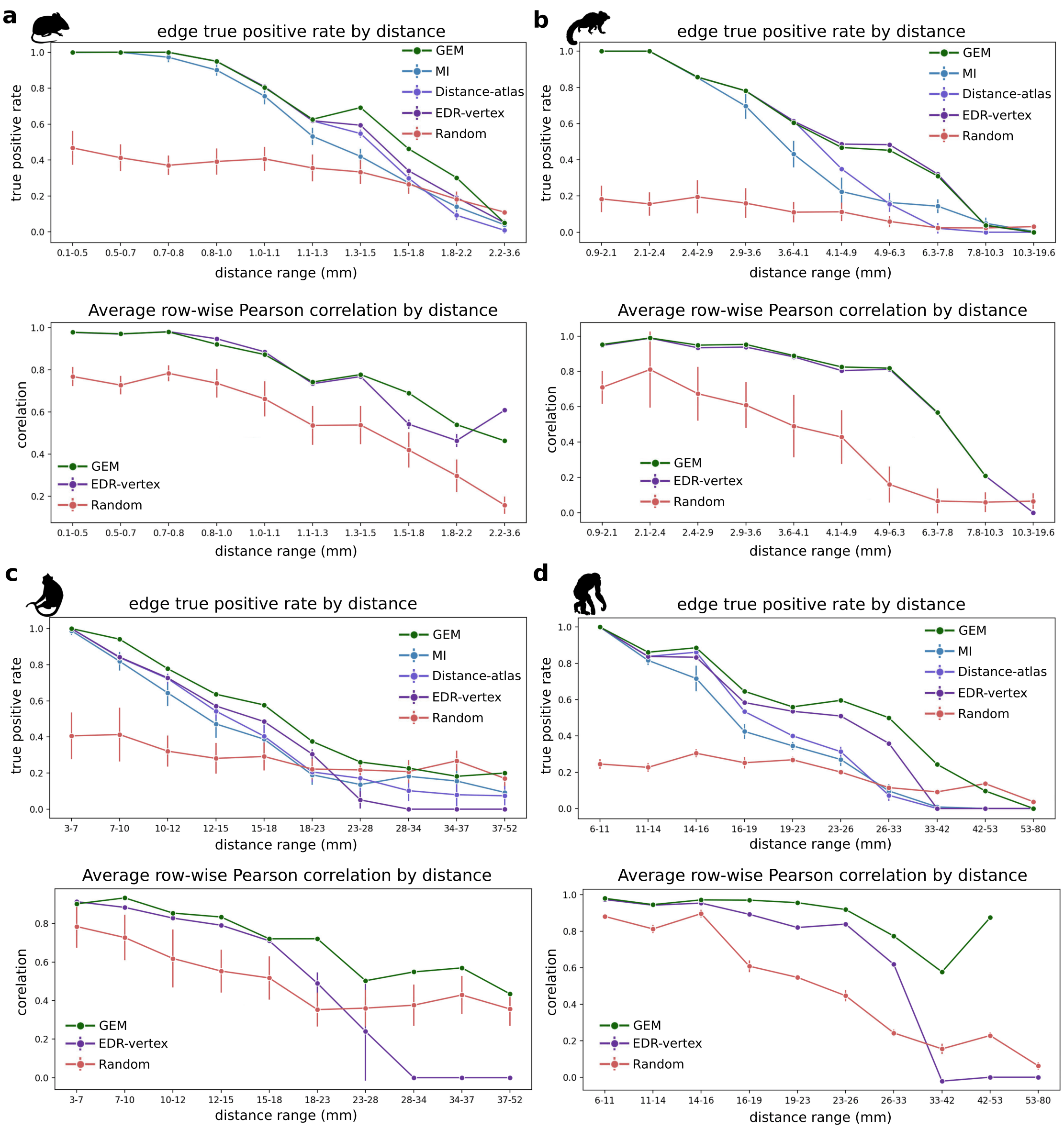


**Figure S45**: **The GEM better captures empirical edges across connection lengths in non-human mammalian species.** For each species, we assessed the GEM's ability to predict empirical edges across 10 connection-length ranges, each containing an equal number of edges; see section 0.1.9 for details. For the mouse and macaque, we used their sparse connectomes to compare with binary atlas-based benchmarks. We found that the GEM generally outperforms or matches the prediction accuracy of other benchmarks across length ranges. **a:** Edge-wise true positive rate (binary) in the mouse sparse connectome as a function of connection length for the GEM, MI, Distance-atlas, EDR-vertex, and Random models (top). Average row-wise Pearson correlation across connection lengths for the GEM, EDR-vertex, and Random models (bottom). **b:**, **c:**, and **d:** show the same analysis, but for the marmoset, macaque, and chimpanzee connectomes, respectively.
